# Two-photon driven photoprotection mechanism in echinenone-functionalized Orange Carotenoid Protein

**DOI:** 10.1101/2024.09.06.611699

**Authors:** Stanisław Niziński, Elisabeth Hartmann, Robert L. Shoeman, Adjélé Wilson, Jochen Reinstein, Diana Kirilovsky, Michel Sliwa, Gotard Burdziński, Ilme Schlichting

## Abstract

Orange carotenoid protein (OCP) is a photoactive protein that mediates photoprotection in cyanobacteria. OCP binds different ketocarotenoid chromophores such as echinenone (ECN), 3’- hydroxyechinenone (hECN) and canthaxanthin (CAN). In the dark, OCP is in an inactive orange form known as OCP^O^; upon illumination, a red active state is formed, referred to as OCP^R^, that can interact with the phycobilisome. Large gaps still exist in the mechanistic understanding of the events between photon absorption and formation of the OCP^R^ state. Recent studies suggested that more than one photon may be absorbed during the photocycle. Using a two-pulse excitation setup with variable time delays we demonstrate that canthaxanthin-functionalized OCP^O^ forms the OCP^R^ signature after absorption of a single photon. By contrast, OCP^O^ complexed with hECN or ECN does not photoconvert to OCP^R^ upon single photon absorption. Instead, OCP^R^ is formed only upon absorption of a second photon, arriving roughly one second after the first one, implying the existence of a metastable light-sensitive OCP^1hv^ intermediate. To the best of our knowledge, a sequential 2-photon absorption mechanism in a single biological photoreceptor chromophore is unique. It results in a non-linear response function with respect to light intensity, effectively generating a threshold switch. In the case of OCP, this prevents down regulation of photosynthesis at low light irradiance.

## Introduction

Sunlight is essential for photosynthesis, but high light intensity can result in photo damage. In cyanobacteria, Orange Carotenoid Protein (OCP) plays a crucial role in a photoprotective process referred to as non-photochemical quenching (NPQ)^1–8^. Under strong illumination conditions, OCP interacts with the antenna complex, the phycobilisome, thereby reducing the energy transfer to the photosystems and thus minimizing the accumulation of generated reactive singlet oxygen radicals. While this process protects the photosynthetic apparatus, it also reduces photosynthesis yield. Thus, for cyanobacteria to thrive in changing light conditions, the interaction between OCP and the phycobilisome must be tightly regulated as a function of light intensity. The regulation is rooted in OCP’s intrinsic light-sensitivity. In the dark or at low light intensity, the protein exists as an orange form, OCP^O^, with low affinity to the phycobilisome.

Under strong illumination conditions, OCP^O^ switches to its metastable active red form, OCP^R^, that binds to the phycobilisome, initiating NPQ (Figure 1). The lifetime of OCP^R^ is usually significantly longer than one minute^3^. In order to fulfil its biological function, the yield of the photoproduct OCP^R^ must depend strongly on light intensity.

**Figure 1.**
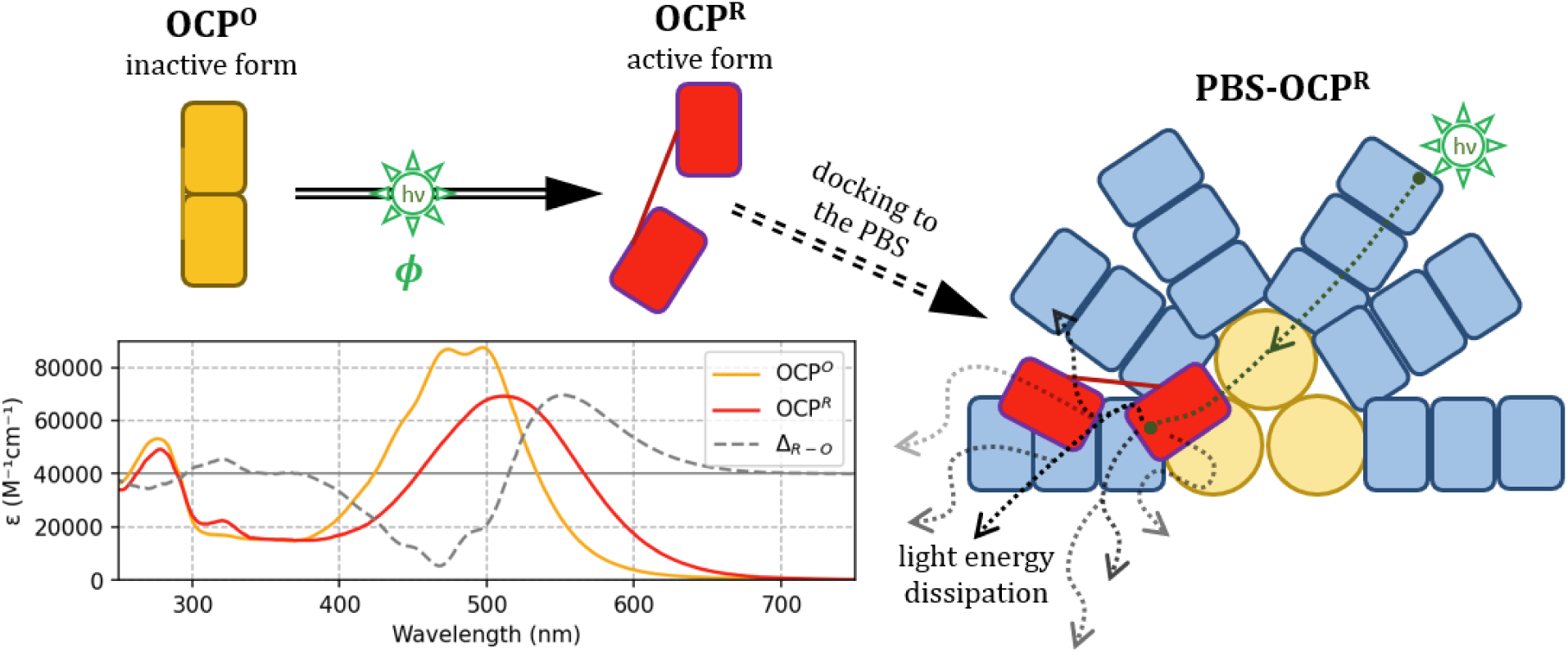
Properties of OCP. Upon illumination with blue-green light, the dark orange form of OCP (OCP^O^), which can be monomeric (shown) or dimeric, transforms into the biologically active monomeric red form (OCP^R^) that can bind to the phycobilisome (PBS, the antenna complex of cyanobacteria), inducing non-photochemical quenching (NPQ). OCP can bind different ketocarotenoid chromophores (omitted). Shown are UV−vis stationary spectra of echinenone-functionalized OCP^O^ from *Synechocystis* (recorded in the dark, with the sample kept at 11 ℃) and after full photoconversion to OCP^R^ (452 nm irradiation of 3.2 mW/cm^2^ until full red form saturation was reached). The absorption is scaled to the molar absorption coefficient reported by Maksimov *et al.* 2020^13^.

The photocycle of OCP is initiated by absorption of a green-blue photon, resulting in formation of a number of electronic ground-state intermediates (P1, P2, P3, PN, PM, PX …) on the ps to µs time-scale^3,9–28^. Most of the studies performed with visible and infrared transient absorption spectroscopy focused on the early events of the reaction. However, it was not verified whether the observed dynamics do indeed result in formation of OCP^R^. Proving this is not trivial, because most time-resolved spectroscopy techniques involve high-repetition rate lasers – restricting the temporal observation window – while OCP^R^ formation occurs on a much longer timescale.

Moreover, the quantum yield of the final OCP^R^ product is extremely low^3^, which renders even normally routine experiments challenging.

Recently, we observed a surprising property of the OCP^O^→OCP^R^ photoconversion process when measuring the speed of accumulation of OCP^R^-like absorbing species upon continuous irradiation^29^. Starting from the dark-adapted state, and analyzing the first time-points of the reaction after switching on the irradiation light, one can calculate a so-called differential quantum yield *ϕ*_*d*_^30^ representing the probability of OCP^R^ formation after photon absorption by OCP^O^. This simplistic approach allows minimizing the number of *a priori* assumptions concerning the photoconversion mechanism. We found that *ϕ*_*d*_ increases with the irradiation intensity, resulting in a non-linear dependency^29^. This is very unexpected, as the quantum yield is a measure of the events per photon absorbed and thus “normalized” with respect to the irradiation intensity. We concluded that we were not observing a reaction that yields the final photoproduct OCP^R^ after absorption of only one photon, but a sequential process, where one of the reaction intermediates absorbs another photon in order to form OCP^R^. In such a case it is much more appropriate to define two (or more) quantum yields describing two (or more) subsequent light-dependent steps. Interestingly, similar conclusions were drawn in a recent report by Rose et al. which showed that OCP photoactivation involves two light-driven reactions mediated by a metastable reaction intermediate^31^. Moreover, Tsoraev et al. reported that repeated photoexcitation within at least 30 s results in a significant increase of the quantum yield of the photoproduct OCP^R32^.

OCP can bind different carotenoid chromophores; 3’-hydroxyechinone (hECN) is the natural chromophore of OCP in *Synechocystis*^1,33^. Due to difficulties of generating enough hECN when overexpressing OCP (even in native strains), the community has studied predominantly OCPs complexed with echinenone (ECN) and canthaxanthin (CAN), respectively, since the proteins involved in their biosynthesis can be overexpressed in *E. coli*. All three chromophores are photoactive and afford OCP^R^ formation^2,3,34,35^. It was shown recently that the photoconversion efficiency of CAN-functionalized OCPs is higher than that of their ECN counterparts^10^, implying a significant influence of the additional carbonyl group in the β2 ionone ring in CAN compared to ECN^17^.

Here we investigate the photoconversion mechanism of OCP on the millisecond to minute time scale, addressing specifically the number of light-sensitive steps occurring along the reaction coordinate. We analyze the time-evolution of photoexcited OCP from *Synechocystis* (Syn) and *Planktothrix* (Plk) cyanobacteria complexed with different carotenoid chromophores (hECN, ECN, CAN). To this end, we use an experimental setup involving one or two submillisecond laser pulses with a variable time delay between the latter. We show that for ECN- and hECN- complexed OCP the spectral signature associated with OCP^R^ formation is only observed when two excitation pulses separated by a time-delay of about 1 second are employed. Therefore, OCP functionalized with ECN/hECN exhibits a two-photon mechanism 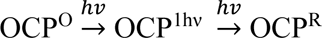 involving a light-sensitive intermediate OCP^1hν^, resulting in a strictly nonlinear irradiation intensity dependent response. By contrast, for CAN complexed OCP, the spectral signature associated with OCP^R^ is observed already when using only a single light pulse.

## Results

### Setup for the detection of a sequential two-photon photoconversion mechanism

The most straightforward approach to investigate the number of photon absorption events required to drive the OCP^O^→OCP^R^ photoconversion is to utilize multiple excitation pulses. Such pump-pump-probe techniques are used to characterize nonlinear photochromic materials^36^ and have demonstrated, for example, the increase of photoproduct formation in experiments applying two femtosecond excitation pulses delayed by few picoseconds^37^. Here we investigated the dynamics of OCP^R^ formation on the millisecond-to-second time-scale using two submillisecond pump pulses and a continuous visible light probe. If the OCP^O^→OCP^R^ photoconversion process is single-photon driven, one excitation pulse should be sufficient to obtain a proper OCP^R^ signature, otherwise another pulse is required. However, observing OCP^R^ is far from trivial, not only because of its low quantum yield (less than a few percent^3,16,25,27,29^, note the reported values are highly inconsistent), but also due to the fact that most time-resolved experiments operate at high repetition rate and do not allow to check whether or not the process initiated by the excitation laser pulse terminates in an expected way on a long timescale (> 100ms). Moreover, it is challenging to ensure that for each repetition of the experiment a fresh aliquot of the sample, free of any unrelaxed intermediates, is probed. This is complicated even more by the requirement that acquisition of seconds-long kinetics requires a non-exchanging (still) sample without diffusion processes mixing an irradiated sample aliquot in the beam focus and the rest of the sample. In order to overcome these challenges, we developed a dedicated visible broadband transient absorption spectroscopy setup for pump-pump-probe experiments probing in the millisecond-second time regime. The OCP solution was mounted in a quartz cuvette with a 100 µm light path length, which greatly limits diffusive sample exchange due to capillary forces (see Figure 2A). The laser irradiation pulses were focused to a small ≈190 µm spot (FWHM), so that one cuvette could be probed at multiple places. Each kinetic was measured continuously for 20 s with ≈3 ms temporal resolution and then the cuvette was translated. This way, after each measurement, a fresh and completely dark-adapted sample aliquot is probed, with no memory effects of previous pulses.

**Figure 2.**
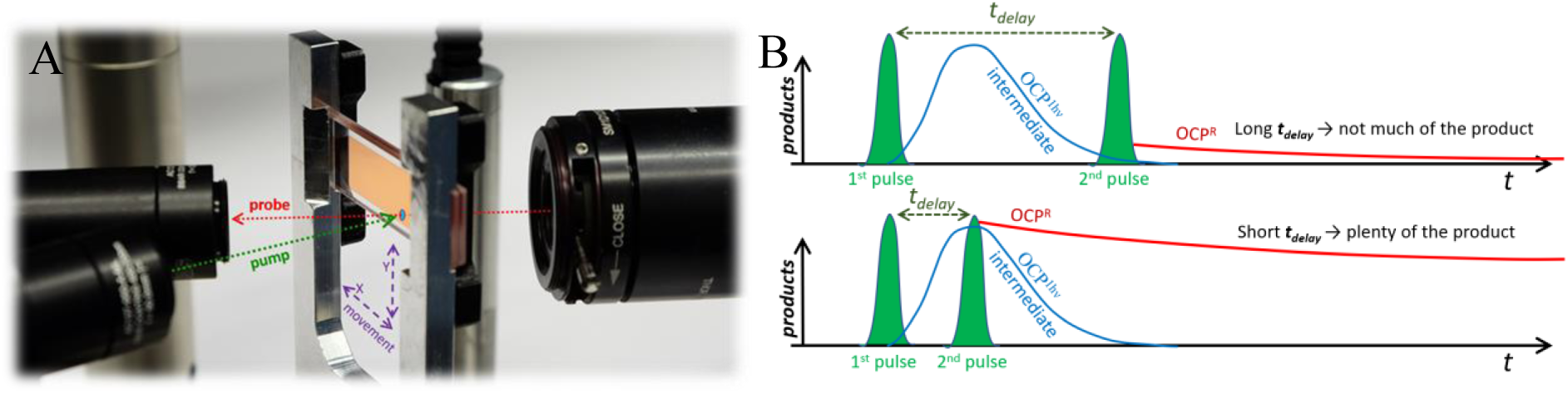
**Dedicated visible broadband transient absorption spectroscopy setup for pump- pump-probe experiments on OCP**. A) Geometry of the experiment. The sample is contained in a flat cell with 100 µm optical pathlength. A translation stage displaces the sample each time after the kinetics has been recorded. B) Experimental concept. In a two-photon mechanism, the first irradiation pulse populates an intermediate photoproduct OCP^1hv^, while the second pulse can photoconvert it to OCP^R^. Its yield is low if the lifetime of the one-photon generated intermediate OCP^1hv^ is much shorter than the time delay ***tdelay*** between the two light pulses (shown in green, upper trace). By contrast, if the peak of the OCP^1hv^ intermediate concentration coincides with the second light pulse, the OCP^R^ yield is high (lower trace). Thus, fine-tuning of t ***tdelay*** is necessary to observe significant formation of OCP^R^.

We recorded transient absorption kinetics observed after a single irradiation pulse (denoted as 1*hv*) and after a pair of pulses that are temporarily separated by *t*_*delay*_ (denoted as 2*hv*), respectively. In both cases, the total absorbance difference Δ*A* is determined with respect to the absorbance recorded without irradiation pulses, *i.e.* Δ*A* represents the cumulative change in absorption caused by a single or two excitation pulses, respectively. As illustrated in Figure 2B, a variable *t*_*delay*_ allows to probe the formation and decay time of an OCP^1hv^ intermediate, should it exist. If the OCP^O^ → OCP^R^ photoconversion is a purely single-photon initialized process, then the 2*hv* kinetics are *t*_*delay*_-independent and simply resemble rescaled 1*hv*-kinetics.

### Kinetics obtained for ECN and hECN OCPs after a single pump pulse and two pump pulses

We used this experimental setup to investigate hECN, ECN and CAN-functionalized OCPs from *Synechocystis* (and *Planktothrix,* see Supplement and below) cyanobacteria. The 6-His affinity tags used for purification were removed proteolytically, to avoid effects of tagging^10,12^. The experimental results are shown in Figure 3, demonstrating how the investigated proteins react to single-pulse and two-pulse excitations. OCP^R^ is characterized by a red-shifted spectrum compared to OCP^O^ (Figure 1). The kinetics probed at 585 nm (Figure 3D) are especially well suited to track the presence of the OCP^R^ product because they take into account mostly absorption of the OCP^R^ form and minimize the contributions from the bleached OCP^O^ population and intermediate forms populated after the first pulse, respectively. For ECN-OCP no measurable positive transient absorption signal at 585 nm is observed after one laser pulse (1hν labeled kinetics). The only spectral feature observed is a pair of negative bands at ≈495 nm and ≈545 nm, of which only the first one lives longer than a few seconds (Figure 3A-C, dashed spectra). A recent report by Chukhutsina *et al.* describes a similar peculiar spectral signature with a dominating negative ΔA contribution occurring on the µs-ms timescale^17^. This feature may indicate a *trans*-*cis* isomerization of the conjugated double bond system of the carotenoid, which is usually associated with a hypochromic shift and change in vibrational structure^38,39^. Importantly, our data clearly show that even after a strong short laser pulse (about 50mJ/cm^2^) the protein cannot photoconvert to species resembling the OCP^R^ protein form, for which a large band at 550 nm is expected. Such an intense laser pulse was used intentionally, because it demonstrates that even multi-round excitation of an OCP molecule does not result in OCP^R^ formation if the re-excitation occurs within a short period of time (below single milliseconds), see also reference^11^. An OCP^R^-like spectral signature appears only after a properly delayed second pulse (Figure 3B, continuous line), with maximal yield for a time-delay of about 1 s. For a 10 ms or 100 s delay, the positive transient absorption band is negligible (Figures 3A and 3C, respectively). *t*_*delay*_ must be roughly between 10 ms and 10 s in order to re-excite OCP^1hν^ and yield OCP^R^. This peculiar timing dependence is apparent in Figure 3E, showing that the OCP^R^ yield depends strongly on *t*_*delay*_, peaking at *t*_*delay*_ = 1s. Experiments performed on hECN- functionalized OCP show very similar kinetics as for ECN-functionalized protein (see Figure S8 in the Supporting Information).

**Figure 3.**
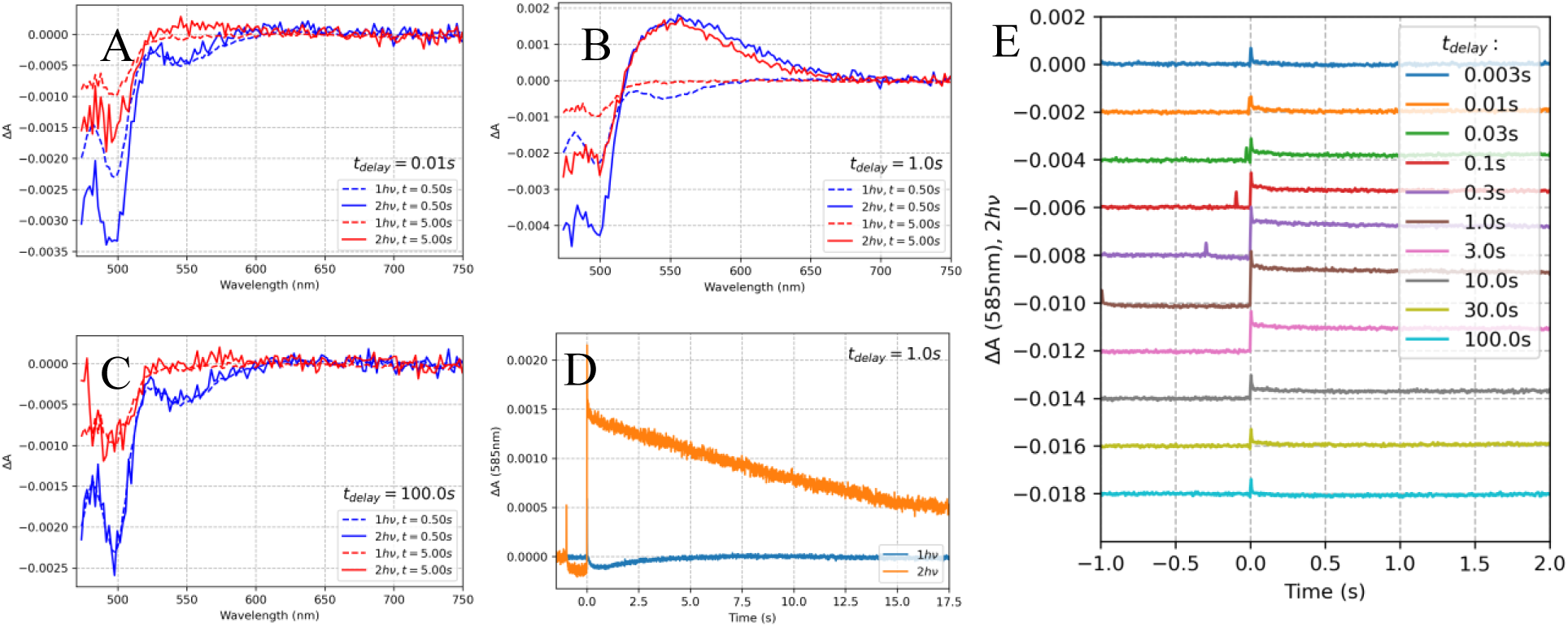
Two-pulse excitation experiment analyzing ECN-functionalized OCP from *Synechocystis* without His-tag. Transient absorption spectra obtained after two 512 nm excitation pulses separated by *t*_*delay*_ of A) 10 ms, B) 1 s, C) 100 s. D) Kinetics at 585 nm measured after excitation with a single pulse (1hν) and a pair of pulses (2hν) separated by 1s. E) Kinetics at 585 nm recorded after two pulses separated by various delays, time zero is set at the 2^nd^ pulse.

We conclude that the ≈495 nm negative ΔA feature represents a reaction intermediate (which we refer to as OCP^1hν^), which turns into OCP^R^ after re-excitation: 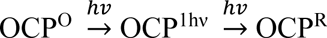. Of note, the *t*_*delay*_ value resulting in rise of this positive OCP^R^-like spectral signature (maximum at 550 nm) must be shorter than the decay of the negative ≈495 nm feature visible after only one excitation pulse (Figure 4). However, if *t*_*delay*_ value is very short (<10 ms), no OCP^R^ signature is observed either. This suggests that the OCP^1hν^ intermediate is formed after the decay of an earlier intermediate form that is characterized by a pair of negative transient absorption bands at ≈495 nm and ≈545 nm. Note that OCP^1hν^ is characterized solely by the ≈495 nm negative ΔA band.

**Figure 4.**
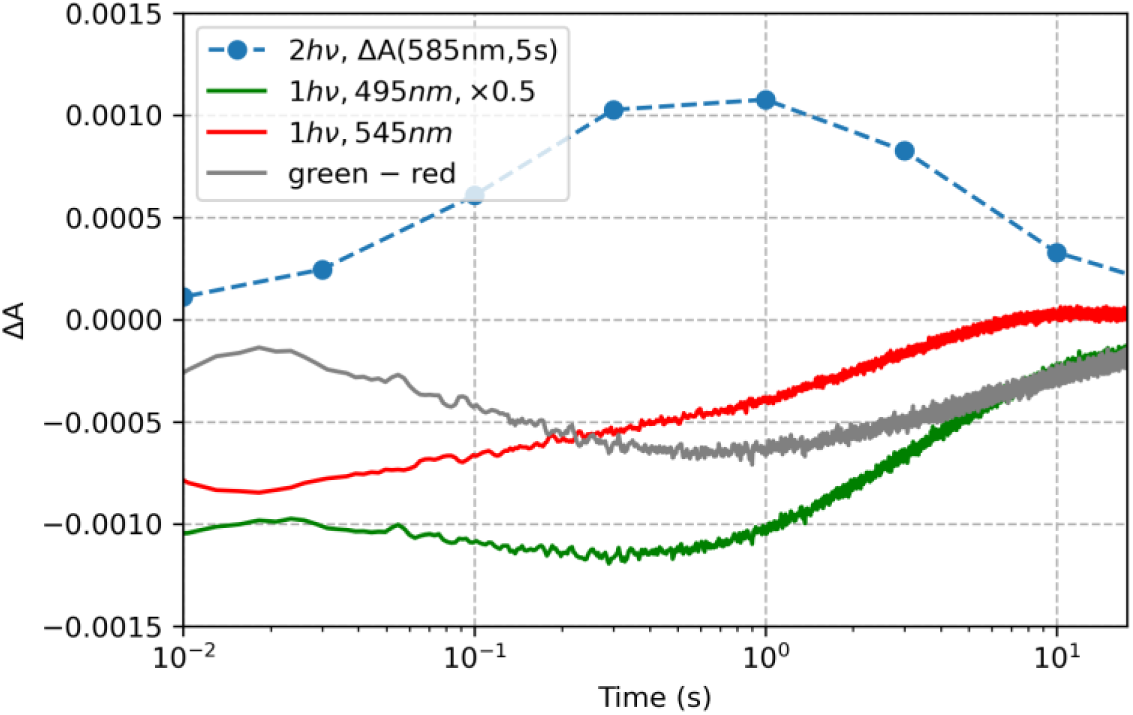
Temporal evolution of spectroscopic signatures. The green and red curves show the transient absorption kinetics probed at 495nm and 545 nm, respectively, obtained after pumping with only one pulse, plotted as a function of time (scaled by a factor of 0.5 and 1, respectively). Only the 495 nm signal lives long enough to be considered as OCP^1hν^. This suggests that the OCP^1hν^ intermediate is associated exclusively with the negative ΔA peak at 495 nm, while an earlier intermediate form is associated with both negative ΔA peaks at 495 nm and 545 nm, respectively. If so, one should be able to obtain OCP^1hν^ concentration profile by subtracting 545 nm kinetics from the 495 nm kinetics (with proper weights). Indeed, the difference between the green (495 nm) and red (545 nm) curves plotted as the grey curve, mirrors the temporal dependence of the blue curve, which represents the OCP^R^ product yield (ΔA observed at 585nm, 5s after the second of two excitation pulses) plotted as a function of *t*_*delay*_. The data show ECN-functionalized OCP from *Synechocystis* (50mJ/cm^2^ pump).

### Kinetics obtained for CAN-OCPs after single and double pump pulses

CAN-functionalized OCP reacts very differently than ECN-complexed OCP (see Figure 5). Already after one excitation pulse, a long-living transient absorption signal ΔA appears (see Figure 5D) that resembles the OCP^R^ spectral signature (Figure 5A-C, dashed spectra). In order to obtain signals of comparable magnitude as for ECN-OCP, a much weaker pump pulse of 3 mJ/cm^2^ (Figure 5) is sufficient (see Supplementary Information for the results of a laser energy titration). The growth kinetics of the 585 nm absorbance resembles a Heaviside (step) function (Figure 5E). After a second pulse, another ΔA step is observed that has the same height as the first one and decays equally fast (Figure 5D, 1hν label and Figure S18A,D in the Supporting Information). The spectra obtained after two pulses with *t*_*delay*_ values of 10 ms, 1 s and 10 s do not show significant variability (Figures 5A-C). In contrast to ECN-OCP, the 585 nm transient absorption signal is invariant of *t*_*delay*_ for CAN-OCP (Figure 5E). In conclusion, in CAN-OCP the OCP^R^ form is populated after only one laser pulse; use of two pulses leads to an additive effect (see the comparison in Supporting Information, Figure S18A, D). These results explain not only the higher photoconversion efficiency of CAN-functionalized OCPs compared to their ECN counterparts observed previously^10^, but also the observation that upon photoexcitation with a ns pulse CAN-OCP^O^ converts to OCP^R^ whereas ECN-OCP^O^ does not^10^. We note that for higher pump pulse energies (see Figures S7 and S16 in Supporting Information) the second pulse leads to a faster decay and slightly broader spectral signature with an additional 550-630 nm flank (see also the direct comparison in Figure S18 in the Supporting Information), similar to observations by Rose *et al.*^31^ and by Wilson *et al.*^10^ (Fig. 2B in reference^10^) upon a longer period of continuous irradiation. Since these effects are absent after weaker irradiation pulses (3 mJ/cm^2^), they must have a different origin than a two-photon photoconversion process.

**Figure 5.**
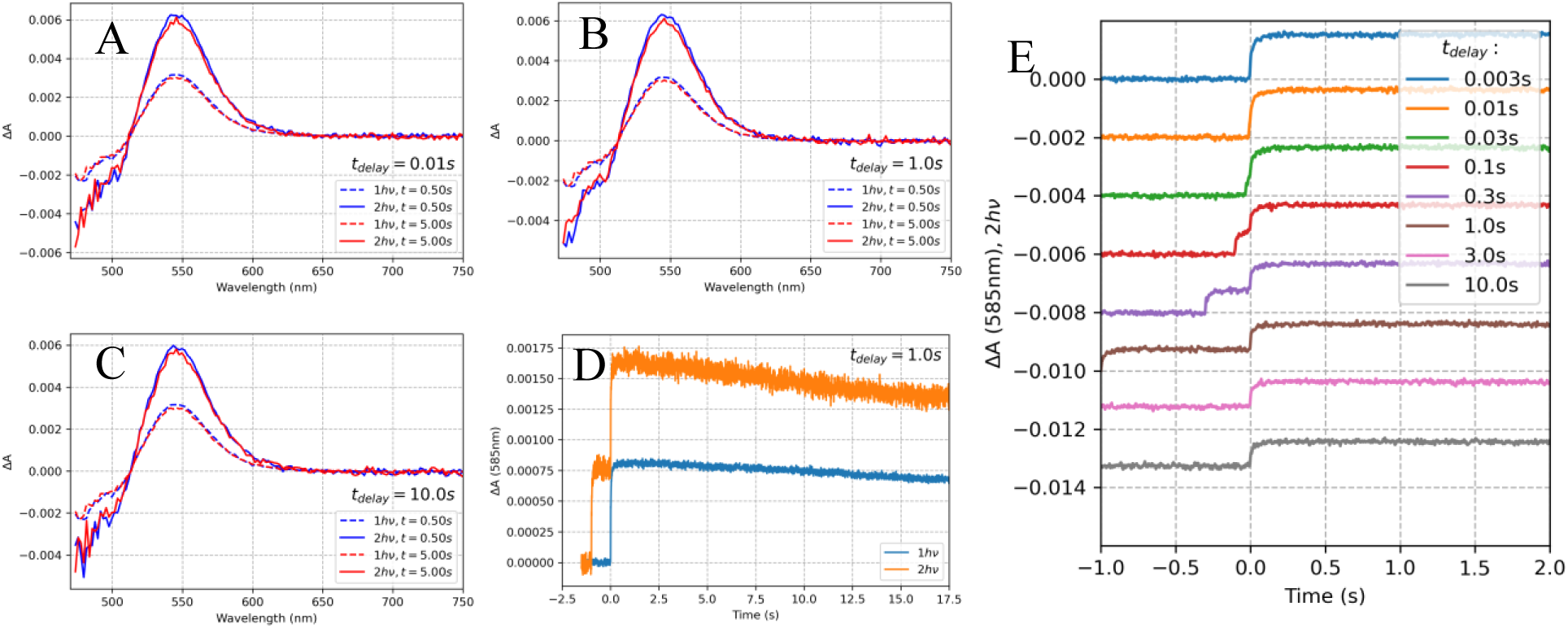
Two-pulse excitation experiment analyzing CAN-functionalized OCP from *Synechocystis* without His-tag. Transient absorption spectra obtained after two 512 nm excitation pulses (3mJ/cm^2^) spaced by *t*_*delay*_ of A) 10 ms, B) 1 s, C) 10 s. D) Kinetics at 585 nm measured after excitation with a single pulse (1hν) and a pair of pulses (2hν) separated by 1s. E) Kinetics at 585 nm recorded after two pulses separated by various delays, time zero is set at the 2^nd^ pulse.

### Chromophore type determines 1hν or 2hν driven photoconversion mechanism

We tested whether the difference in reaction mechanism observed for CAN- and ECN-functionalized OCP from *Synechocystis* also applies to OCP from *Planktothrix* (see Supplementary Figures S6 and S10). The dependence of the transient absorption signal at 585 nm associated with an OCP^R^-like product on the time delay *t*_*delay*_ between the two excitation pulses is shown in Figure 6. A strong maximum at *t*_*delay*_=1 s is observed for ECN-functionalized proteins from both cyanobacterial strains, implying that the OCP^1hν^ population is peaking. By contrast, the transient absorption signal at 585 nm does not depend on *t*_*delay*_ in CAN-functionalized proteins from both strains. For higher excitation pulse energies, there is a minor dependence of the OCP^R^ signal of *t*_*delay*_ (see Figure S19 in the Supporting Information).

**Figure 6.**
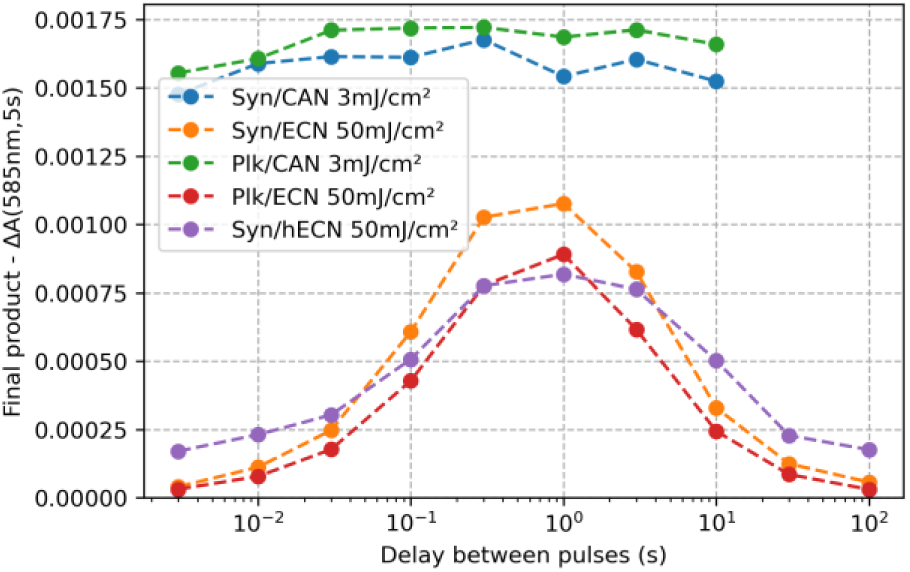
**Comparison of CAN- and ECN-functionalized OCP from *Synechocystis* and *Planktothrix.*** The change in absorption at 585 nm recorded 5 s after the second pulse, representing the yield of OCP^R^, is plotted against the time delay *tdelay* between the two laser excitation pulses. It is striking that the photoresponse of CAN-functionalized OCPs differs strongly from ECN and hECN complexed OCPs. Stationary spectra of the samples used are shown in Figure S4B in the Supporting Information.

However, its absence in the spectra obtained after the weakest excitation pulse (3 mJ/cm^2^) excludes the possibility that it is a manifestation of a two-photon mechanism.

Since most studies on OCP are performed using His-tagged variants we also tested whether these proteins behave analogously to the un-tagged proteins. Indeed, ECN-OCPs from *Synechocystis*, His-tagged at the N- and C- termini, respectively, photoconvert according to the two-photon mechanism (see Figures S20 and S21 in supporting information, respectively). However, we note that the spectral signatures of their OCP^1hν^ intermediates show additional positive contributions compared to the tag-free variant, resembling OCP^R^ more. In conclusion, the photoconversion mechanism is preserved among different OCP variants and depends only on the bound chromophore.

### Comparison with kinetics obtained upon continuous light irradiation

We wondered how these results correlate with kinetic measurements performed using continuous light irradiation of OCP. Previously we had proposed the need for a 2-photon mechanism of the 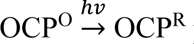 photoconversion^29^ based on the intensity dependence of the differential quantum yield *ϕ*_*d*_ of ECN-complexed OCP from *Planktothrix*^30^. Here, we plot *ϕ*_*d*_ versus irradiation photon flux density for CAN-functionalized OCPs (Figure 7A). The *ϕ*_*d*_ value is constant, except for high irradiation intensity where it decreases due to fast saturation of the OCP^R^ population, resulting in an early plateau (as modelled in our previous report^29^). Thus, the true quantum yield for CAN-OCPs is invariant of the irradiation intensity, allowing to describe the 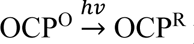 photoconversion by a single, well-defined quantum yield. By contrast, for ECN-functionalized OCP *ϕ*_*d*_ is negligible for weak continuous irradiation intensity, whereas it grows significantly for stronger continuous irradiation (Figure 7A). Strikingly, at low irradiation intensity no photoactivity is observed for ECN-OCPs compared to significant OCP^R^ accumulation for CAN- OCPs (Figure 7B). The finding that ECN-OCPs are photoactive only above a specific light intensity threshold is in line with the two pulse excitation results.

**Figure 7.**
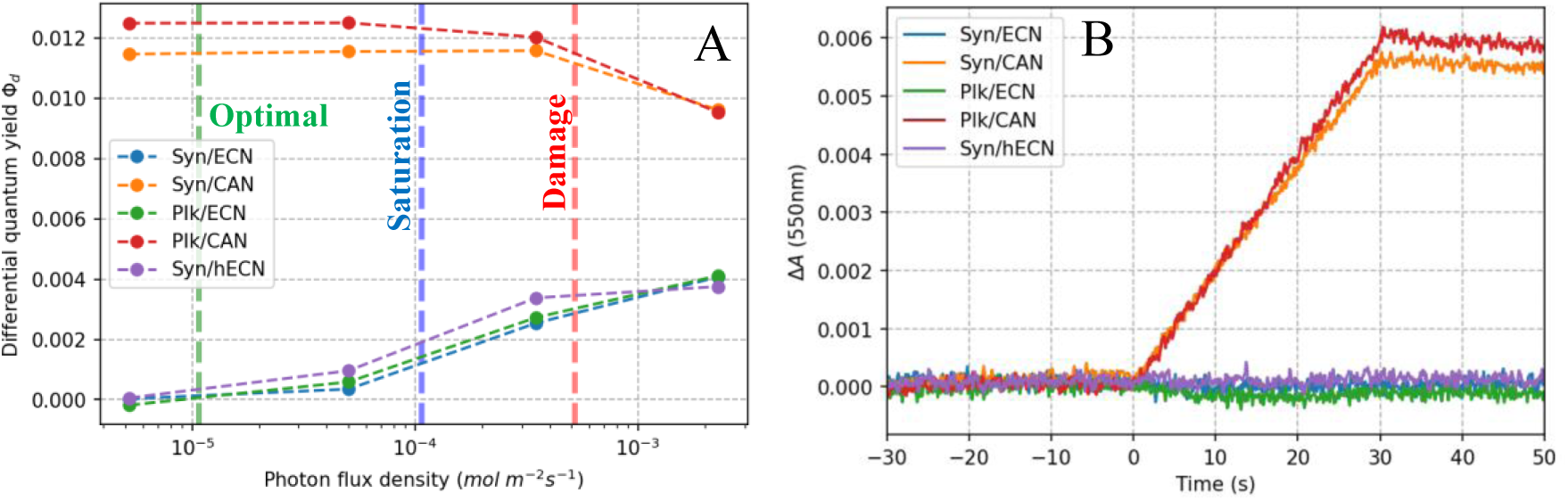
Differential quantum yields of CAN and ECN complexed OCP exposed to stationary irradiation. A) We used the method described previously to derive the differential quantum yield *ϕ*_*d*_^29^ as a function of the photon flux density at 472 nm. Vertical lines mark “optimal”, “saturation” and “damage” photon flux densities that correspond to photosynthetic photon flux densities (PPFD) of 30, 300 and 1460 µmol photons s^-1^ m^-2^ discussed in the context of cyanobacterial photosynthesis. Note, since OCP absorbs only a fraction of the photosynthetically active radiation (PAR) the values were recalculated using a conversion factor of 2.8, more details are in the Methods section. B) Difference absorbance kinetics at 550 nm observed upon weak irradiation using a 472 nm LED (photon flux density 5.2·10^-6^ mol m^-2^ s^-1^). The irradiation light is switched on at t=0 and off at t=30 s, respectively, the sample absorbance is A=0.62 (10 mm pathlength, absorption maximum).

### Investigation of the monomeric R27L mutant

OCP exists as a monomer-dimer equilibrium, with a dissociation constant of ≈14 µM^11,40^. Since our experiments were performed at an OCP concentration of ≈60 µM and thus with mainly dimeric protein, it is conceivable that the two-photon mechanism derived for ECN-functionalized OCP is rooted in the absorption of one photon by each monomer instead of the absorption of two photons by a single monomer and thus chromophore. To distinguish between these possibilities, we investigated the R27L mutant of OCP which had been shown to be monomeric^40^. The CAN- complexed R27L mutant photoconverts according to a single-photon mechanism, like its WT counterpart (Figure 8C). Compared to CAN/WT the spectral response of the CAN/R27L mutant to an irradiation pulse more closely resembles a step function. As shown in Figure 8B, the ECN- complexed R27L mutant clearly shows the distinct spectral features of the OCP^1hν^ intermediate, with OCP^R^ formation peaking for *t*_*delay*_ = 1s as observed for wildtype OCP (Figure 8A).

**Figure 8.**
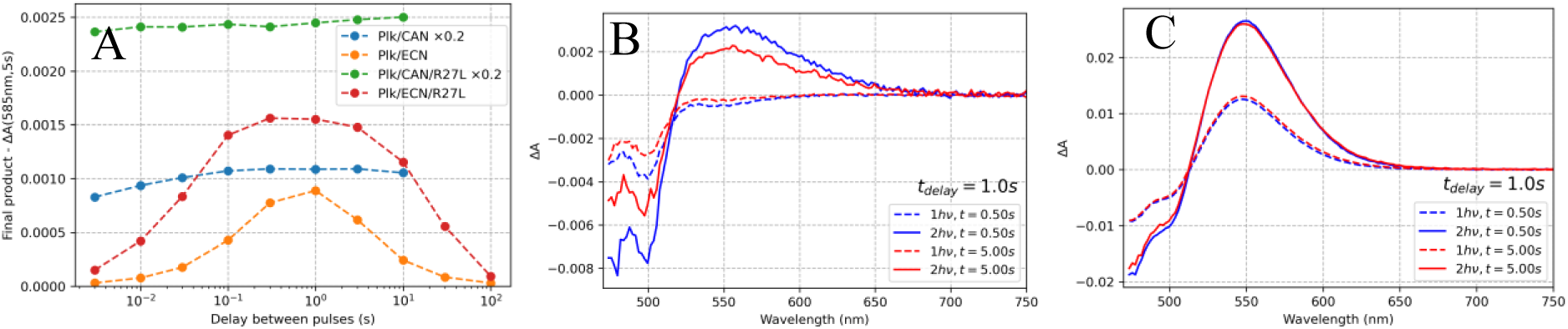
Two-pulse excitation experiment analyzing R27L mutants without His-tag. A) The change in absorption at 585 nm recorded 5 s after the second pulse, representing the yield of OCP^R^, is plotted against the time delay *t*_*delay*_ between the two laser excitation pulses. The results for equivalent complexes of the R27L mutant and WT are shown. B) and C) Transient absorption spectra obtained after one and two 512 nm excitation pulses spaced by *t*_*delay*_=1 s, for the Plk/ECN/R27L and Plk/CAN/R27L mutants, respectively.

Interestingly, for both chromophores the R27L mutant exhibits higher yields after the first and second excitation pulse, respectively, than WT (see Figures 8A, and S26, S27 in Supporting Information for detailed data overview). This may reflect the missing energy barrier involved in monomerization of dimeric WT OCP along the reaction coordinate towards OCP^R^. In conclusion, in ECN-OCP the two absorption events occur within a single monomer in order to yield OCP^R^. Thus, the two-photon feature is not directly associated with the monomerization reaction as proposed by Rose *et al.*^31^.

## Discussion

We investigated the millisecond to second kinetics of the 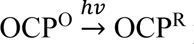 photoconversion using a pump-pump-probe transient absorption setup and found that CAN functionalized OCP forms OCP^R^ upon absorption of a single photon whereas ECN and hECN functionalized OCP requires the sequential absorption of two photons. In the latter two cases the OCP^R^ yield depends strongly on the time delay between the two light pulses, peaking for *t*_*delay*_ = 1 s. This is due to formation of a metastable OCP^1hν^ intermediate that reverts thermally to OCP^O^ if no photon is available within its lifetime or, alternatively, forms OCP^R^ upon absorption of a photon (see Figure 9). It is thus not surprising that OCP is such an enigmatic system since the 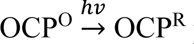 photoconversion of ECN and hECN-functionalized OCPs is “by design” insensitive to single, short light pulses that are a standard temporally well-defined tool widely used in most spectroscopic experiments.

**Figure 9.**
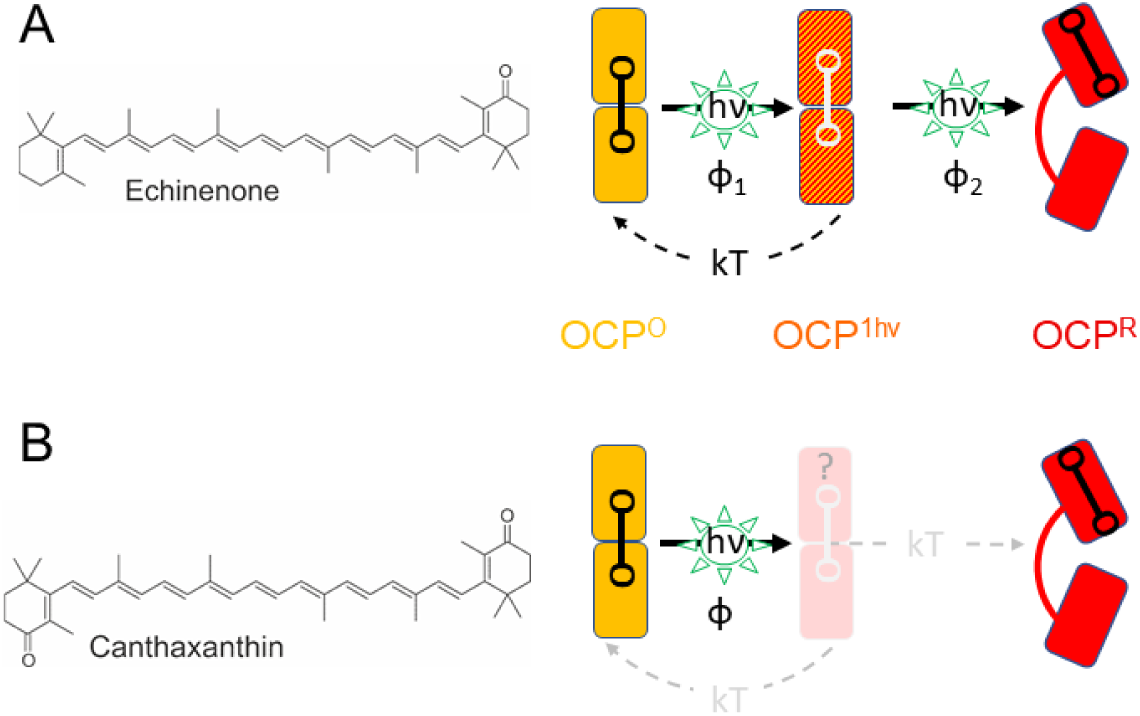
Simplified illustration of the two-photon and single-photon photoconversion mechanisms in OCP. A) ECN (and hECN) functionalized OCP displays a sequential two-photon photoconversion mechanism, involving two absorption events, characterized by quantum yield *ϕ*_1_ and *ϕ*_2_, respectively. Upon absorption of the first photon by OCP^O^ a meta-stable intermediate OCP^1hν^ is formed with *ϕ*_1_. Upon absorption of a second photon by OCP^1hν^, OCP^R^ is formed with *ϕ*_2_. If the light intensity is too low to “provide” another photon within the lifetime of OCP^1hν^ it reverts thermally to OCP^O^. We note that the *ϕ*_1_ and *ϕ*_2_ quantum yields quantify the final outcome of specific cascades of thermally decaying intermediate states (not shown here) to OCP^1hν^ and OCP^R^, respectively. Populating the latter states takes some time since the cascade of prior intermediates needs some finite time to decay and then form these products (resulting in the bell-shaped curve in Figure 6). B) By contrast, CAN functionalized OCP is converted to OCP^R^ upon absorption of a single photon with a quantum yield *ϕ*. We cannot exclude formation of an OCP^1hν^ intermediate with an absorption spectrum identical to OCP^R^ which decays thermally to OCP^R^ as indicated in grey. The OCP chromophore (shown on the left) is visualized by a dumbbell in the scheme, its location in OCP^1hν^ is unknown. For simplicity only monomeric OCP is shown, despite the fact that OCP can be dimeric. We show that the need for two distinct photon absorption events for the photoconversion of ECN-OCP to OCP^R^ does not depend on the oligomerization state. Our data do not indicate when the dimer-to-monomerization step occurs during the reaction.

Rose et al.^31^ recently proposed a two-photon photoconversion mechanism for CAN-functionalized OCP that starts with photodissociation of dimeric OCP^O^ to yield two monomeric intermediates one of which absorbs a photon to form OCP^R^. These intermediates have been reported to have an absorption spectrum almost identical to OCP^R^, except for a 550-630 nm flank that is exclusive to the OCP^R^ form^31^. This is at odds with our observations that support a single photon mechanism for CAN-OCP. In our data, a species possessing a broader absorption band with a 550-630 nm flank is only observed at high pump energy density, but not at 3mJ/cm^2^ (Figure 4). Moreover, at low pump energy density the spectral properties are independent of the time delay *t*_*delay*_ between the first and the second excitation pulse. Therefore, we conclude that there is no reason to assign the occurrence of the 550-630 nm flank to the second photon absorption event. Instead, it is very likely related to a different off-pathway effect such as OCP^R^- OCP^R^ dimerization. Importantly, long-living species without the 550-630 nm flank observed upon using 3mJ/cm^2^ pump pulses have a sufficiently long lifetime and proper spectral signature to be assigned as the OCP^R^. The situation differs for ECN-functionalized OCP which does exhibit a clear sequential two-photon mechanism, involving an OCP^1hν^ intermediate with very different spectral properties than OCP^R^. While we cannot say when monomerization occurs during the reaction, it is clear that it is not a prerequisite for the two-photon mechanism since both CAN and ECN functionalized monomeric R27L-OCPs show the same mechanistic features as the corresponding WT complexes (Figure 8). Moreover, the data on the mutant show that the two photons are absorbed by the same chromophore.

Stepwise two-photon absorption mechanisms in single chromophore molecules have been described for photochromic materials^36,41,42^, but – to the best of our knowledge – not for biological photosensory molecules. The mechanism described here for OCP differs from the one reported for dimeric BLUF-domain containing photosensors that also require two photon absorption events for high enzymatic output yield^43^. In the latter case, each monomer absorbs a single photon resulting in a structural change that is transmitted to the respective enzymatic output domain via the α3 helix that is part of the dimer interface. Thus, like interacting gear wheels the two α3 helices introduce cooperativity, requiring both sensors to be activated for optimal transduction and thus output. The requirement common to both two-photon mechanisms is that the transient one-photon induced species needs to be long-lived and have a large absorption cross section, so that sunlight is capable of completing both steps. Two-photon absorption mechanisms result in a non-linear response of the overall reaction of the light intensity, generating a threshold switch. In case of OCP, it prevents the down regulation of photosynthesis at low light irradiance.

The OCP^R^ forms of both CAN and ECN-complexed OCPs (*Synechocystis* and *Planktothri*x) bind to the phycobilisome and induce fluorescence quenching^10,44^. However, in CAN-OCPs (which accumulate OCP^R^ faster than ECN-OCP), fluorescence quenching is slightly slower and less efficient and the detachment from the phycobilisome is significantly faster than for ECN-OCPs ^10^. This suggests that the structures of the red forms of CAN- and ECN-OCPs differ slightly, which affects the binding to the phycobilisome. The expectation that CAN-OCP^R^ and ECN- OCP^R^ forms are similar allows to speculate that the chromophore-mediated difference in the 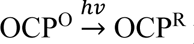 reaction mechanism is rooted in the difference in energy barrier between the one-photon generated intermediate OCP^1hν^ and OCP^R^. If true, this barrier is low enough for CAN-OCP so that it can be crossed thermally, without the need for a second photon (see Figure 9).

CAN, hECN and ECN differ in the β2 ionone ring. Previously, it was reported that the presence of a β2 carbonyl group accelerates the photocycle kinetics of OCP^17^, and influences transient absorption kinetics in the milliseconds time regime^10^. Here we demonstrate that it affects the mechanism of the photoreaction in a fundamental way. Since the carbonyl group in the β2 ionone ring in CAN is very close to Leu37 (Supplementary Fig. S25A), it is conceivable that steric effects may play a role in the mechanistic differences. Therefore, we investigated the L37V and L37A variants of OCP complexed with ECN and CAN. However, we found only a decrease in the OCP^R^ yield, without alteration of the number of photons required to form OCP^R^ (Figures S25B, S22, S23, S24 in Supporting Information). Another property that may play a significant role in the reaction mechanism is the charge distribution along the carotenoid polyene chain: carbonyl groups have an electron-withdrawing property; having one on either side of the chromophore (CAN) instead of only one (ECN) results in a different charge distribution. In the latter case the charge distribution is asymmetric (often associated with the presence of the ICT excited state) which may be key for blocking the single-photon photoconversion pathway in OCP. Further studies are needed to address this hypothesis and to characterize the structural features of OCP^1hν^.

In general, little is known about which carotenoid is bound to natively expressed OCPs *in vivo.* Only three OCPs were isolated from native cyanobacteria cells. *Arthrospira maxima*^45^ and *Synechocystis* 6803 bind hECN^3^. However, when the OCP concentration increases, for example by overexpression, these proteins also bind ECN in addition to hECN^34,46^. By contrast, the *Tolypothrix* (*Fremyella*) OCP, which was isolated from a strain overexpressing OCP, binds CAN. When OCPs are expressed in an *E. coli* heterologous expression system in the presence of CAN and low concentrations of ECN, they all bind CAN and ECN but in different ratios: *Anabaena* OCP binds around 97% CAN, *Synechocystis* around 70-75% CAN and *Arthrospira* 50% CAN^47^. This suggest that the affinity of OCPs for the various ketocarotenoids differs. Thus, which carotenoid(s) is bound to OCP in cyanobacteria *in vivo* depends on the strain specific carotenoids^48^ and their OCP affinities.

Cyanobacteria are diverse organisms that inhabit various environments, including deep ocean layers (down to ≈150 m where there is still some residual sunlight^49^), turbid lake waters (where light can sometimes penetrate only a few millimeters through blooming waters^50^), deserts and even the atmosphere^51^. Consequently, they need responsive photoprotective mechanisms to adapt to these highly diverse and variable conditions. The optimal photosynthetic photon flux density (PPFD) for cyanobacteria is about 30 µmol photons s^-1^ m^-2^, while oxygen production grows linearly with photon flux density up to 100 µmol photons s^-1^ m^-2^ and saturates at about 300 µmol photons s^-1^ m^-2^ ^52–55^. Above these photosynthetic photon flux densities (defined as photon flux density within 400-700 nm, referred to as photosynthetically active radiation - PAR), the photosynthetic apparatus receives an excessive amount of harvested light energy that would lead to damage without photoprotective mechanisms. The corresponding values (marked in Figure 7A, for calculation details see Methods section) overlap very well with the “threshold” regime identified here where the two-photon OCP photoconversion mechanism becomes active.

Interestingly, it was found that *Synechocystis* cyanobacteria exhibit photoinhibition above 800 µmol photons s^-1^ m^-2^ which is reversible up to 1460 µmol photons s^-1^ m^-2^.^56^ (As a reference, full sunlight PPFD at the equatorial equinox corresponds to 2200 µmol photons s^-1^ m^-2^ ^57^.) Therefore, it seems that ECN and hECN-functionalized OCPs are perfectly evolved to moderate the photoprotective function in cyanobacteria. However, it is conceivable that the various carotenoids modulate the “dynamic range” of photoprotection performed by OCPs, thereby extending the range of ecosystems available for cyanobacteria.

## Conclusions

The photoprotective process mediated by OCP is necessary in harsh irradiation conditions because it leads to safe dissipation of excessive excitations which otherwise would flood the photosystems. This, in turn, due to their inability to utilize these excitations productively, would result in large amounts of toxic reactive oxygen species. From an evolutionary point of view, it makes sense to optimize a sensor-protein like OCP such that its efficiency is low, and nonzero only if the triggering light is sufficiently intense and persistent for a long enough time to induce damage. In conditions of weak irradiation, activation of the photoprotective mechanism mediated by OCP is disadvantageous because the presences of the active OCP^R^ species dissipating scarce light energy could lead to starvation. In consequence, the OCP photoactivation mechanism should be as selective as possible with respect to the irradiation intensity. The linear response observed in case of CAN-functionalized OCP will lead to some percentage of active OCP^R^ even in low light conditions, when the dissipation mechanism is generally biologically unfavorable.

By contrast, the two-photon photoconversion mechanism associated with hECN and ECN fulfills the selectivity requirement significantly better. As one can see from the continuous irradiation kinetics (Figure 7B), at low light conditions no OCP^R^ is formed, simply because the absorption events are so rare that the probability of the intermediate OCP^1hν^ form absorbing a second photon is extremely low before it relaxes back to OCP^O^. Only beyond a certain photon flux density threshold does this probability increase enough to obtain a significant OCP^R^ population. We show that ECN-OCP and hECN-OCP effectively photoconvert only above a certain photon flux density that coincidentally agrees with the light conditions where cyanobacteria require photoprotection. Our data demonstrate clearly that any scheme attempting to fully describe the OCP^O^→OCP^R^ photoconversion of ECN or hECN-functionalized OCPs must incorporate at least two reaction quantum yields. It is striking that even after a strong laser pulse of 50 mJ/cm^2^ energy density, allowing for multiple reexcitations, the system cannot undergo an observable photoconversion to the final OCP^R^ product. Remarkably, the latter only occurs upon two excitation pulses delayed by approximately 1 second. To the best of our knowledge, the finding of a sunlight-induced two-photon triggered biologically relevant process occurring in a single protein molecule is unprecedented.

## Methods

### Sample preparation

CAN and ECN functionalized OCP from *Synechocystis* and *Planktothrix* was expressed and purified as described recently^10^. hECN functionalized OCP was expressed as C-terminal His-tagged OCP in *Synechocystis* PCC 6803 and purified as described^10^. The sample was stored at - 80 °C for more than 15 years, with two thawing events during this time. Nevertheless, the protein is functional. To allow removal of the N-terminal poly-histidine tag we introduced a TEV cleavage site right after the last histidine of the tag (protein sequence MGSSHHHHHHENLYFQ^↓^SSFTV), resulting in an additional serine residue before the N-terminal start of the native sequence (SSFTV). The genes for the wildtype constructs containing the additional residues introducing a TEV cleavage site after the poly-histidine tag and the L37V and L37A mutant variants were ordered from Eurofins and cloned into a pCDFDuet™ vector.

We ordered the entire genes for the *Synechocystis* variants. For the *Planktotrix* variants, we exchanged the wildtype sequence between the Nco and Hind III restriction sites of the OCP gene with the synthesized (modified) parts, coding for the additional residues for the TEV cleavage site (see above) and the L37V and L37A mutations, respectively. The correctness of the new constructs was verified by sequencing. All samples used in the two-pulse experiments were concentrated to 20 mg/mL in a buffer containing 40 mM Tris-HCl pH 8.0, 25 mM NaCl. An extinction coefficient of *ε*_500*nm*_ ≈119,000 M^-1^ cm^-1^ was used for protein concentration determination.

### Multi-pulse millisecond transient absorption spectroscopy setup

The experimental setup is based on two noncollinear beams, one used as probe and the second serving as excitation pulse (Figure S1 in Supporting Information shows the geometry of the setup.). Continuous probing light is generated by a Thorlabs MBB1F1 fiber-coupled LED (spectral range 470 - 850 nm), fed through a 200 µm SMA fiber (M92L02, Thorlabs), recollimated using a Thorlabs RC12SMA collimator, cut by an aperture, and focused to a 134×124 µm spot (H×V, FWHM) in the cuvette. The aperture blocks the majority of the probe light before the sample, ensuring very low intensity probing conditions. This light is collected and recollimated through a AC127-030-A-ML lens after the sample and fed to a thick multi-fiber SMA optical fiber using a F810SMA-543 collimator. This fiber is terminated at the entrance slit of an Andor spectrometer coupled with a Newton Peltier-cooled camera operating in unamplified mode (1600 pixels, 150 groves/mm grating). The probe light is efficiently collected, and its intensity adjusted such that the camera operates much below saturation threshold. The camera is operated in full vertical binning mode.

Pump pulses are produced by an Oxxius LBX-520-800-HPE-PP laser diode module, fed to the SMA 200 µm fiber (M92L02, Thorlabs), then recollimated using a F810SMA-543 collimator and focused in the sample using an AC127-050-A-ML lens, yielding a 193×185 µm spot (H×V, FWHM). These pulses are triggered by a TTL trigger; the length determines the pulse energy. The temporal shape of this pulse is fast-growing followed by an exponential decay with a time constant of ≈290 µs.

A 100 µm pathlength quartz cuvette (Hellma) is mounted using a custom-made aluminum magnet-secured holder (see Figure 1A) which guarantees that the sample is always in the plane of the XY stage motion (the third axis is always fixed at the probe/pump focal plane) and that the sample mounting repeatability is very high. The experiment is performed in complete darkness at 22±1℃, stabilized by air conditioning.

All triggers (camera trigger, laser pulse trigger and probing LED trigger) are generated by the ATmega32U4 microcontroller located on an Arduino board. A special procedure written in C uses internal counters in order to generate a proper trigger sequence for the camera and laser module with sufficient temporal precision dictated by onboard quartz oscillator (16 MHz). A custom Python-written procedure executes the experimental sequence, including movements of the XY axes and communicating the proper trigger setting to the Arduino board before each kinetic is being recorded. The Andor Solis software runs independently in a loop and after each camera trigger saves the acquired kinetic into a *.sif file marked by incremental number.

The experimental sequence consists of 4 cycles executed repeatedly for a sequence of delays *t*_*delay*_ (see Fig. 10). Delays of 3 ms, 10 ms, 30 ms, 100 ms, 300 ms, 1 s, 3 s, 10 s, 30 s, 100 s were used. In the first cycle, the probing light is switched on and a kinetic time course is collected (spectra recorded with sampling period 2.61 ms). In the second cycle, a kinetic time series with one pump pulse is collected (the pump pulse is generated 2.5 s after the probe is switched on). In the third cycle, two pulses with predetermined *t*_*delay*_ between them are used to pump the sample. The second of these pulses is aligned such that it occurs 2.5 s after switching on the probe light, meaning that for long *t*_*delay*_ delays only the evolution after the second pulse will be recorded. This is necessary, otherwise, there would be a large effect of probe exposure for long *t*_*delay*_ delays. Last, the fourth cycle is dark, no pump or probe light is present. All generated pulses are identical.

**Figure 10.**
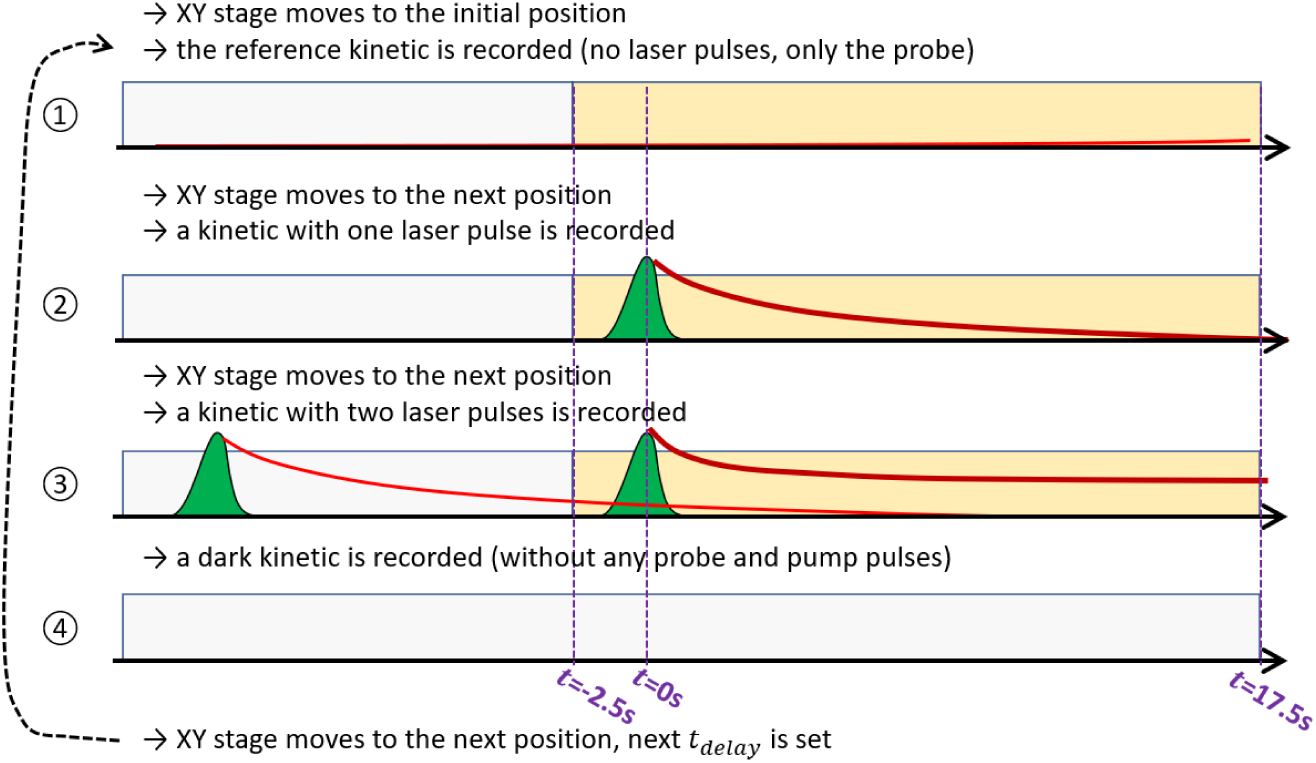
The experimental sequence consists of 4 steps performed repeatedly for a grid of *t*_*delay*_ values.

These 4 cycles are repeated multiple times for each of the 10 delays investigated (3 ms, 10 ms, 30 ms, 100 ms, 300 ms, 1 s, 3 s, 10 s, 30 s, 100 s) such that after the last delay the system returns to the first one, and continues until a sufficient amount of data is accumulated (usually 1000- 2500 cycles in total). After each cycle, the probe and pump are switched off and the sample XY stage is translated by 1 mm (or to the next row separated vertically by about 500 µm) in order to always probe a fresh sample aliquot (except after the 3^rd^ cycle because the sample does not need to be refreshed to record a correct dark kinetics). Because the sample surface is limited, the stage will revisit the same place after a very long time (typically slightly more than 3 hours and 30 minutes).

This sequence was designed so that a large amount of control data is recorded; in particular 10 times more data is recorded for 1-pulse excited kinetics than for 2-pulse kinetics, which is important in data post-processing.

Each kinetics dataset starts 2.5 s before the first pulse (or the second pulse in case of step ③) and ends 17.5 s after this pulse, so that the entire time window is 20 s. We limited the experiment to this time period because the absence of a probe influence on the data cannot be guaranteed for longer time spans. The probe intensity was carefully minimized; however, it is still a focused beam, and it must have non-negligible intensity to obtain good enough spectra. Unfortunately, when using continuous probing, the probe effects accumulate over time (especially in a pulse- insensitive system like OCP) whereas pulse-induced effects are imprinted only once or twice on the sample. The probe light spectrum was selected such that it has a very small amount of light below 520 nm to avoid interaction with the sample. The disadvantage of this approach is the poor quality of the spectra at the carotenoid bleaching region.

The recorded data are binned and averaged to calculate: i) one dark kinetic from step ④, ii) one reference kinetic from step ① (only probe no pump), iii) one single pulse kinetic from step ② (probe and one pump pulse), iv) 10 kinetics recorded for each *t*_*delay*_ in step ③. The following grid of *t*_*delay*_ values was used 3 ms, 10 ms, 30 ms, 100 ms, 300 ms, 1 s, 3 s, 10 s, 30 s, 100 s.

Before averaging, the intensities were median-filtered (exemplary diagnostics are shown in Figure S2 in Supporting Information). The averaging procedure takes probe light intensity at 700 nm, composes the distribution of this value for all kinetics collected in given bin and calculates the median value. Then, averaging is done using all kinetics except those that deviate by more than 5% from the median. This kind of filtering allows to discard data from air bubbles that form after some time in the cuvette. Usually no more than 10% of the kinetic time points were discarded this way. After binning, the averaged intensities of the light are used to calculate the transient absorption signal after a single and two or more pulses using the formula:

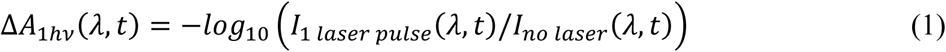

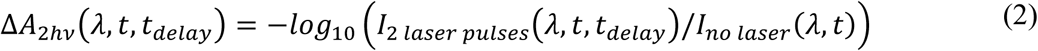

The above formula assumes that dark intensities were already subtracted. The transient absorption obtained for *t*_*delay*_ delays up to 10 s are zeroed at 100 ms before the first pulse in a given cycle. Zeroing is done to compensate for long-term probe drifts occurring during the experiment and random differences in sample absorption in the different parts of the cuvette. In kinetics involving two pulses (step ③), both pulses are within the probing window only for *t*_*delay*_ < 2.5 s. Therefore, for longer *t*_*delay*_ delays, the 1-pulse kinetic (recorded in step ②) is used to correct the two-pulse kinetic properly, to ensure that ΔA = 0 is set 100 ms before the 1^st^ pulse. Unfortunately, for *t*_*delay*_ >17.5 s (*t*_*delay*_=30 s and 100 s) such correction is not possible, because the 1-pulse kinetics is too short. Therefore, for these two kinetics ΔA = 0 was exceptionally set 100 ms before the 2^nd^ pulse. In other words, in those two cases we used the absorbance before the 2^nd^ pulse as approximation of the absorbance before the 1^st^ pulse. Based on the one pump pulse data, one can see that the error introduced by this approximation is virtually zero for probe > 525 nm in ECN-OCP and hECN-OCP proteins. Therefore, this correction does not affect absorbance at 585 nm which is the most relevant parameter. There is some residual negative signal for wavelengths < 525 nm remaining at 17.5 s after the pump pulse, which slightly affects the shape of the calculated spectra below 525 nm produced for *t*_*delay*_=30 s and 100 s. In CAN proteins, there are long-living products even after a single pulse. Therefore, only kinetics with *t*_*delay*_≤10 s are presented.

Finally, the signal derived from buffer is subtracted from the Δ*A* OCP sample kinetics to remove any effects unrelated to the protein itself. In conclusion, the experiment was designed to consider all tradeoffs and fulfill all criteria necessary to obtain reliable kinetics for OCP in the temporal window investigated (long sample relaxation time, practical absence of sample diffusion and sufficient S/N ratio).

### Absorbance changes upon continuous irradiation determined in a stationary UV-vis spectrometer

The differential quantum yield was calculated using the following formula:

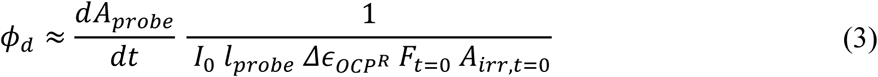

Where the photokinetic factor *F* is defined as:

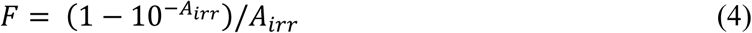

*A*_*irr*_ is the sample absorbance at the irradiation (LED) wavelength (with 4 mm irradiation pathlength), *Δ∈*_*OCP*_*R* is the differential molar absorption coefficient defined as *Δ∈*_*OCP*_*R* = *∈*_*OCP*_*R* − *∈*_*OCP*_*O*, *l*_*probe*_ is the probing path length (10 mm), *I*_0_ is the photon flux density of the irradiation LED, *A*_*probe*_ is the absorbance measured using a 10 mm probing path length. Because the used LED (Thorlabs M470L5) has a broad spectrum, the *I*_0_ *F*_*t* = 0_ *A*_*irr,t*=0_ term was integrated over the LED spectrum and the OCP^O^ absorption spectrum, respectively, to take into account different absorbances at different spectral components of the irradiation light. Therefore, the above equations can be rewritten as:

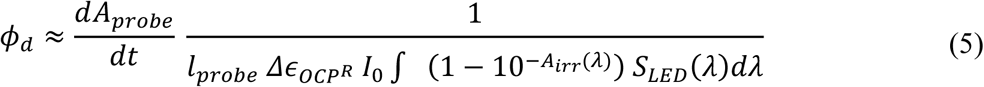

Where *S*_*LED*_(*λ*) represents the normalized spectrum of the Thorlabs M470L5 LED, so that ∫ *S*_*LED*_(*λ*)*dλ* = 1. Nevertheless, the difference in correction obtained using eq. 5 vs eq. 3 is very minor. A quartz cell of 4 × 10 mm optical path lengths was utilized (applicable for irradiation and probing, respectively), the sample temperature was stabilized at 22±1 ℃. The irradiation LED light was collimated and directed towards the sample (perpendicular to the probing beam). A Jasco V-650 spectrophotometer was used for probing with an additional 550 nm bandpass filter placed after the sample cuvette to remove scattered irradiation light. Each kinetics was recorded with 30 s irradiation time. Then a linear regression was performed using the linear part of each kinetics (see Fig. S3 in Supporting Information). In case of high irradiation intensity, the recorded absorption signal starts to saturate for some kinetics after only a few seconds of irradiation. Therefore, in this case only the initial datapoints were used. The irradiation intensity was determined using Gentec PH100-Si-HA detector head.

Maksimov *et al.*^13^ reported a molar absorption coefficient of 87 270 M^-1^ cm^-1^ (495 nm) for ECN-functionalized OCP from *Synechocystis* (using a triple mutant devoid of tryptophans except one, Trp-288). Using this molar absorption coefficient, we derive a differential molar absorption coefficient of Δɛ(550 nm) = 29525 M^-1^ cm^-1^ (taken from Figure 1).

### Conversion between PAR and monochromatic photon flux density

In the cyanobacteria literature, the light intensity is quantified using the unit of photosynthetic photon flux density (PPFD) which is a measure of photosynthetic active radiation (PAR). It takes into account only sunlight in the 400-700 nm spectral range. In the experiments described here we used a 472 nm LED to irradiate the sample. Its spectrum is much narrower (however not monochromatic) compared to the spectrum of sunlight in the PAR range. Thus, because the majority of the PAR radiation falls outside the carotenoid absorption spectrum, a direct comparison between the PPFD values reported in the literature and the photon flux densities given here is not meaningful. In order to calculate a conversion factor between these values, we first calculated the photosynthetic photon flux density (assuming PAR spectral range) that results in absorption of 1 photon per carotenoid per second on average. Then, we calculated the photon flux density assuming 472 nm LED light, resulting in the same condition of 1 photon absorption per carotenoid per second. The ratio between these values yields the desired conversion factor. It allows to compare the literature values describing cyanobacteria functioning with the values obtained here describing OCP functioning. For PAR, we assume a standard AM1.5G spectrum^58^ (given in W m^-2^ nm^-1^ as a function of wavelength) truncated to the 400-700 nm range and recalculated into mols photons m^-2^ nm^-1^ and normalized to unity, so that:

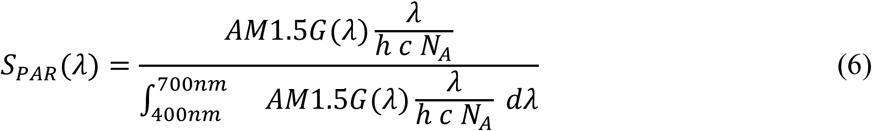

Where *h* is the Planck constant, *c* the velocity of light, *N*_*A*_the Avogadro number. A similar normalized spectrum *S*_*LED*_(*λ*) was calculated for the 472 nm LED source (Thorlabs M470L5). Both normalized spectra have units of nm^-1^. The number of photons absorbed per molecule per second is calculated using formula:

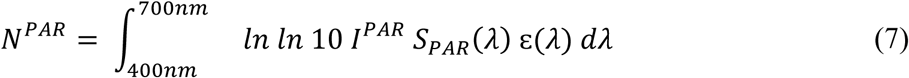

Note that the photosynthetic photon flux density *I*^*PAR*^ (mol m^-2^ s^-1^) and molar absorption coefficient of the carotenoid ɛ(*λ*) (mol^-1^ dm^3^ cm^-1^) are defined using different units, so an additional factor of 0.1 is necessary to ensure proper unit conversion. Using these equations one can calculate a conversion factor between PPFD and the photon flux density of the 472 nm LED by assuming *N*^*PAR*^ = *N*^*LED*^:

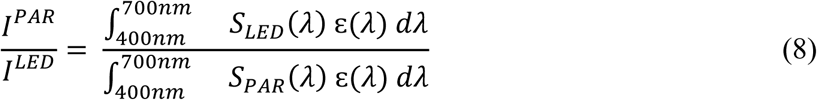

It allows to convert between the photon flux densities given for both types of radiation, ensuring that the considered quantities are effectively the same from the carotenoid’s perspective, *i. e*. they result in the same number of photons absorbed. It can be calculated that for a PAR spectrum and photosynthetic photon flux density of *I*^*PAR*^=148 µmol s^-1^ m^-2^ there is one photon absorbed per molecule per second on average (so that *N*^*PAR*^=1 s^-1^). For the 472 nm LED spectrum (Thorlabs M470L5), and photon flux density of *I*^*LED*^=54 µmol s^-1^ m^-2^ there is also one photon absorbed per molecule per second on average (so that *N*^*LED*^=1 s^-1^). The conversion factor calculated using formula (8) is equal to *I*^*PAR*^/*I*^*LED*^=2.8. It means that a 2.8-fold lower photon flux density is needed when using the LED compared to sunlight to result in the same number of photons absorbed.

## Data availability

Data will be provided upon reasonable requests.

## Code availability

The custom written code has been deposited at https://github.com/Stahux/MultipulseTransientAbsorption.

## Acknowledgments

We are very grateful to Thomas R. M. Barends for expert support during the experiment development, to Herbert Zimmermann for expert help with literature search and to Chris Roome and Mario Hilpert for assistance with data storage and other IT issues. We thank Klaus Brettel for sharing preliminary data on similar photoconversion experiments. We acknowledge fruitful discussions with Jacques-Philippe Colletier at the initial stages of the project.

## Author contributions

S.N., G.B., I.S., D.K. and M. S. conceived the research. I.S. supervised the project. S.N. designed the experimental setups, performed experiments and analyzed the data. E.H. and A.W. prepared samples. S.N. wrote the initial version of the manuscript which was expanded by S.N., I.S., D.K., M.S. and G.B. R.L.S. and J.R. helped with the experiment development. All authors contributed to discussions and the writing of the manuscript.

## Competing interests

The authors declare that they have no competing financial interests or personal relationships that could influence the work reported in this paper.

## Notes

### Competing Interest Statement

The authors have declared no competing interest.

