## Supporting information for "Two-photon driven photoprotection mechanism in echinenone-functionalized Orange Carotenoid Protein"

### Supplementary text

*Sample heating by the pump laser pulses*

Strong irradiation pulses deposit a lot of energy in the sample which may result in sample heating. Ignoring the Gaussian spatial profile of the pump pulse, an area of 193 µm · 185 µm ≈ 0.036 $mm^2$ is excited. The sample layer thickness is 100 µm, therefore the excited volume is approximately 0.0036 $mm^3$ = 0.0036 µL. Assuming that the sample's density is close to water's density, this corresponds to a weight of $3.6 \cdot 10^{-9}$ kg. When assuming a pulse energy density of 50 $mJ/cm^2$ and no pulse transmission through the sample, $1.8 \cdot 10^{-5}$ J will be deposited in the excited volume. Assuming a specific heat capacity of 4184 J $kg^{-1}$ $K^{-1}$ (ignoring the non-water content of the sample), the temperature increase of the sample is about 1.2 °C after one excitation pulse. This value is not negligible, but much too small to significantly alter the sample properties, including its kinetics. It also cannot explain the observed "memory" effect of OCP, in particular when the excitation pulses are separated temporally by several seconds. The heated sample aliquot is surrounded by glass and "cold" sample solution, therefore efficient cooling will lead to fast heat dissipation. Both the ECN-OCP and CAN-OCP samples are almost identical from the thermal point of view, however they exhibit completely different behavior after the strong excitation pulse.

*Synopsis of the transient absorption spectra shown in the Supplementary Information*

The data shown in the supplement belong to experiments described in the main text, such as investigations on OCP from *Planktothrix* or mutant variants, but also control studies addressing the influence of the pump laser energy density or the wavelength of the pump pulse. In the following text, proteins from *Synechocystis/Planktothrix* will be abbreviated as Syn/Plk, followed by their chromophore; e.g. Syn/ECN is OCP from *Synechocystis* complexed with echinenone.

Similar to the figures shown in the main text, the figures depict transient absorption spectra recorded 500 ms and 5 s after the single pump pulse (dashed lines) and after the second of the two pulses (continuous lines, always for a specific $t_{delay}$). For each dataset, we also show kinetics probed at 585 nm after two pump pulses and transient absorption spectra recorded after a single pump pulse for various delays. The probing time window extends to 17.5 s after the laser pulse. We show only data up to 5 s, for which we are certain that no significant probe effect is present. However even in the full 20 s probing time window, effects of the probe are rather minor (see Figure S5). Since the probe is continuous, its effects accumulate in the sample over an extended period of time (t >> 5 s). The goal of these experiments was to capture effects of the photoconversion, not the decay kinetics. In order to record the full decay kinetics, it may be advantageous to use single probe pulses to completely avoid probe effects. This, however, will require significantly more repetitions of the experiment, which is why we used continuous probing.

*Influence of pump pulse parameters – energy density, wavelength, number of pulses*

Figures S10, S11 and S12 present data obtained for Plk/ECN using pump laser energy densities of ~50 mJ/cm$^2$, ~12 mJ/cm$^2$ and ~3 mJ/cm$^2$, respectively. One can see that when the laser energy density increases from 3 mJ/cm$^2$ to 12 mJ/cm$^2$, the 490 nm band intensity only doubles and stays almost the same when increasing the energy density from 12 mJ/cm$^2$ to 50 mJ/cm$^2$. This indicates that either the OCP$^O$ population is almost completely depleted by the 50 mJ/cm$^2$ excitation pulse, or, alternatively, that there is a limiting process that prevents the full transition of the OCP$^O$ population to the OCP$^{1hv}$ form. Note that for all excitation energies, the second pulse is always capable of doubling the 490 nm signal amplitude present after the first pulse. This observation supports the second possibility – there must be some process limiting the number of molecules photoconverted to the OCP$^{1hv}$ form per time unit. It is clear that this process acts on the submillisecond time scale (the laser pulse duration), but ceases to occur at longer timescales ($t \geq$ 3ms – the shortest used $t_{delay}$). This could be an additional inactive dark form in which the majority of the photoexcited OCP is trapped which needs time to decay and that otherwise cannot productively photoconvert towards OCP$^{1hv}$. Alternatively there may be a preferred back photoconversion pathway between an intermediate preceding OCP$^{1hv}$ (which we denote as OCP$^X$) and OCP$^O$, so that the OCP$^X \overset{hv}{\rightarrow}$ OCP$^O$ photoconversion is much more probable than the OCP$^O \overset{hv}{\rightarrow}$ OCP$^X$ photoconversion. Even if the observed nonlinearity is due to the full depletion of the OCP$^O$ population caused by a strong irradiation pulse, the second light-triggered step must be characterized by an extremely low quantum yield. The reason is that regardless of the excitation energy used, the band at 550 nm never significantly exceeds 2 mOD. Due to the negative feature at 545 nm, it is not easy to observe the OCP$^R$ signature (expected at about 550 nm) when using low excitation energy density. At 12 mJ/cm$^2$, not much of the OCP$^R$ form is observed, and for 3 mJ/cm$^2$ it is not detectable. We are convinced that this is due to the combination of the low quantum yields of both photoconversion steps and the existence of the aforementioned limiting processes. Therefore, we conclude that OCP has evolved to be insensitive to short light perturbations (there are more than one possible implementation of such effect). The practical consequence is that the possible number of “productive” photoconversion effects per time unit is finite and independent of the irradiation intensity.

Figures S13 and S14 show datasets acquired using a 488 nm excitation pulse with an energy density of approximately 50 mJ/cm$^2$ and 12 mJ/cm$^2$, respectively. The signals obtained with 488 nm excitation have comparable spectral shapes and intensities to ones obtained with 512 nm excitation. This excludes an effect of the OCP$^O$ heterogeneity on the photoconversion mechanism[1,2]; different OCP dark-adapted subpopulations photoconvert to OCP$^R$ form in the same way.

Figure S15 demonstrates how a larger number of pulses leads to buildup of the $OCP^R$ spectral signature in Plk/ECN observed after irradiation with continuous light. It shows that with an increasing number of excitation pulses, the peculiar negative spectral band observed after only one excitation pulse becomes more and more obscured by the well-known symmetrical $OCP^R$ signature (with both negative and positive transient absorption contributions). Note that the negative 490 nm and positive 550 nm bands grow differently with the number of excitation pulses. This graph closes the gap between discrete and continuous light regimes, demonstrating the advantage of the two-pulse excitation approach.

Figures S7, S16 and S17 present data obtained for Syn/CAN using pump laser energy densities of ~50 $mJ/cm^2$, ~12 $mJ/cm^2$ and ~3 $mJ/cm^2$, respectively. As discussed in the main text, when using excitation pulses with the lowest energy density of ~3 $mJ/cm^2$, the second excitation pulse results in the same absorbance change as the first pulse, regardless of $t_{delay}$. This, however, does not hold for higher excitation energy densities. After ~12 $mJ/cm^2$ and ~50 $mJ/cm^2$, the second excitation pulse results in a higher absorbance change compared to the first pulse, and there is a small but consistent dependency of $t_{delay}$. The quantification of the amount of $OCP^R$ product as a function of $t_{delay}$ is visualized in Figure S19. Note also, the signal magnitude scales less than linearly with the excitation pulse energy density, indicating saturation and presence of some "limiting processes" similar to ones already discussed above for ECN-functionalized OCPs. Another effect present only for excitation energy densities above ~3 $mJ/cm^2$ is the faster decay of signals observed after the pair of excitation pulses, compared to signals observed after only one pulse (Figure S18).

*Results obtained on modified proteins – tags, mutations*

The rationale behind the experiments as well as the choice of mutants is described in the main text. Figures S20 and S21 show the results obtained from tagged Syn/ECN variants (N- and C-terminal His-tags, respectively). The behavior of these samples differs from tag-free Syn/ECN (Figure S4): after one excitation pulse one can observe a spectral signature that resembles $OCP^R$ (positive band close to 550 nm). Nevertheless, it is evident that the photoactivation mechanism still requires two photons because this band decays completely within 1 s, clearly indicating that it is not associated with an $OCP^R$ form. The $OCP^R$ signature lives only long enough to be considered as a bona fide $OCP^R$ form, when a properly delayed second excitation pulse has been applied.

Figures S22 and S23 show the results obtained for the L37V mutants (Syn/ECN/L37V, Syn/CAN/L37V, respectively). These mutant variants display significantly lower signal amplitude than WT. Nevertheless, the mutation preserves the type of photoconversion mechanism, indicating that the steric interaction between the β2-keto group of the carotenoid and the L37 residue is not a factor that decides whether the photoconversion mechanism is two or single photon. This conclusion is further confirmed by the results obtained for the L37A mutant (Syn/CAN) shown in Figure S24. Crystal structure analysis of the L37V and L37A mutants complexed with ECN/CAN, respectively, show no structural changes compared to wildtype (data not shown). Figure S25 shows the mutation site in the protein structure and the dependence of the $OCP^R$ product yield of $t_{delay}$.

Figure S26 shows the results obtained for the Plk/ECN/R27L mutant. The R27L mutant protein is known to be monomeric due to disruption of the R27-D19$^{\#}$ interaction at the dimerization interface[3] ($^{\#}$ indicates a residue in the other monomer in the OCP dimer). Compared to WT (Figure S10), the signal is about two times higher (note that an accurate comparison of the yield is not possible because the signal depends also on the accuracy of the pump-probe spatial overlap and the sample absorbance which is not always exactly identical). The R27L mutation does not result in an opening of the single-photon channel, demonstrating that the two-photon effect is not rooted directly in the dimeric state of OCP. The dimeric WT OCP needs to dissociate into monomers upon photoexcitation to enable the CTD-NTD domain separation required for $OCP^R$ formation[4,5]. As this dissociation process requires energy, it is expected that the $OCP^R$ yield is reduced to some extent in a dimeric versus monomeric protein. Nevertheless, since R27L mutant requires two properly timed excitation pulses in order to photoconvert to the $OCP^R$ form, it is clear that the two-photon mechanism must be rooted in something else than the requirement of dimer dissociation.

Figure S27 shows results obtained for the Plk/CAN/R27L mutant. It has the same photoconversion characteristics and thus mechanism as WT (Figure S6). The main difference is an increase of the signal magnitude, analogously to the Plk/ECN/R27L sample.

**Supplementary Figures**

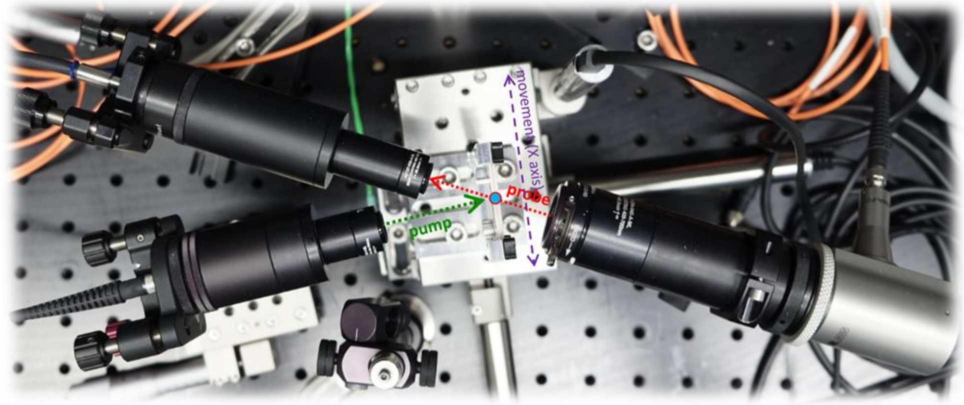


**Figure S1**. Top-view of the experimental setup geometry.

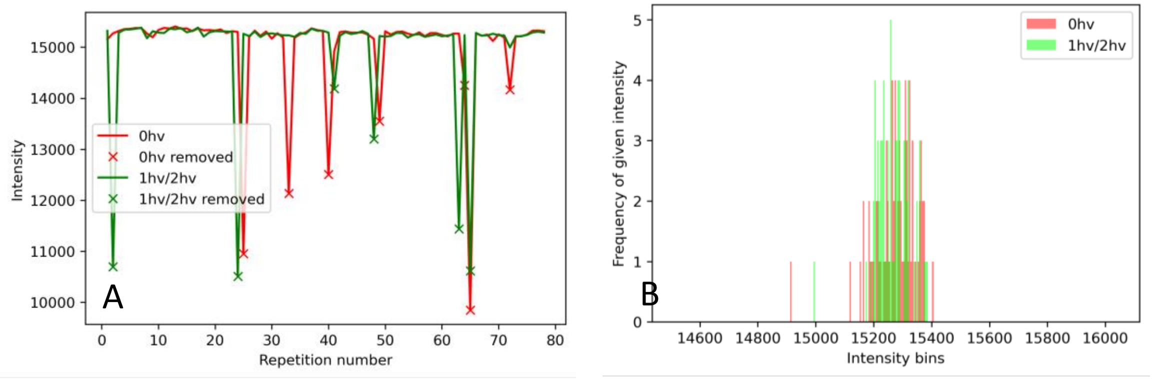


**Figure S2**. Exemplary diagnostics graphs (routinely checked for all datasets) demonstrating the effect of median filtering. The graphs show superimposed data obtained from cycles with one excitation pulse and cycles with the probe only. The plotted probe intensity was recorded at 700 nm, about 8 ms after triggering the camera acquisition (*i. e.* before the excitation pulse is generated). A) Negative spikes are due to small air-bubbles formed within the cuvette that act like a spherical lens and defocus the probe beam, causing a signal drop. Data points removed from the kinetic series are marked by “X”. B) Histogram of signal intensities centered at the median (distant outliers fall outside of the graph window).

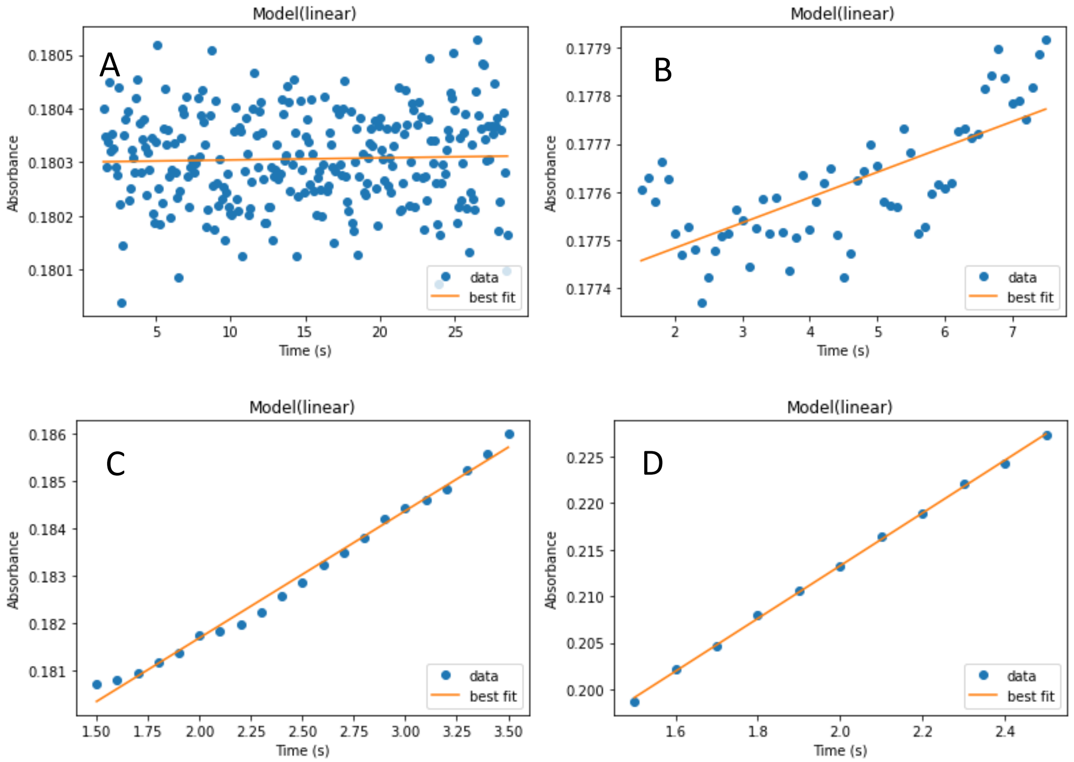


**Figure S3**. Exemplary linear regression fits of the absorbance during the first seconds of irradiation of a Syn/ECN sample. The irradiation intensities are A) 5.22, B) 50.3, C) 348, D) 2289 µmol photons $s^{-1}$ $m^{-2}$.

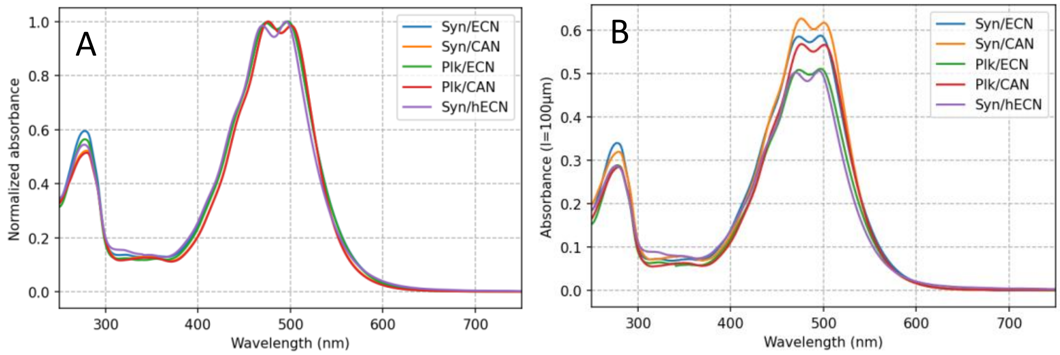


**Figure S4**. UV-vis absorption spectra. A) Normalized UV−vis stationary absorption spectra of OCP samples used to calculate their respective differential quantum yields[6]. The measurement was performed using a sample in a 4×10 mm cuvette. B) Unnormalized spectra of the samples as used in the two-pulse excitation experiment, measured in a 100 µm pathlength cuvette (just before the experiment).

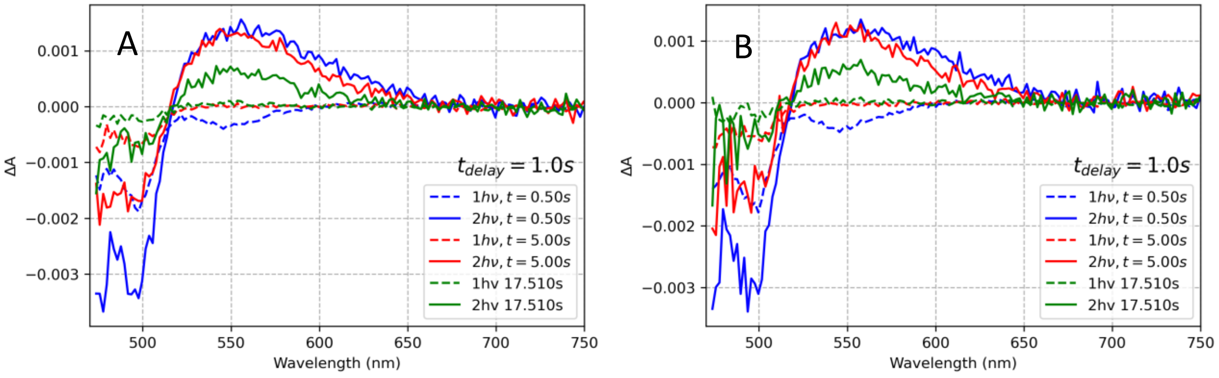


**Figure S5.** Comparison of two different probing intensities used in a two-pulse experiment on *Planktothrix* OCP functionalized with ECN, with His-tag removed (two different cuvette fillings, therefore the concentration may slightly differ). Excitation at 512 nm, energy density about 50 mJ/cm$^2$. A) Standard probe used in all experiments, B) the same probe attenuated by the factor of two.

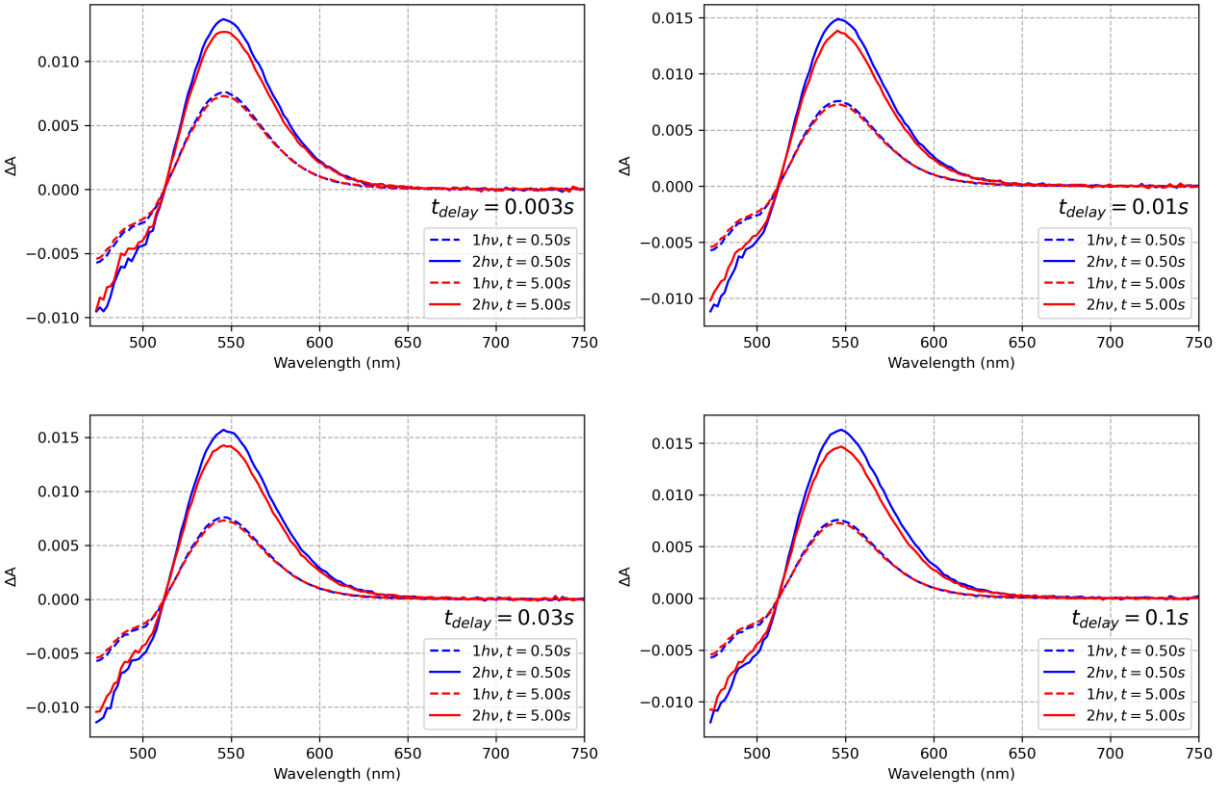

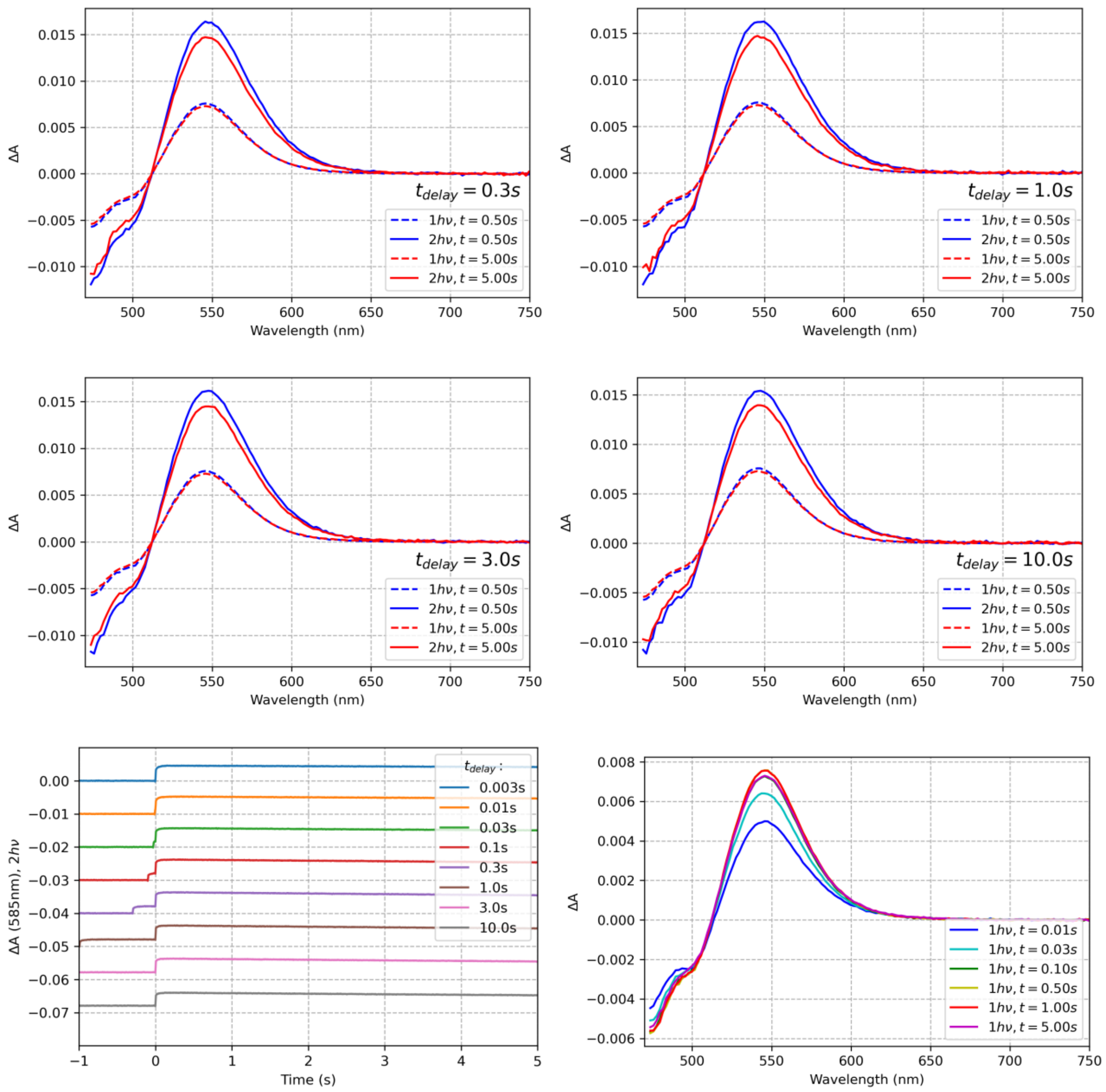


**Figure S6**. Two pulse experiment results obtained for OCP from *Planktothrix* functionalized with CAN, with the His-tag removed. Excitation at 512 nm, energy density about 50 mJ/cm$^2$.

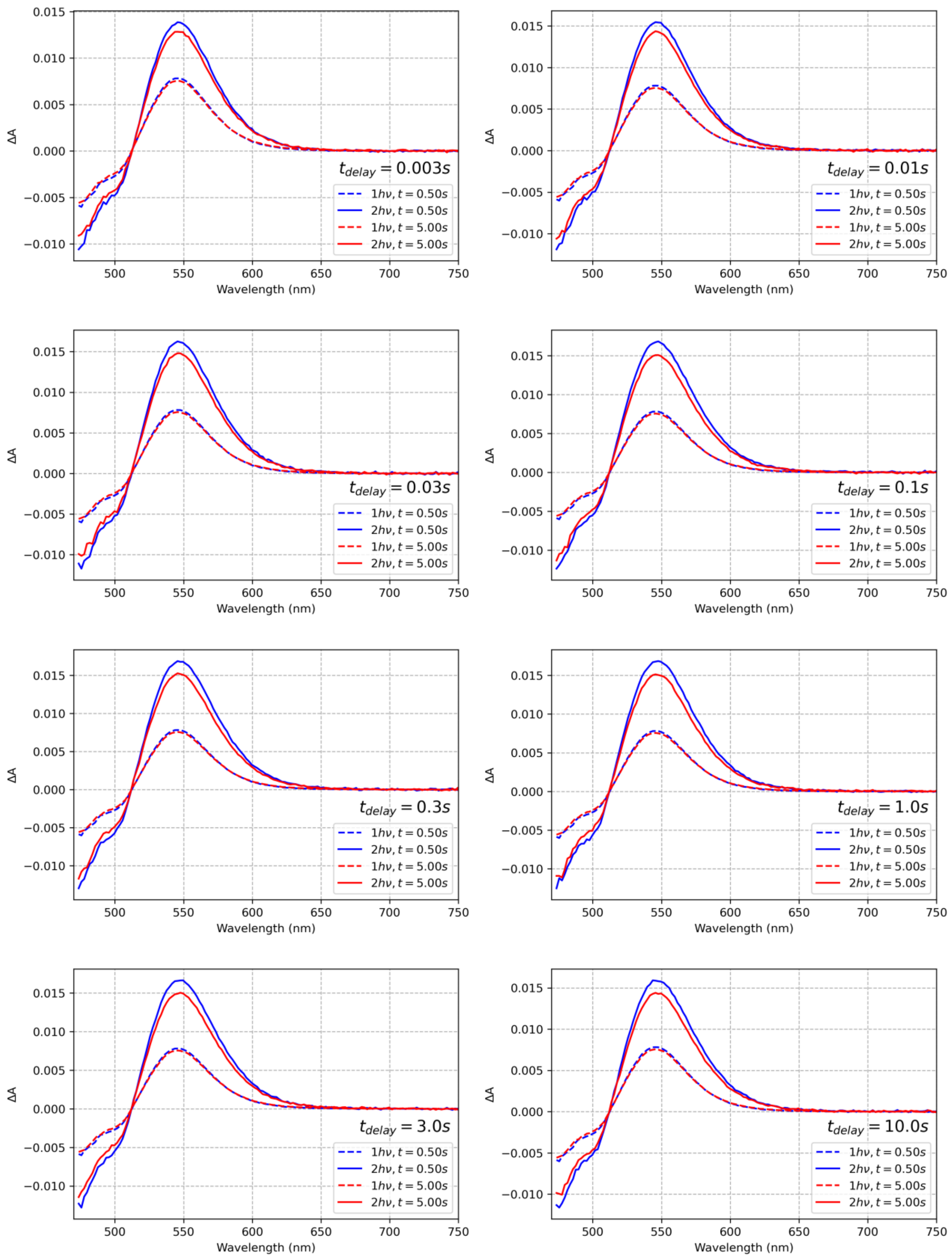

ΔA
Wavelength (nm)
$t_{delay} = 0.003s$
$t_{delay} = 0.01s$
$t_{delay} = 0.03s$
$t_{delay} = 0.1s$
$t_{delay} = 0.3s$
$t_{delay} = 1.0s$
$t_{delay} = 3.0s$
$t_{delay} = 10.0s$
$1h\nu, t = 0.50s$
$2h\nu, t = 0.50s$
$1h\nu, t = 5.00s$
$2h\nu, t = 5.00s$

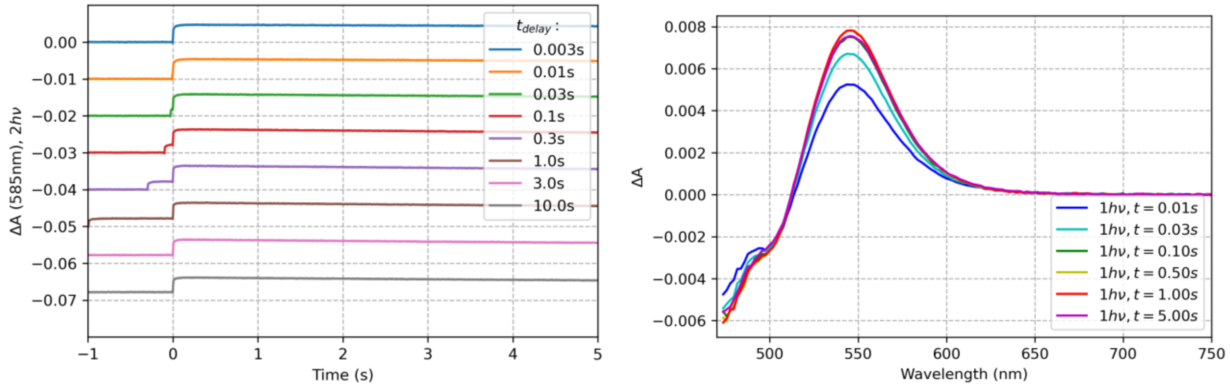


**Figure S7**. Two pulse experiment results obtained for OCP from *Synechosystis* functionalized with CAN, with His-tag removed. Excitation at 512 nm, energy density about 50 mJ/cm$^2$.

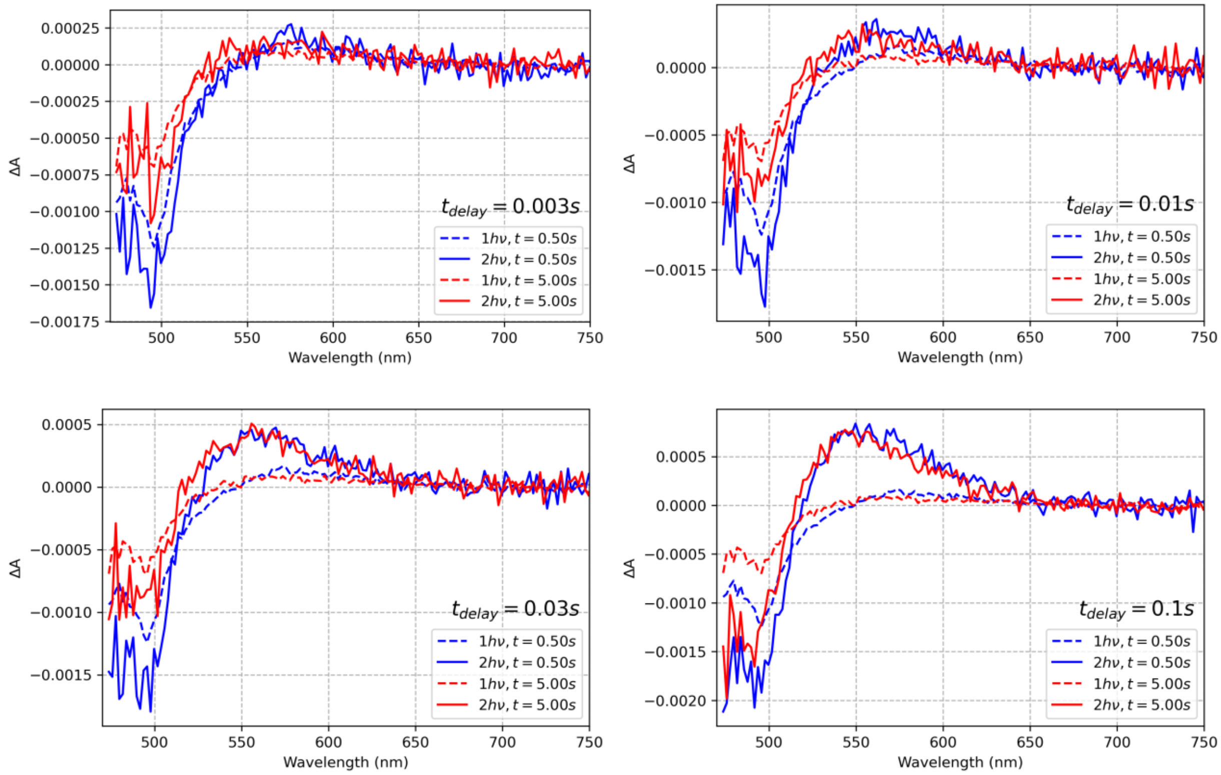

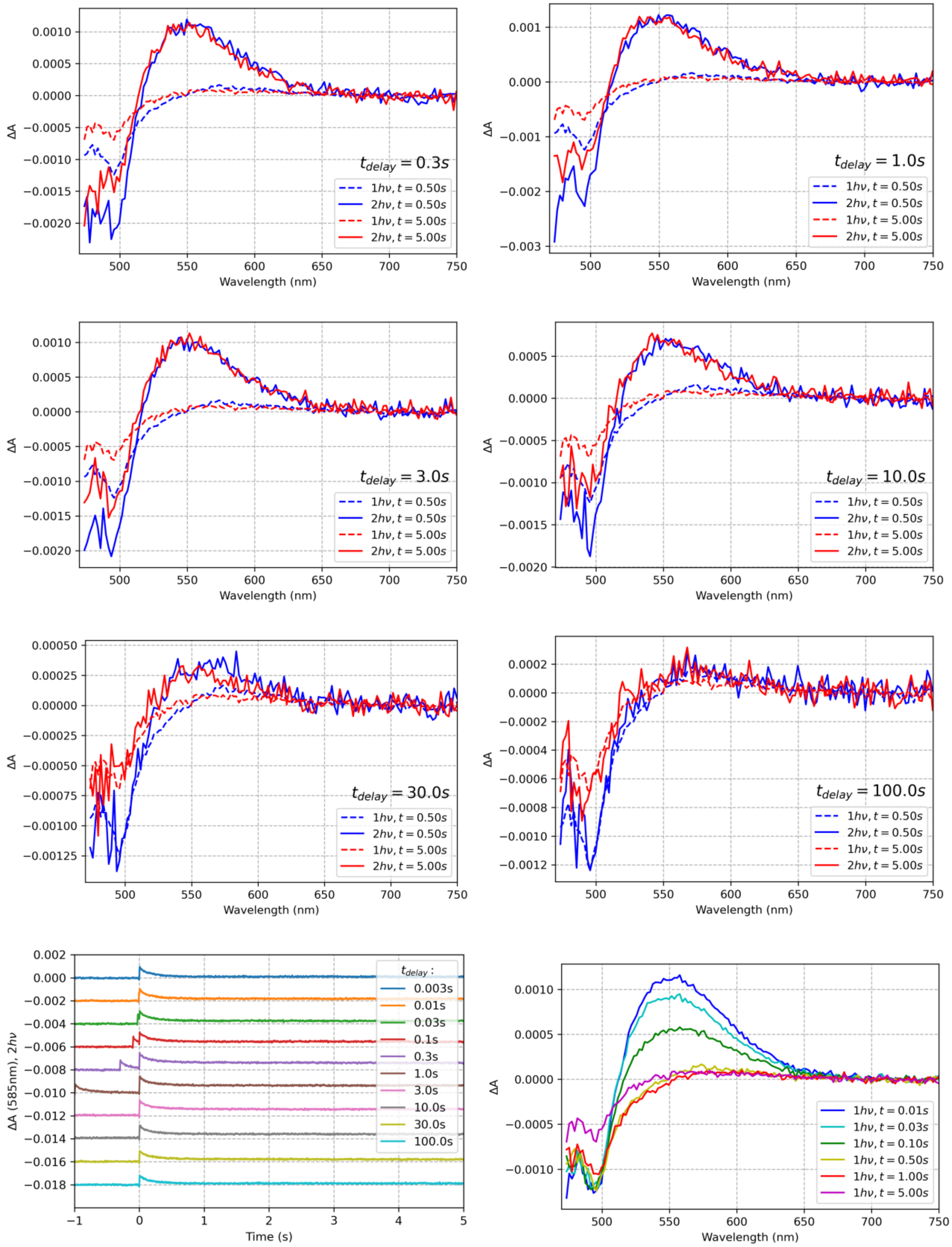


**Figure S8**. Two pulse experiment results obained for OCP from *Synechosystis* functionalized with hECN, with His-tag removed. Excitation at 512 nm, energy density about 50 mJ/cm$^2$.

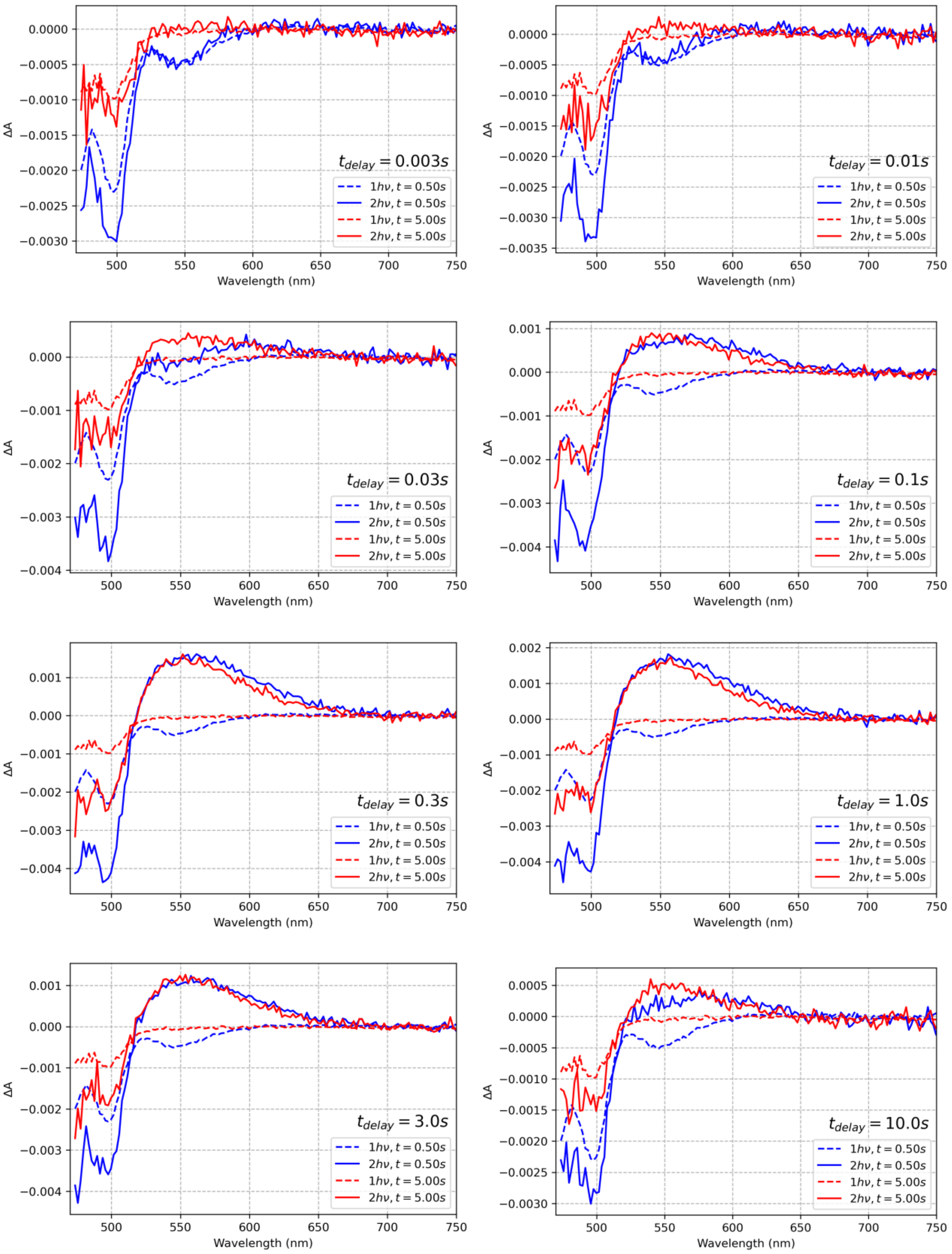

$t_{delay} = 0.003s$
$t_{delay} = 0.01s$
$t_{delay} = 0.03s$
$t_{delay} = 0.1s$
$t_{delay} = 0.3s$
$t_{delay} = 1.0s$
$t_{delay} = 3.0s$
$t_{delay} = 10.0s$
$1h\nu, t = 0.50s$
$2h\nu, t = 0.50s$
$1h\nu, t = 5.00s$
$2h\nu, t = 5.00s$
Wavelength (nm)
ΔA

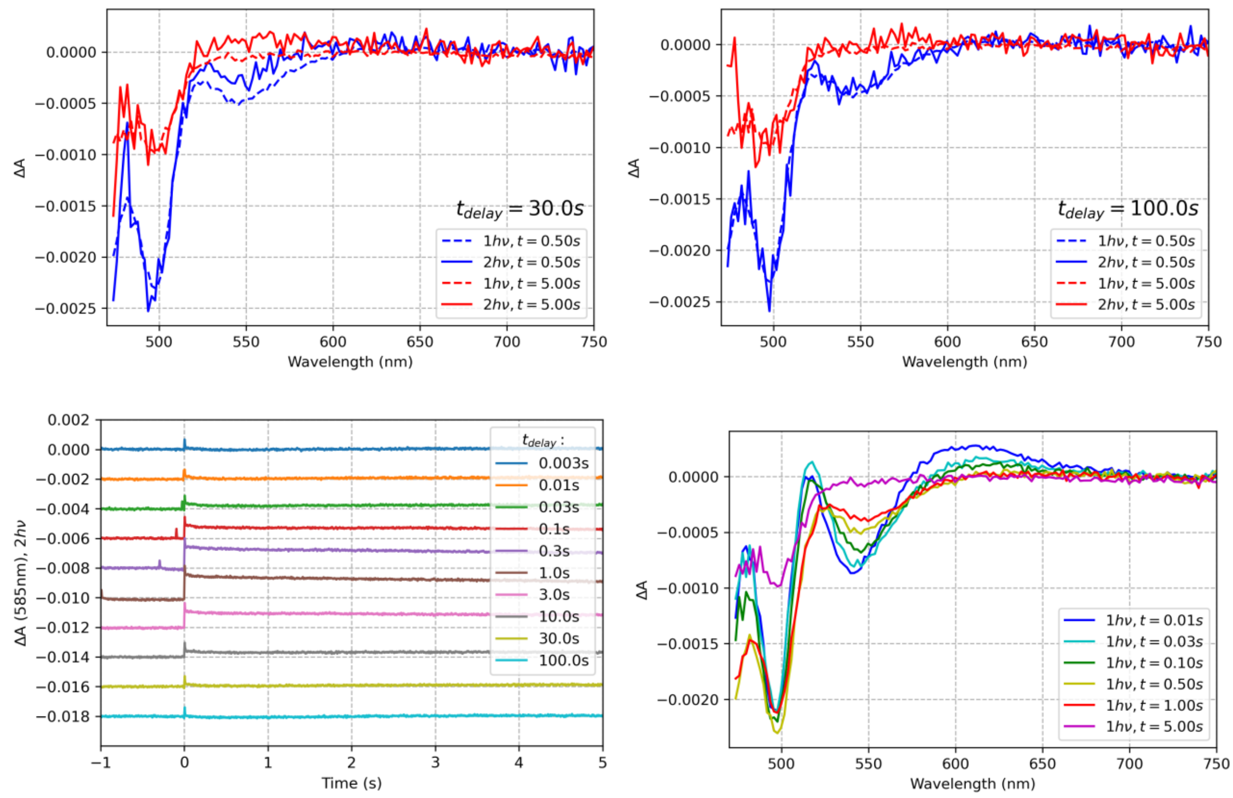


**Figure S9**. Two pulse experiment results obtained for OCP from *Synechocystis* functionalized with ECN, with His-tag removed. Excitation at 512 nm, energy density about 50 mJ/cm$^2$.

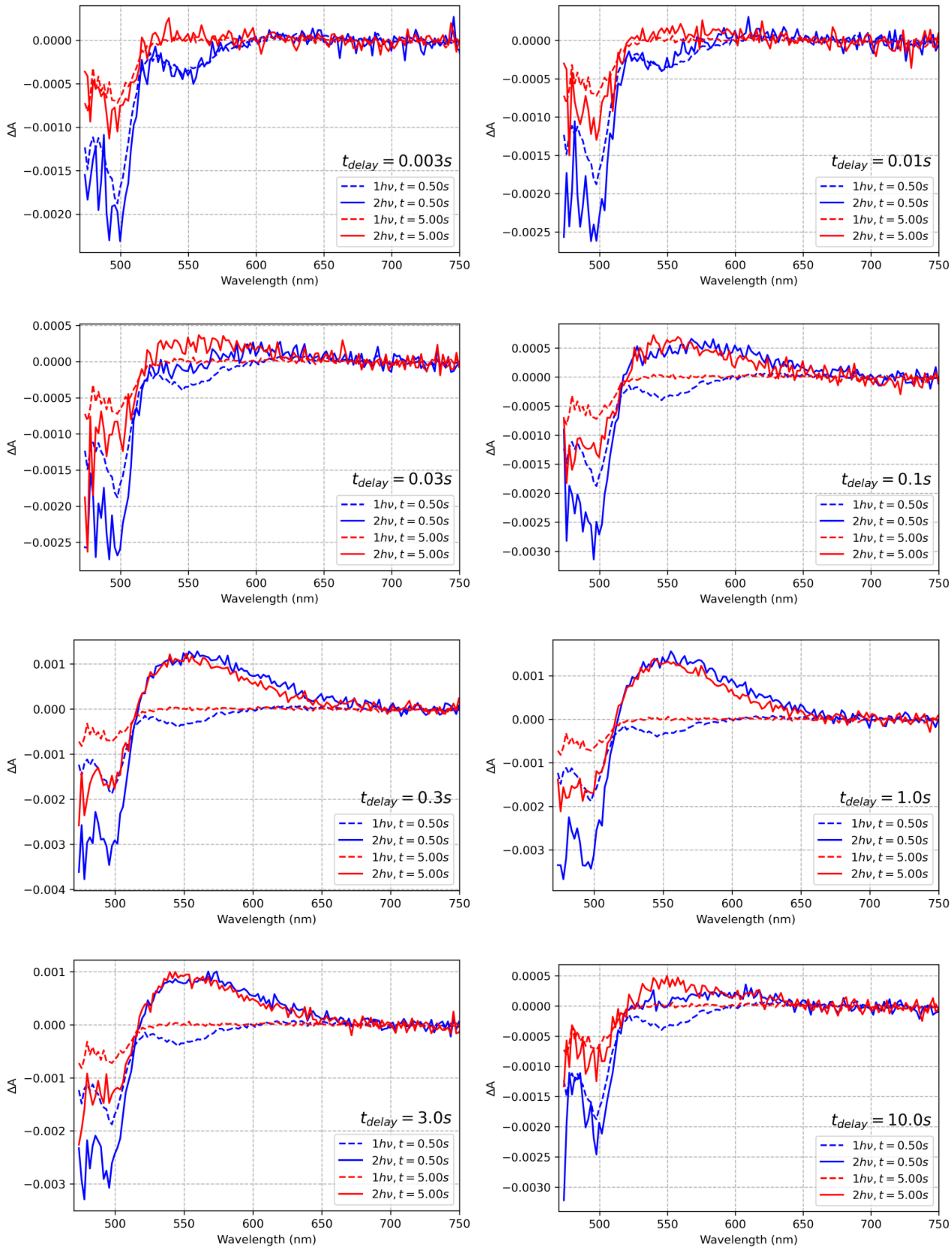
$t_{delay} = 0.003s$
$t_{delay} = 0.01s$
$t_{delay} = 0.03s$
$t_{delay} = 0.1s$
$t_{delay} = 0.3s$
$t_{delay} = 1.0s$
$t_{delay} = 3.0s$
$t_{delay} = 10.0s$
$1h\nu, t = 0.50s$
$2h\nu, t = 0.50s$
$1h\nu, t = 5.00s$
$2h\nu, t = 5.00s$
ΔA
Wavelength (nm)

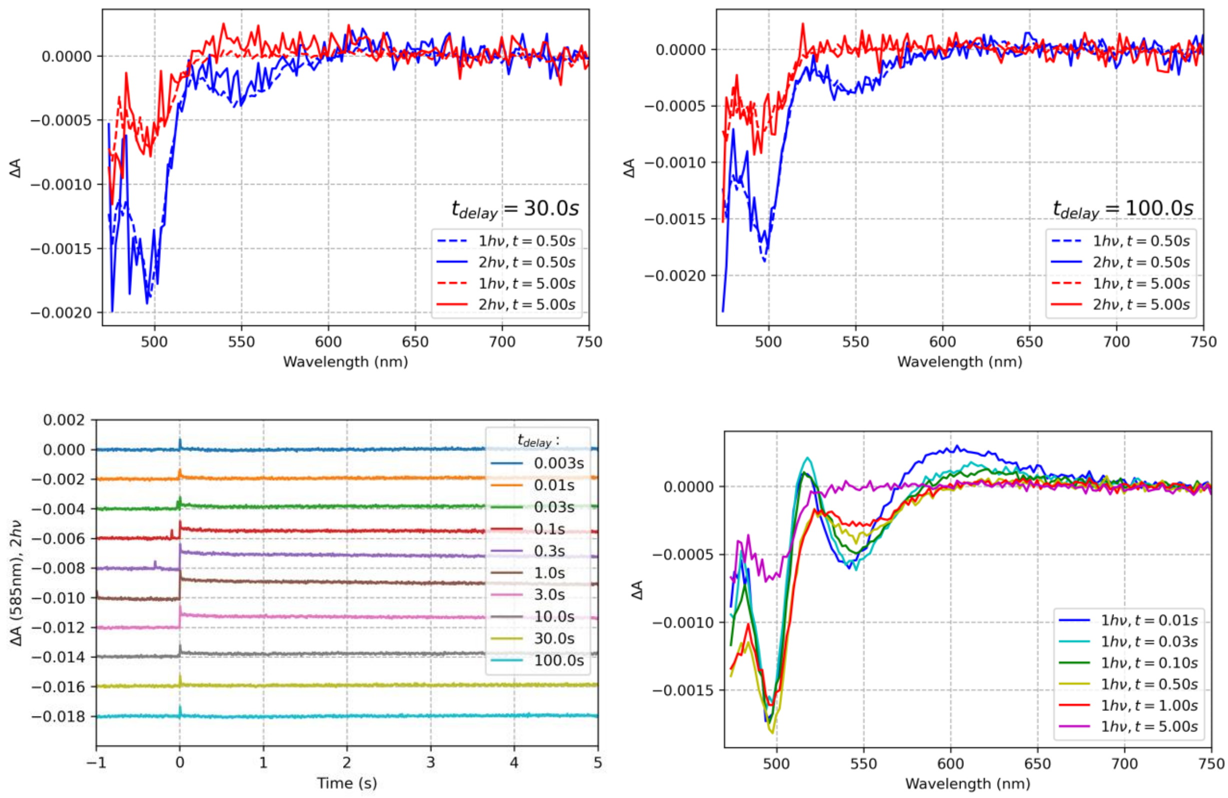


**Figure S10**. Two pulse experiment results obtained for OCP from *Planktothrix* functionalized with ECN, with His-tag removed. Excitation at 512 nm, energy density about 50 mJ/cm$^2$.

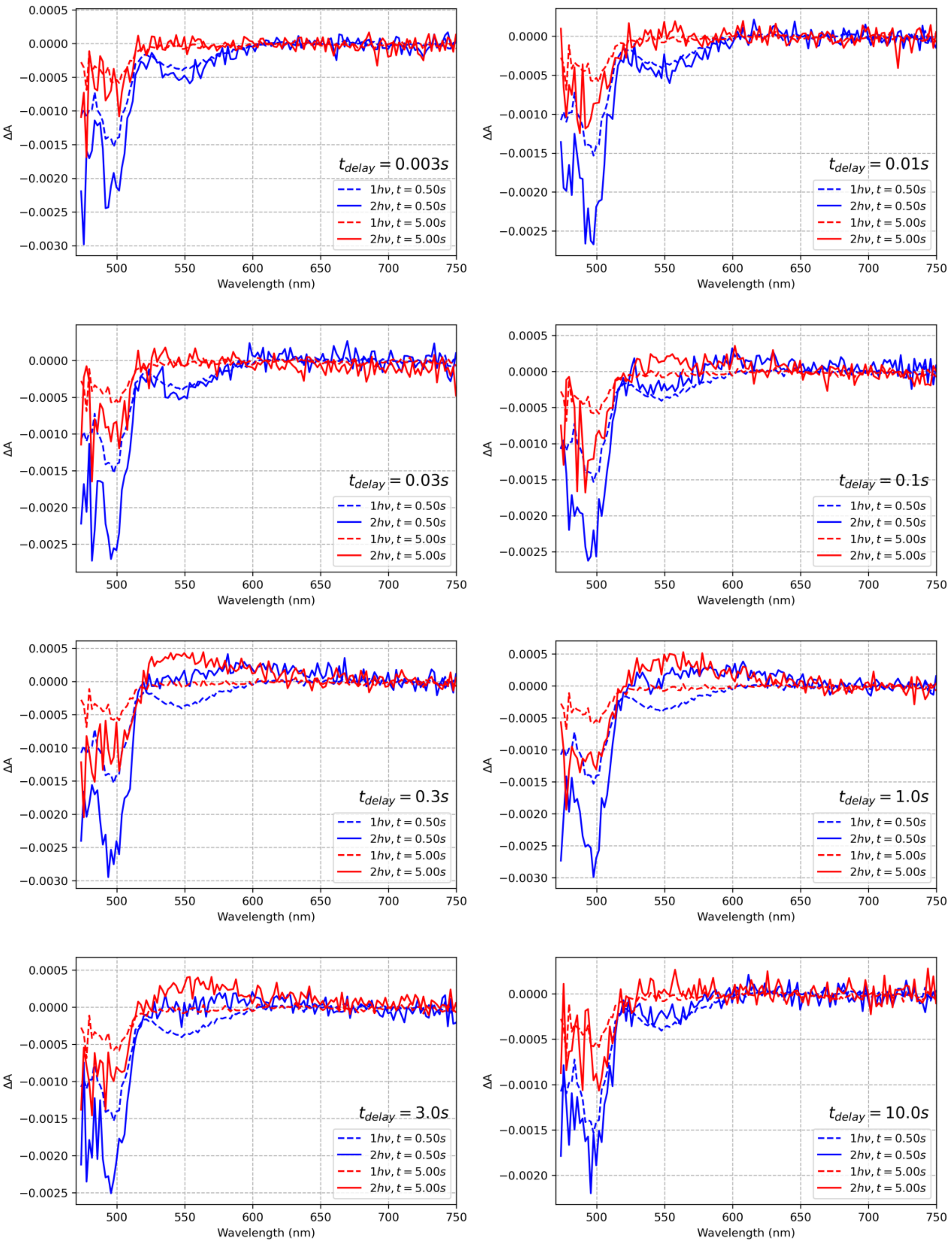
$t_{delay} = 0.003s$
$t_{delay} = 0.01s$
$t_{delay} = 0.03s$
$t_{delay} = 0.1s$
$t_{delay} = 0.3s$
$t_{delay} = 1.0s$
$t_{delay} = 3.0s$
$t_{delay} = 10.0s$
$1h\nu, t = 0.50s$
$2h\nu, t = 0.50s$
$1h\nu, t = 5.00s$
$2h\nu, t = 5.00s$
ΔA
Wavelength (nm)

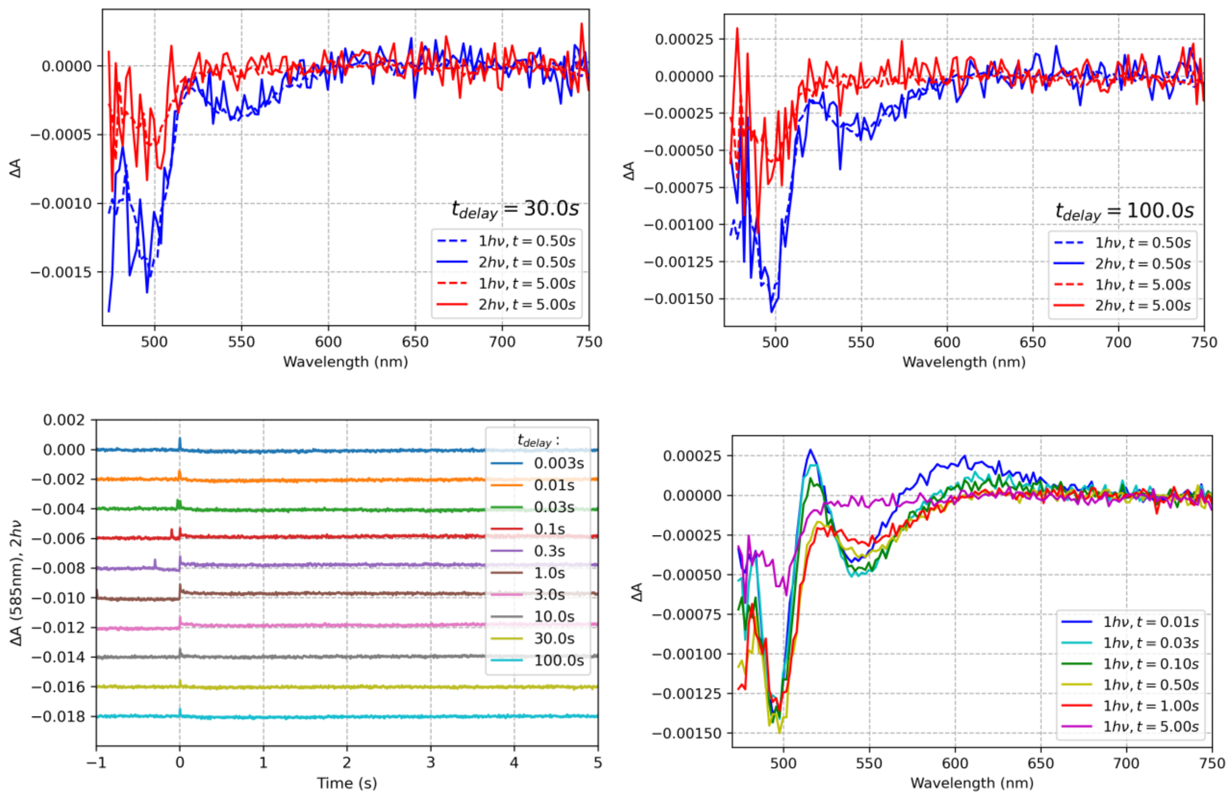


**Figure S11**. Two pulse experiment results obtained for OCP from *Planktothrix* functionalized with ECN, with His-tag removed. Excitation at 512 nm, energy density about 12 mJ/cm$^2$.

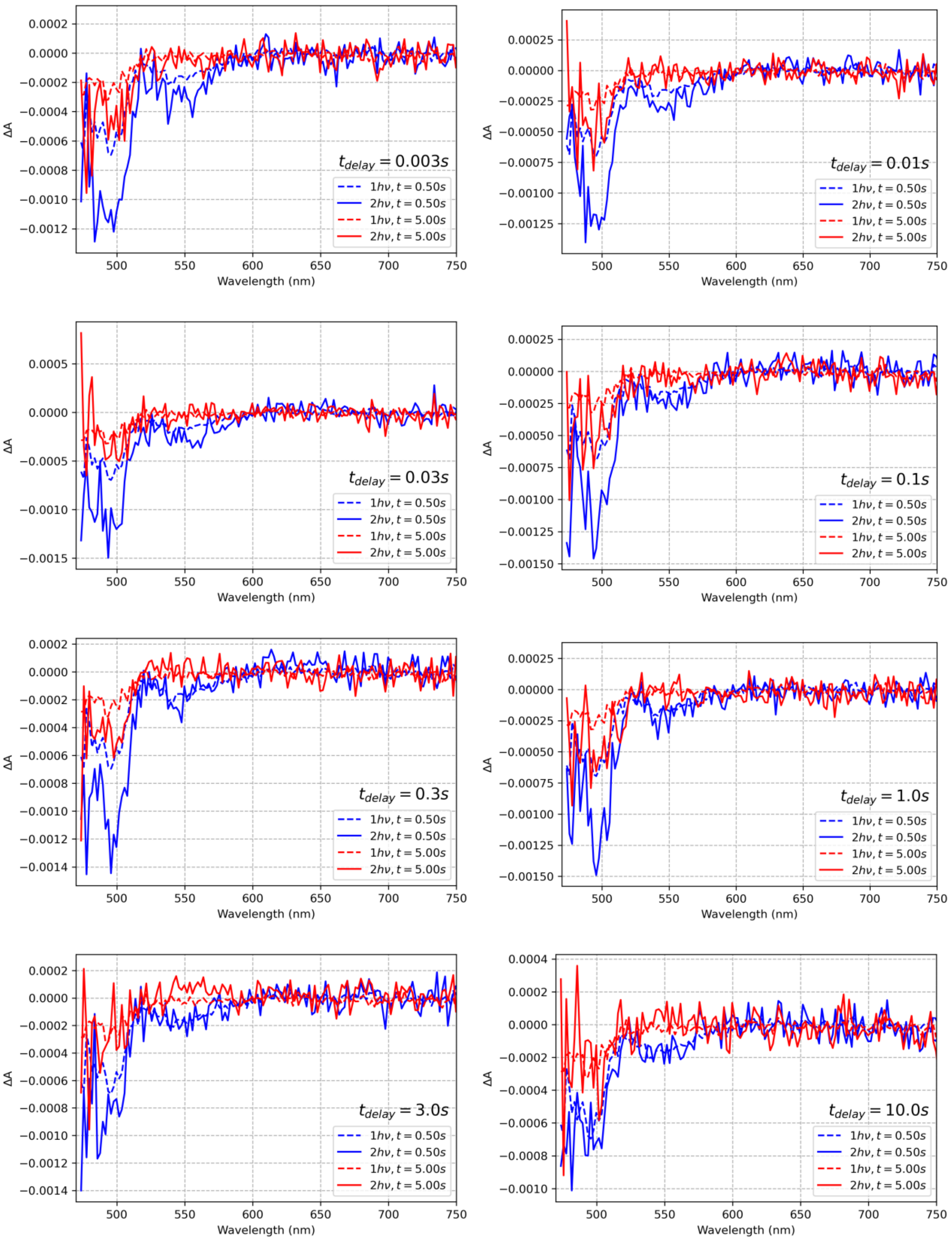

$t_{delay} = 0.003s$
$t_{delay} = 0.01s$
$t_{delay} = 0.03s$
$t_{delay} = 0.1s$
$t_{delay} = 0.3s$
$t_{delay} = 1.0s$
$t_{delay} = 3.0s$
$t_{delay} = 10.0s$
$1h\nu, t = 0.50s$
$2h\nu, t = 0.50s$
$1h\nu, t = 5.00s$
$2h\nu, t = 5.00s$
ΔA
Wavelength (nm)

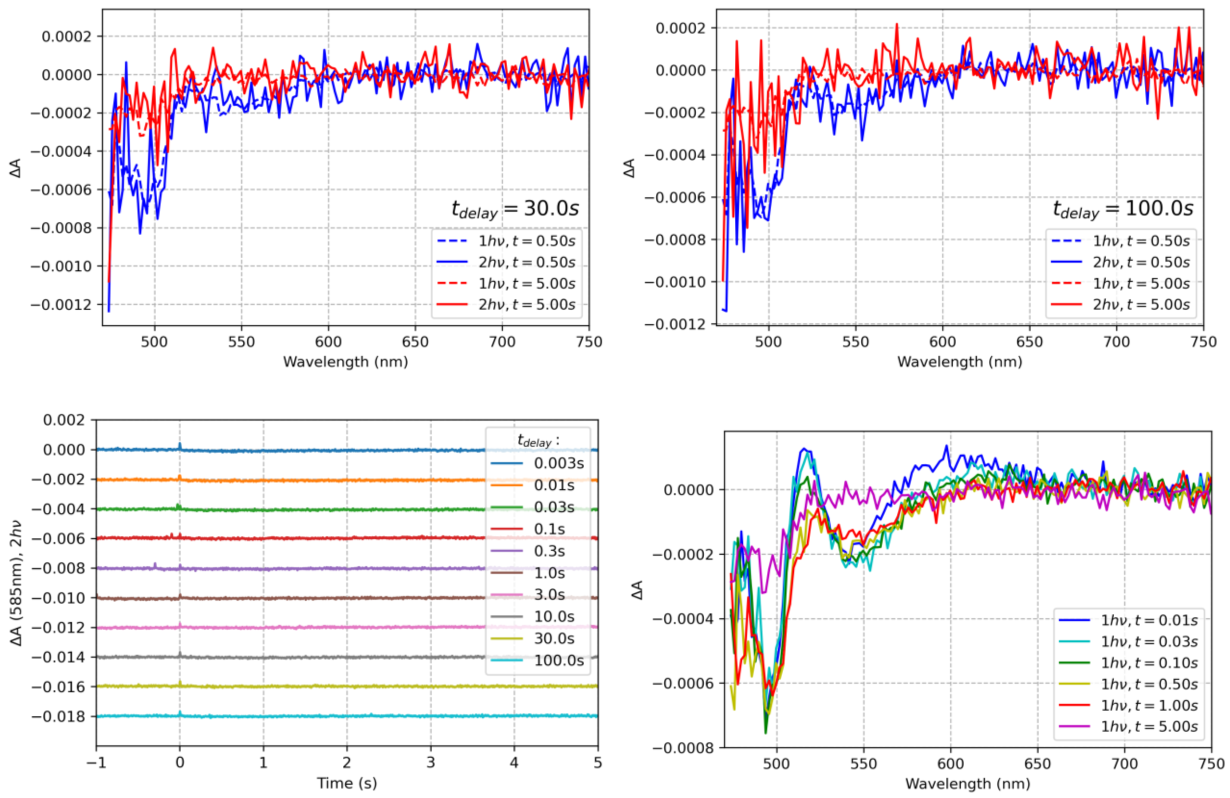


**Figure S12**. Two pulse experiment results obtained for OCP from *Planktothrix* functionalized with ECN, with His-tag removed. Excitation at 512 nm, energy density about 3 $mJ/cm^2$.

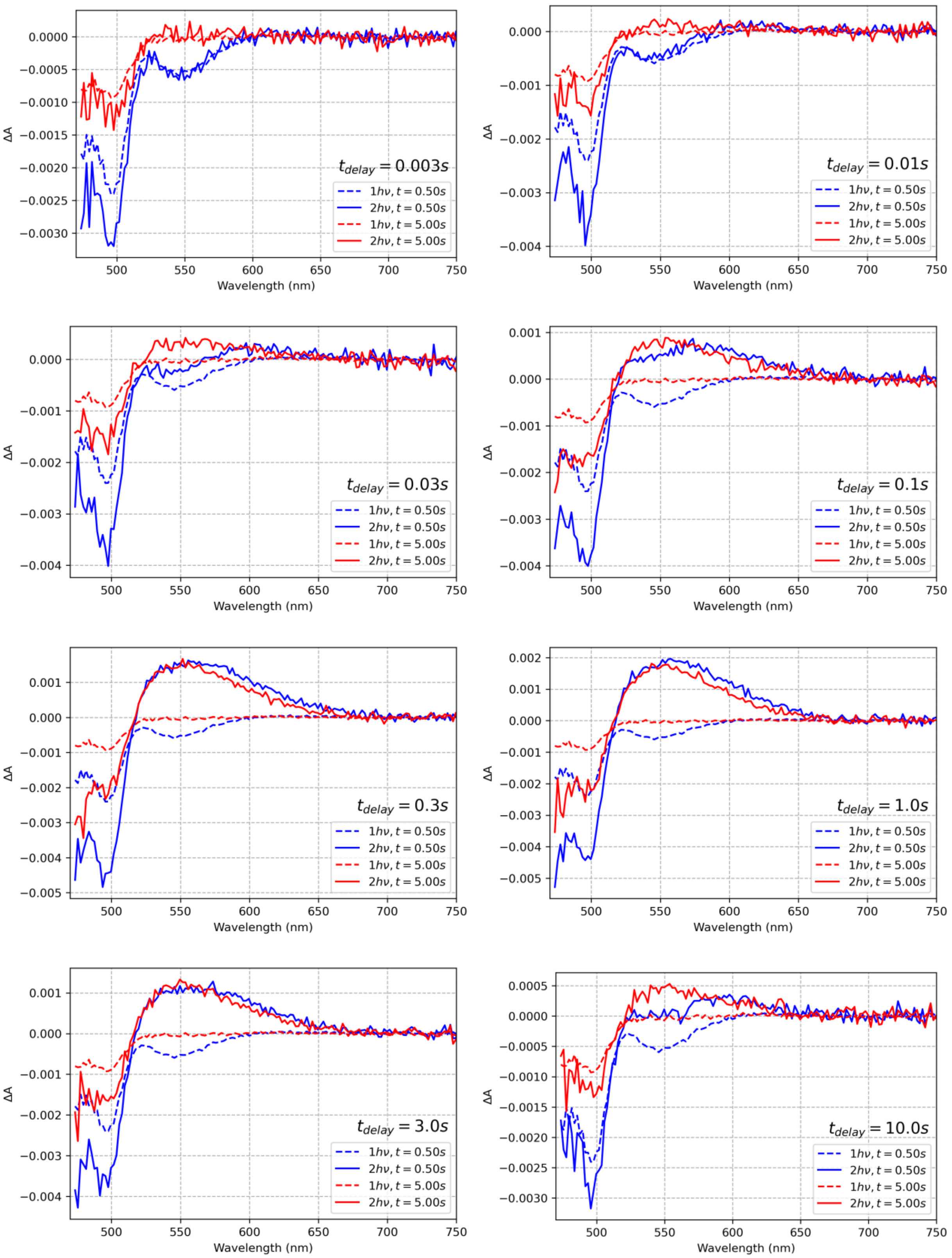
$t_{delay} = 0.003s$
$t_{delay} = 0.01s$
$t_{delay} = 0.03s$
$t_{delay} = 0.1s$
$t_{delay} = 0.3s$
$t_{delay} = 1.0s$
$t_{delay} = 3.0s$
$t_{delay} = 10.0s$
$1h\nu, t = 0.50s$
$2h\nu, t = 0.50s$
$1h\nu, t = 5.00s$
$2h\nu, t = 5.00s$
ΔA
Wavelength (nm)

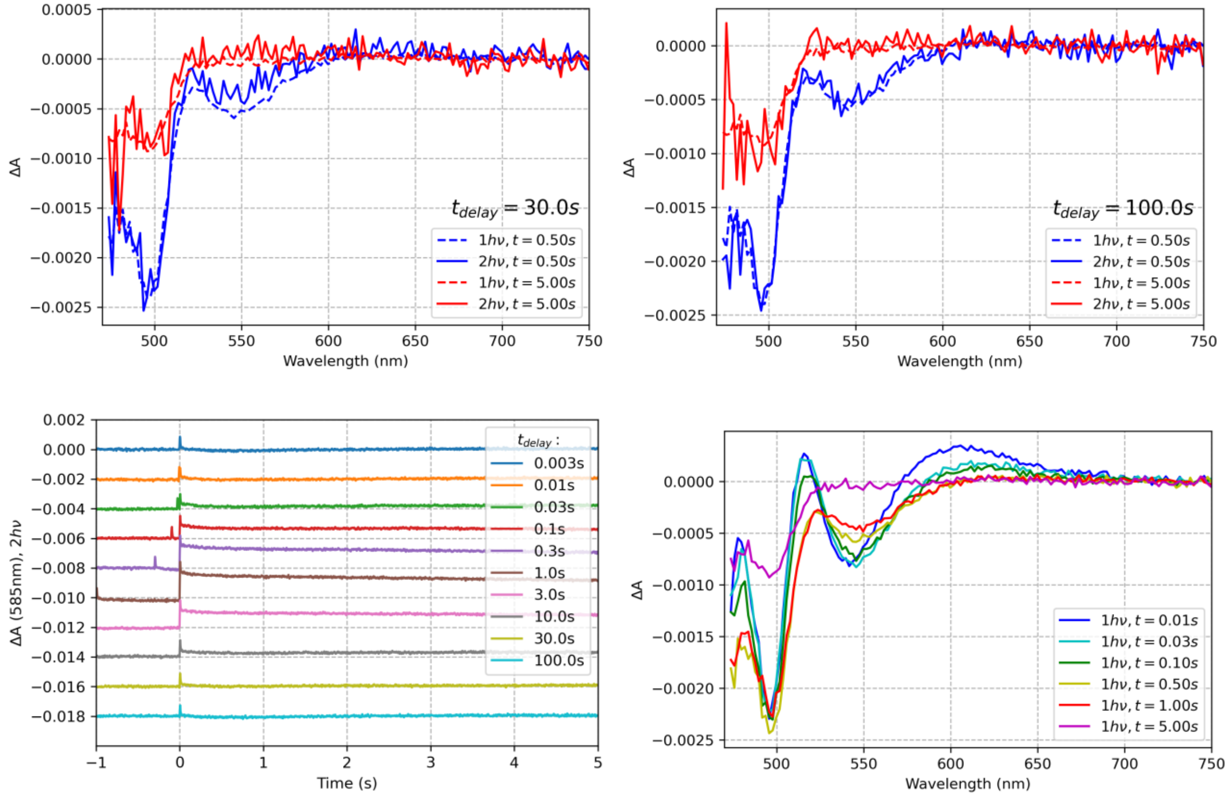


**Figure S13**. Two pulse experiment results obtained for OCP from *Planktothrix* functionalized with ECN, with His-tag removed. Excitation at 488 nm, energy density set very roughly to 50 mJ/cm$^2$.

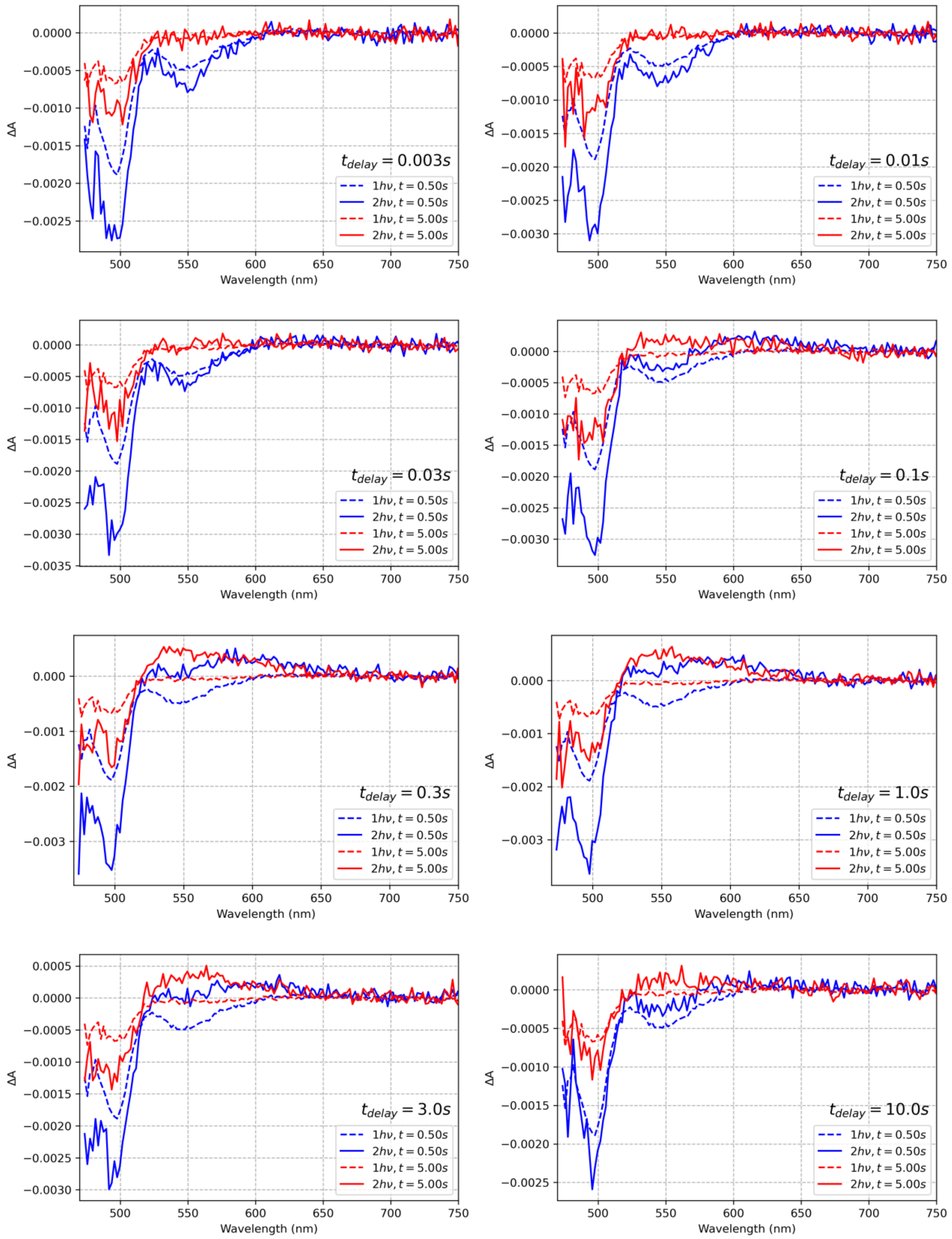

ΔA
Wavelength (nm)
t_delay = 0.003s
t_delay = 0.01s
t_delay = 0.03s
t_delay = 0.1s
t_delay = 0.3s
t_delay = 1.0s
t_delay = 3.0s
t_delay = 10.0s
1hν, t = 0.50s
2hν, t = 0.50s
1hν, t = 5.00s
2hν, t = 5.00s

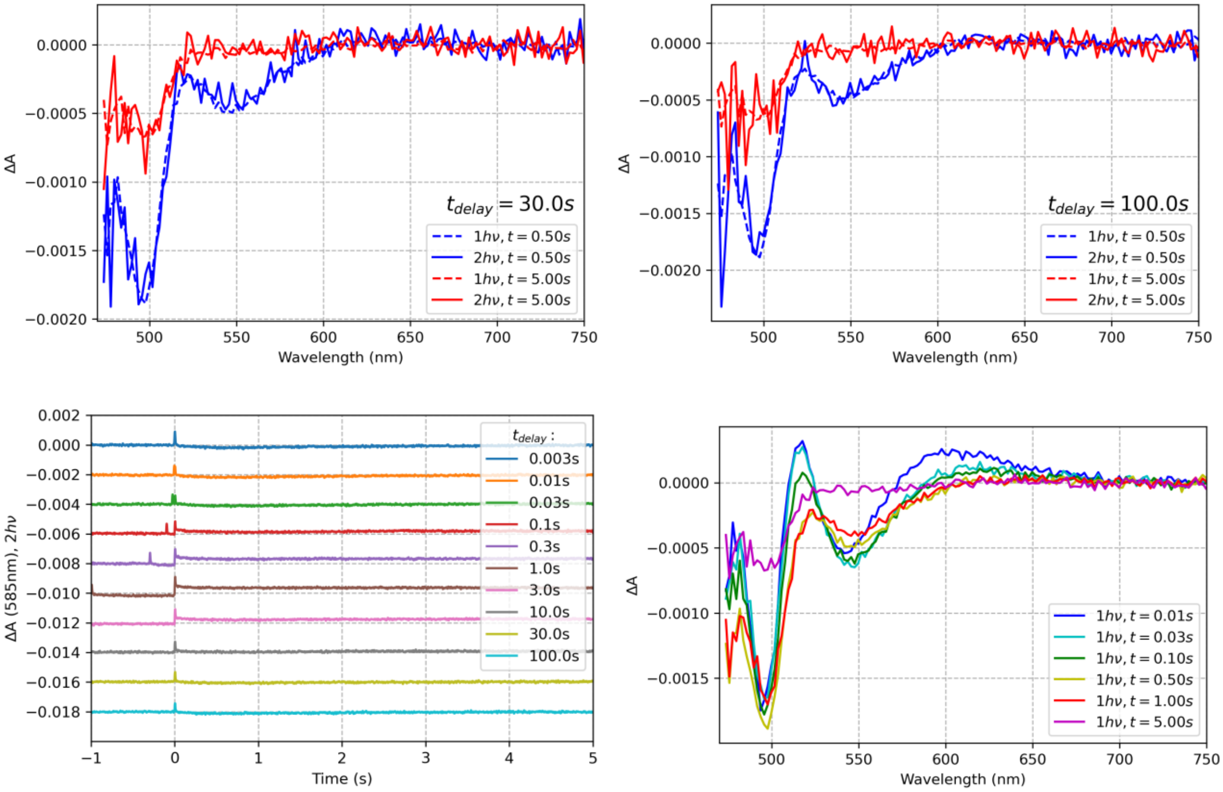


**Figure S14**. Two pulse experiment results obtained for OCP from *Planktothrix* functionalized with ECN, with His-tag removed. Excitation at 488 nm, energy density set very roughly 12 mJ/cm$^2$.

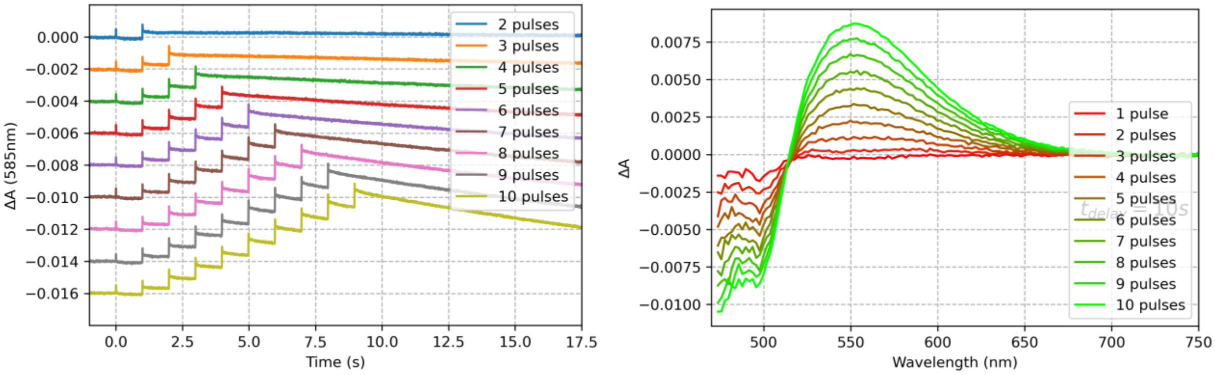


**Figure S15**. Photoconversion triggered by pulse trains comprising different number of pulses in the train. Experiment results obtained for OCP from *Planktothrix* functionalized with ECN, with His-tag removed. Excitation at 512 nm, pulse energy density about 12 mJ/cm$^2$.

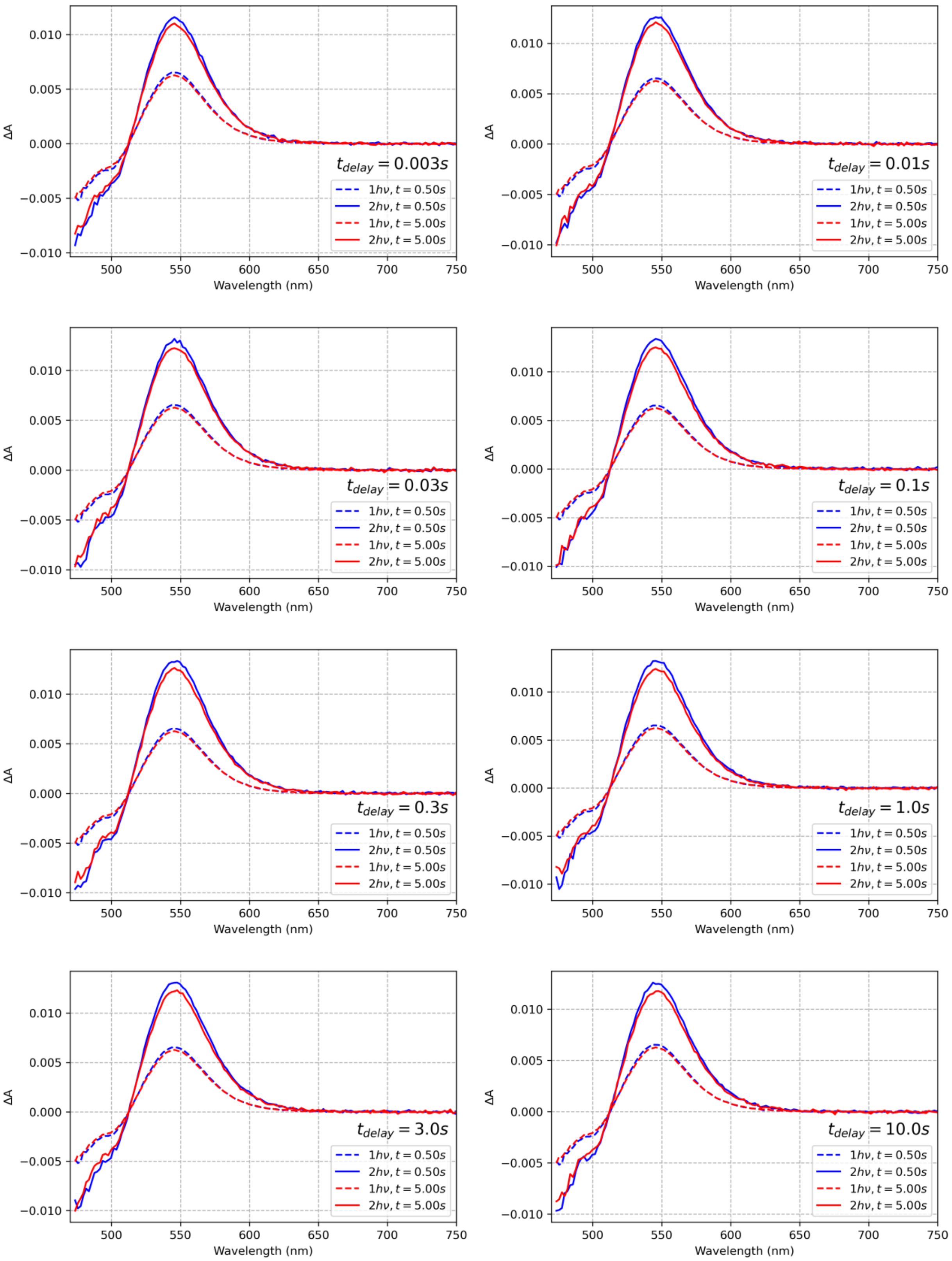
ΔA
Wavelength (nm)
$t_{delay} = 0.003s$
$t_{delay} = 0.01s$
$t_{delay} = 0.03s$
$t_{delay} = 0.1s$
$t_{delay} = 0.3s$
$t_{delay} = 1.0s$
$t_{delay} = 3.0s$
$t_{delay} = 10.0s$
$1h\nu, t = 0.50s$
$2h\nu, t = 0.50s$
$1h\nu, t = 5.00s$
$2h\nu, t = 5.00s$

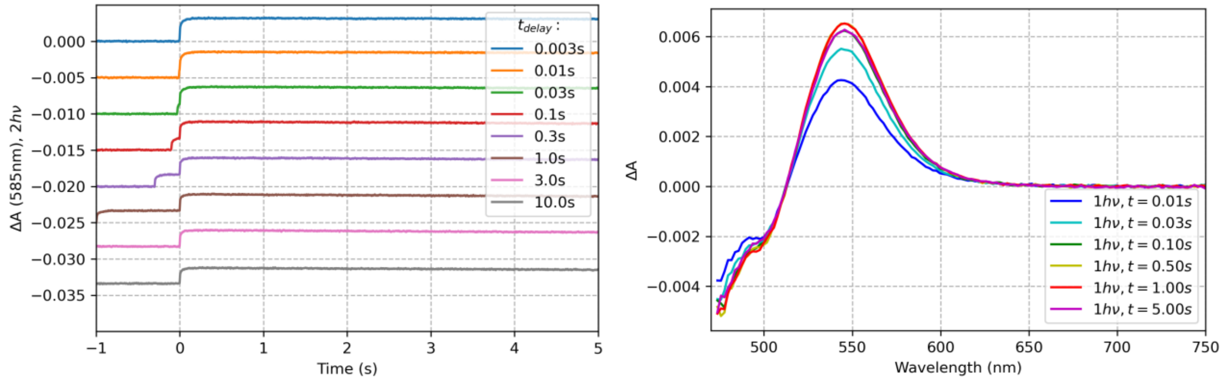


**Figure S16**. Two pulse experiment results obtained for OCP from *Synechocystis* functionalized with CAN, with His-tag removed. Excitation at 512 nm, energy density about 12 mJ/cm$^2$.

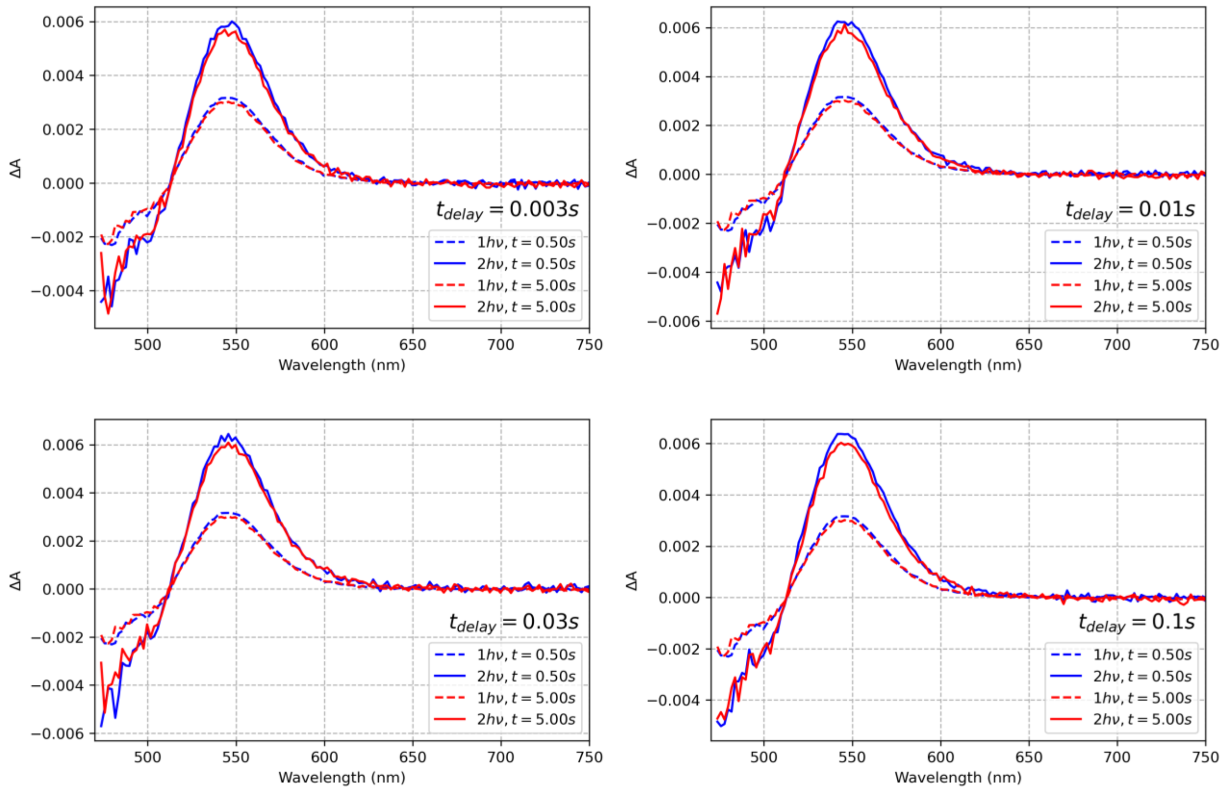

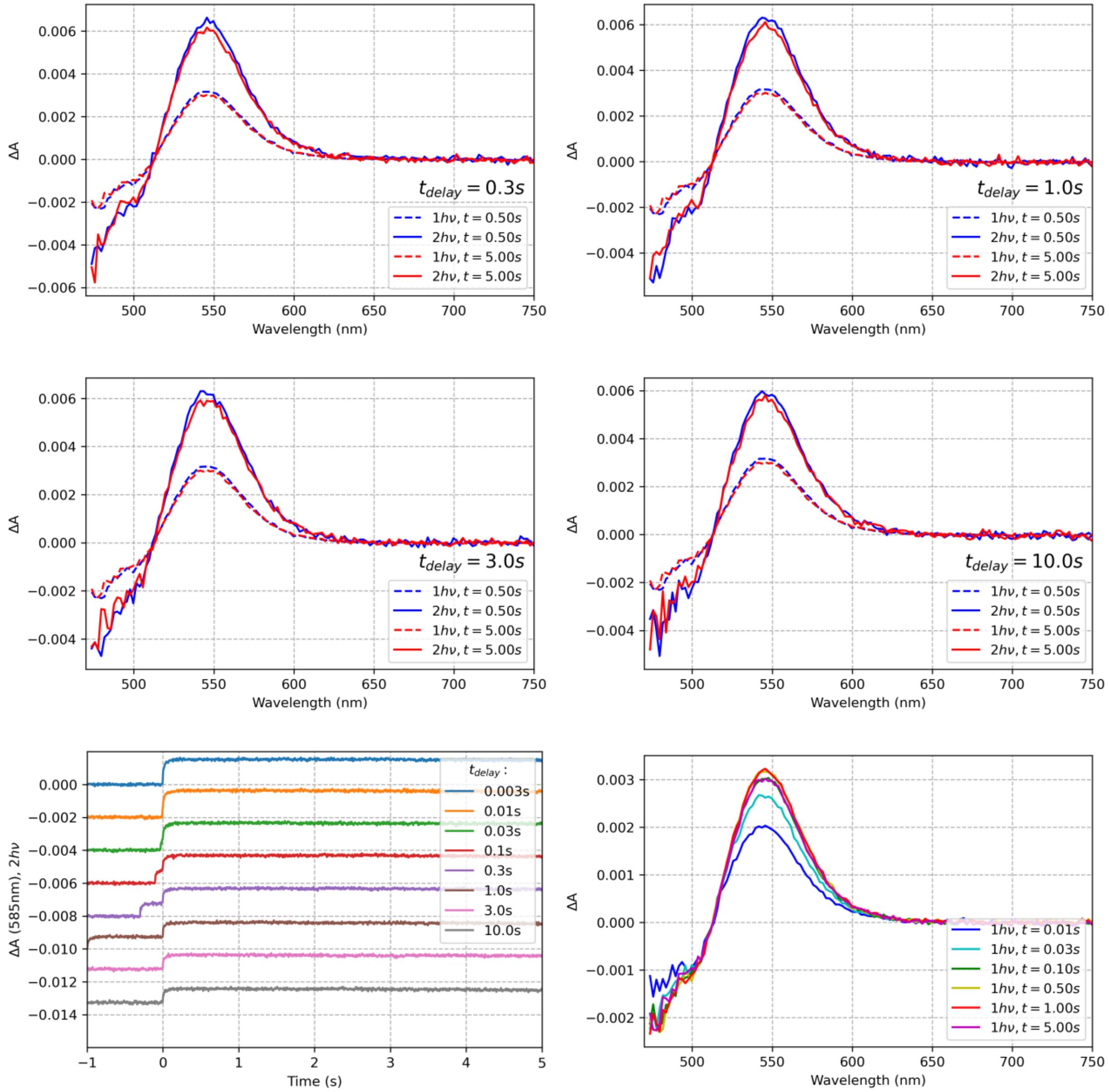

**Figure S17**. Two pulse experiment results obtained for OCP from *Synechocystis* functionalized with CAN, with His-tag removed. Excitation at 512 nm, energy density about 3 mJ/cm$^2$.

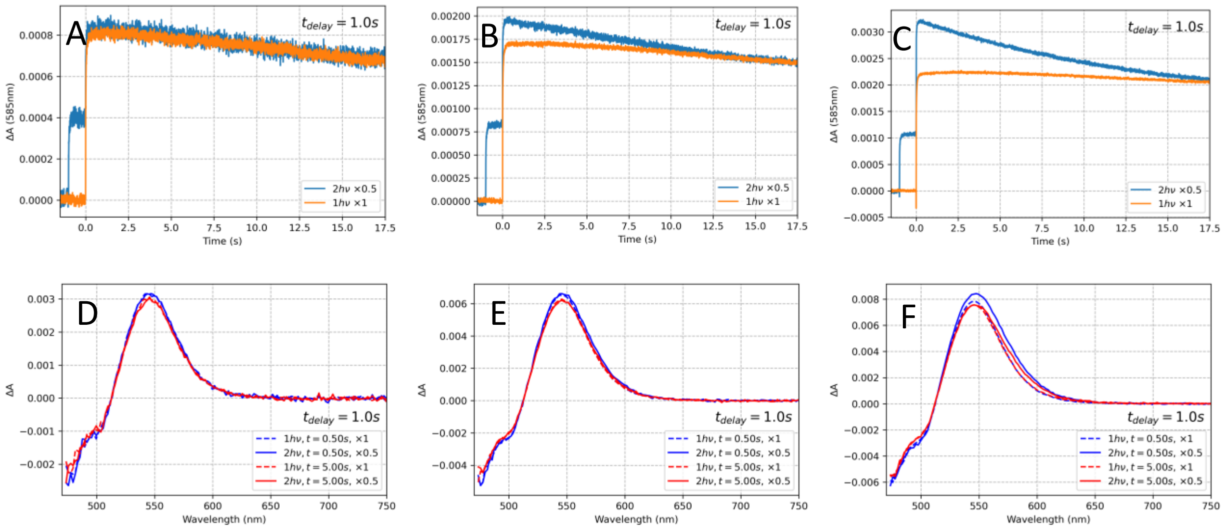


**Figure S18**. Two pulse experiment results obtained for OCP from *Synechocystis* functionalized with CAN, with His-tag removed. Excitation at 512 nm, energy density about 3 mJ/cm$^2$ (A, D), 12 mJ/cm$^2$ (B, E), 50 mJ/cm$^2$ (C, F). Spectra and kinetics are divided by the respective number of pulses used, $t_{delay}$ is 1 s.

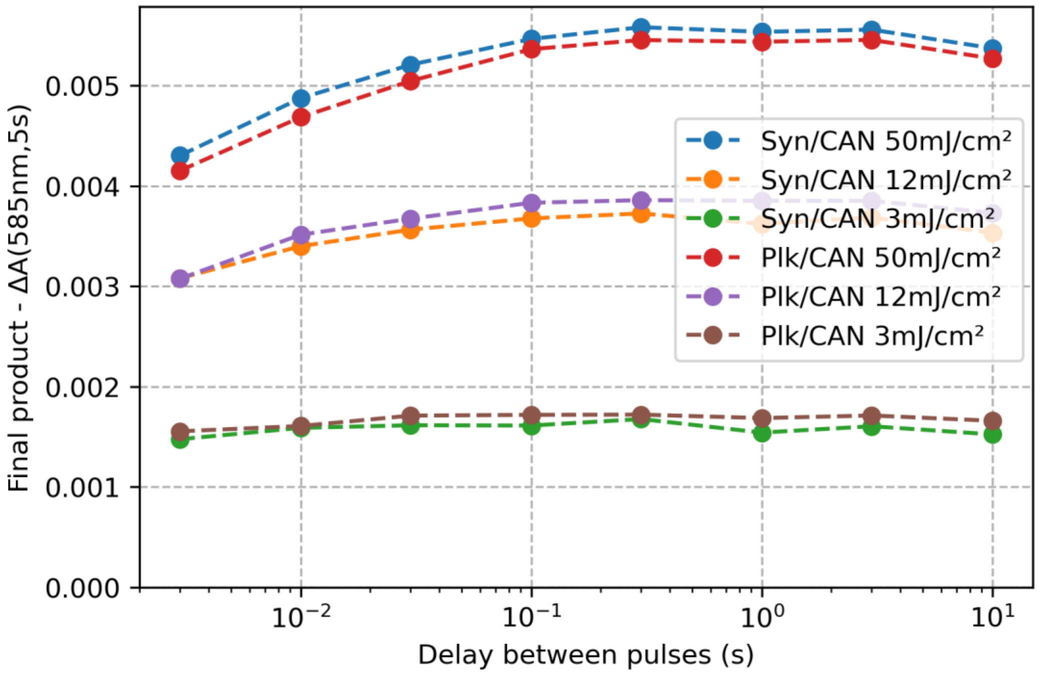


**Figure S19**. The change in absorption at 585 nm (5 s after the second pulse), representing the yield of OCP$^R$, plotted against $t_{delay}$ for CAN-functionalized OCPs and various excitation energy densities (512 nm).

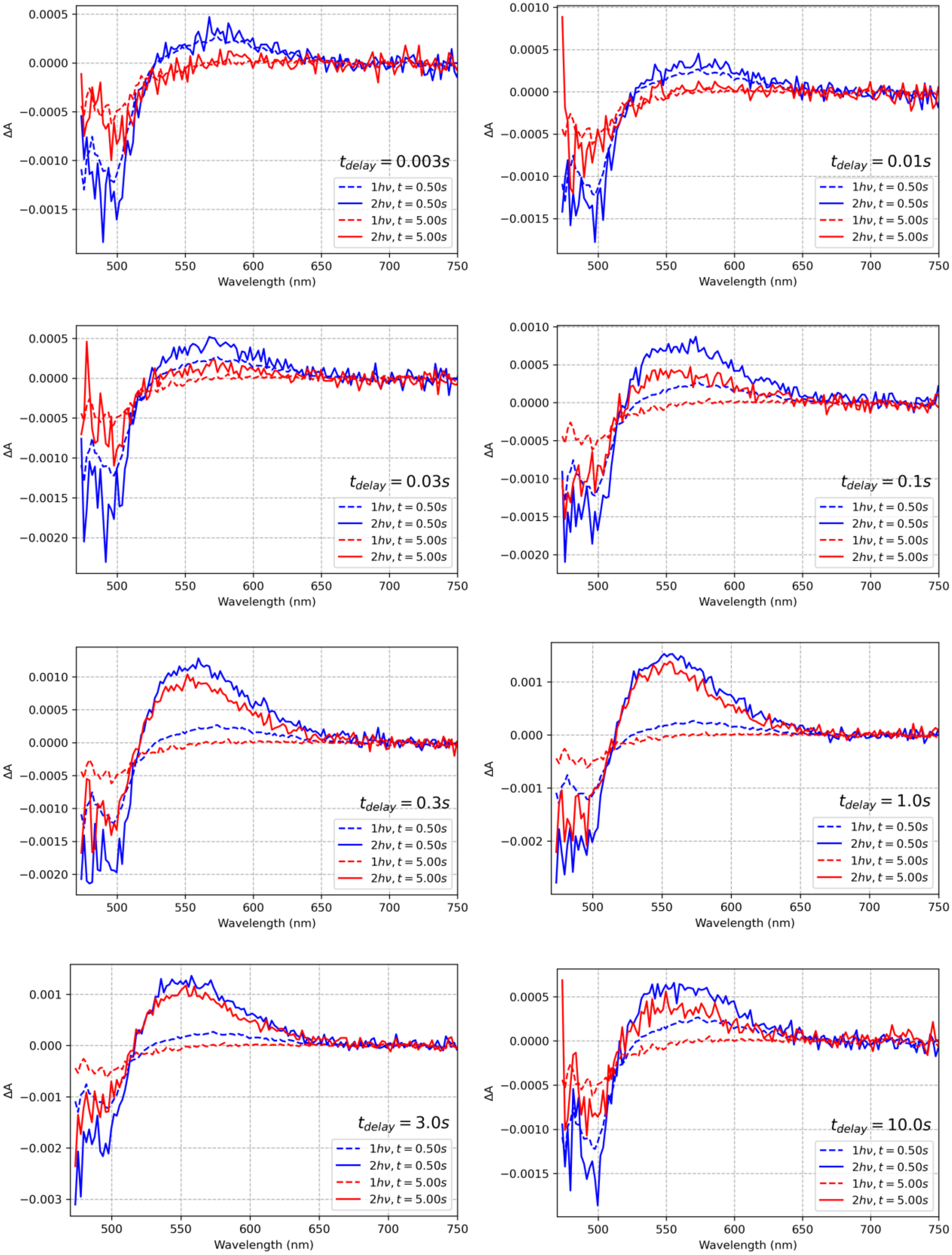
ΔA
Wavelength (nm)
$t_{delay} = 0.003s$
$t_{delay} = 0.01s$
$t_{delay} = 0.03s$
$t_{delay} = 0.1s$
$t_{delay} = 0.3s$
$t_{delay} = 1.0s$
$t_{delay} = 3.0s$
$t_{delay} = 10.0s$
$1h\nu, t = 0.50s$
$2h\nu, t = 0.50s$
$1h\nu, t = 5.00s$
$2h\nu, t = 5.00s$

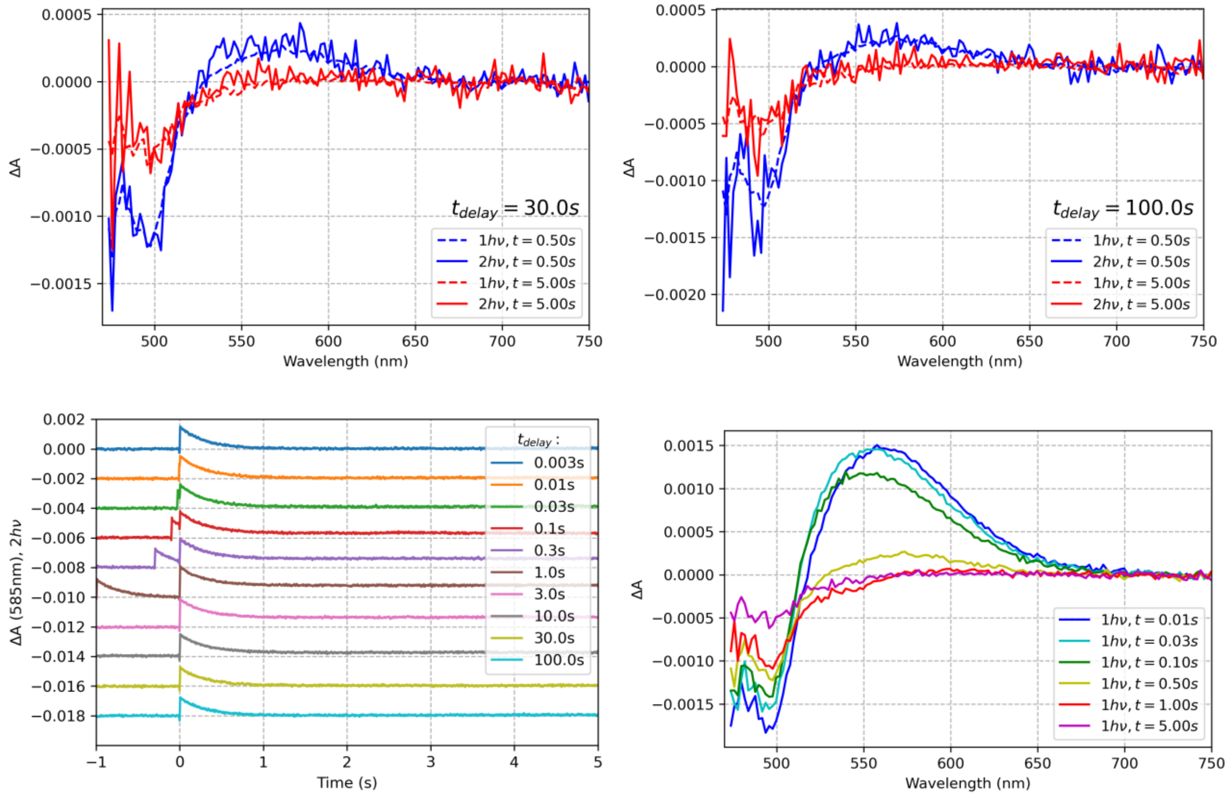


**Figure S20**. Two pulse experiment results obtained for OCP from *Synechocystis* functionalized with ECN with a His-tag at the N-terminus. Excitation at 512 nm, energy density about 50 mJ/cm$^2$.

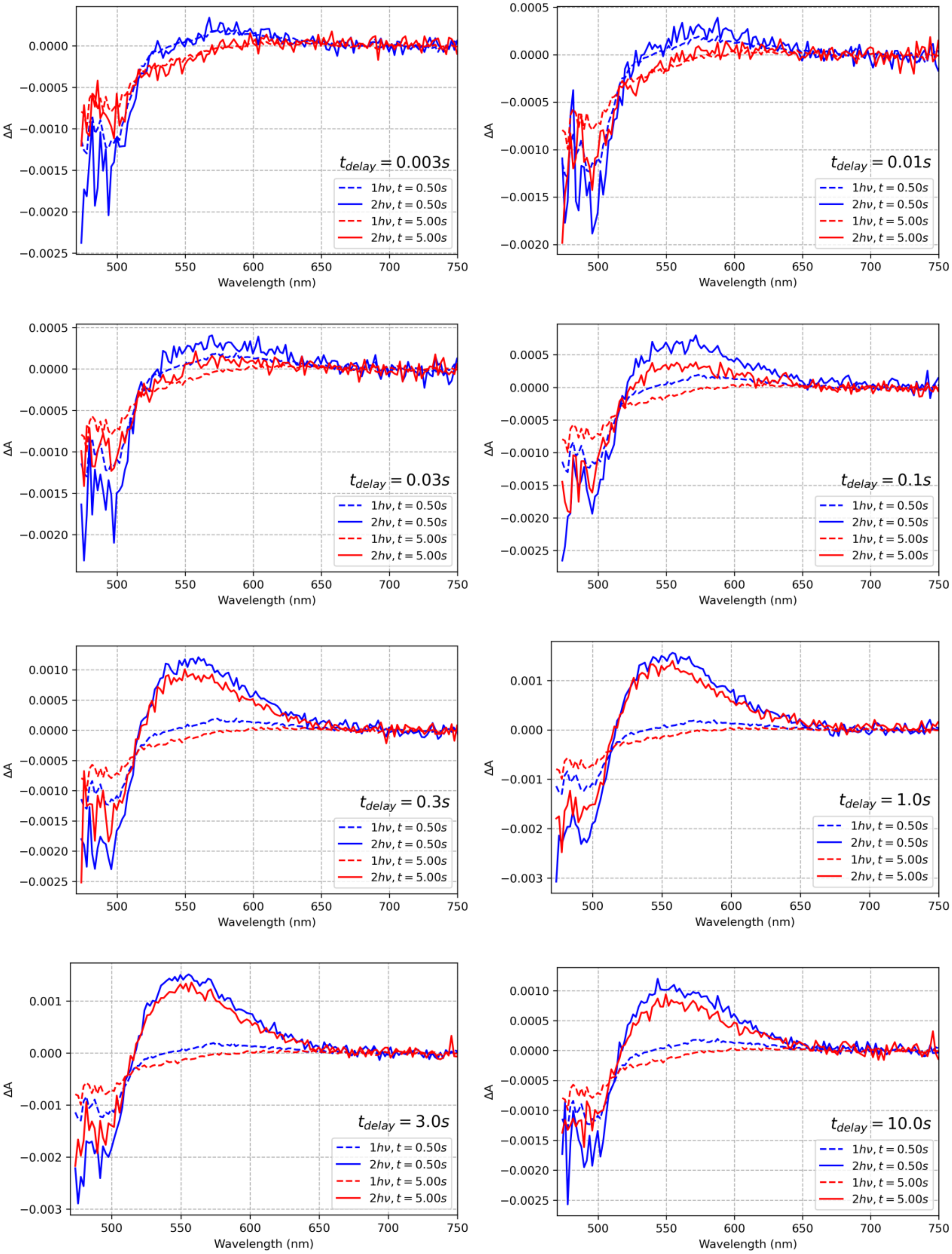

ΔA
Wavelength (nm)
t_delay = 0.003s
t_delay = 0.01s
t_delay = 0.03s
t_delay = 0.1s
t_delay = 0.3s
t_delay = 1.0s
t_delay = 3.0s
t_delay = 10.0s
1hν, t = 0.50s
2hν, t = 0.50s
1hν, t = 5.00s
2hν, t = 5.00s

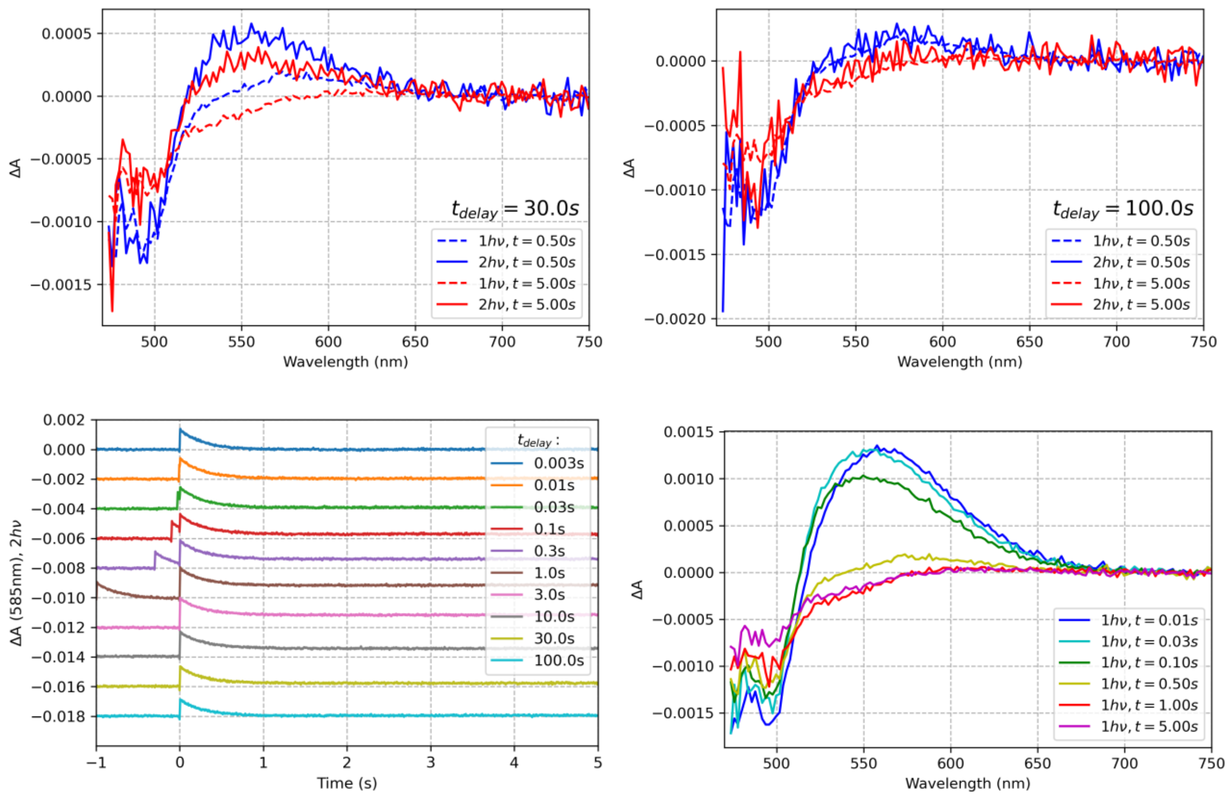


**Figure S21**. Two pulse experiment results obtained for OCP from *Synechocystis* functionalized with ECN with a His-tag at the C-terminus. Excitation at 512 nm, energy density about 50 mJ/cm$^2$.

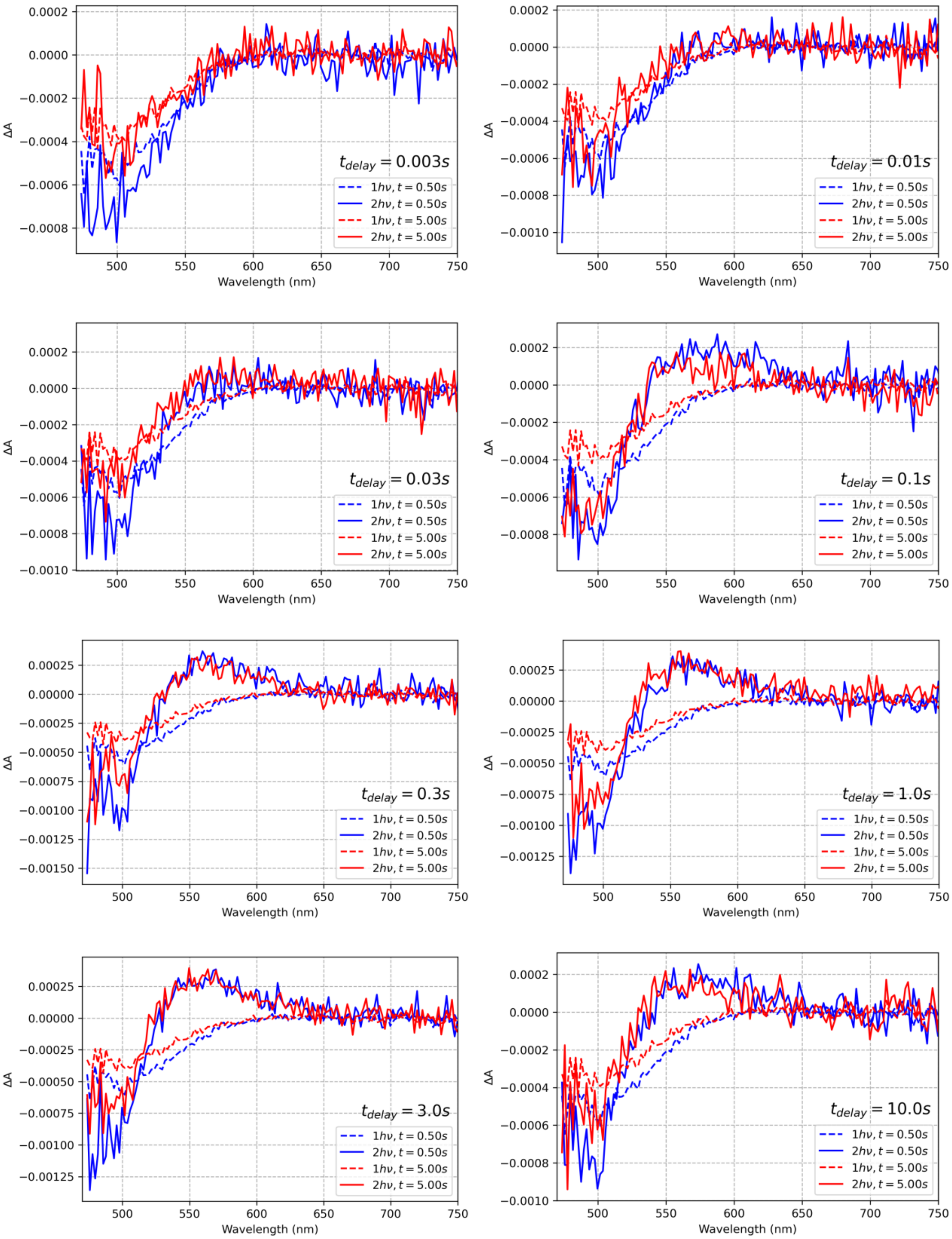
ΔA
Wavelength (nm)
$t_{delay} = 0.003s$
$t_{delay} = 0.01s$
$t_{delay} = 0.03s$
$t_{delay} = 0.1s$
$t_{delay} = 0.3s$
$t_{delay} = 1.0s$
$t_{delay} = 3.0s$
$t_{delay} = 10.0s$
$1h\nu, t = 0.50s$
$2h\nu, t = 0.50s$
$1h\nu, t = 5.00s$
$2h\nu, t = 5.00s$

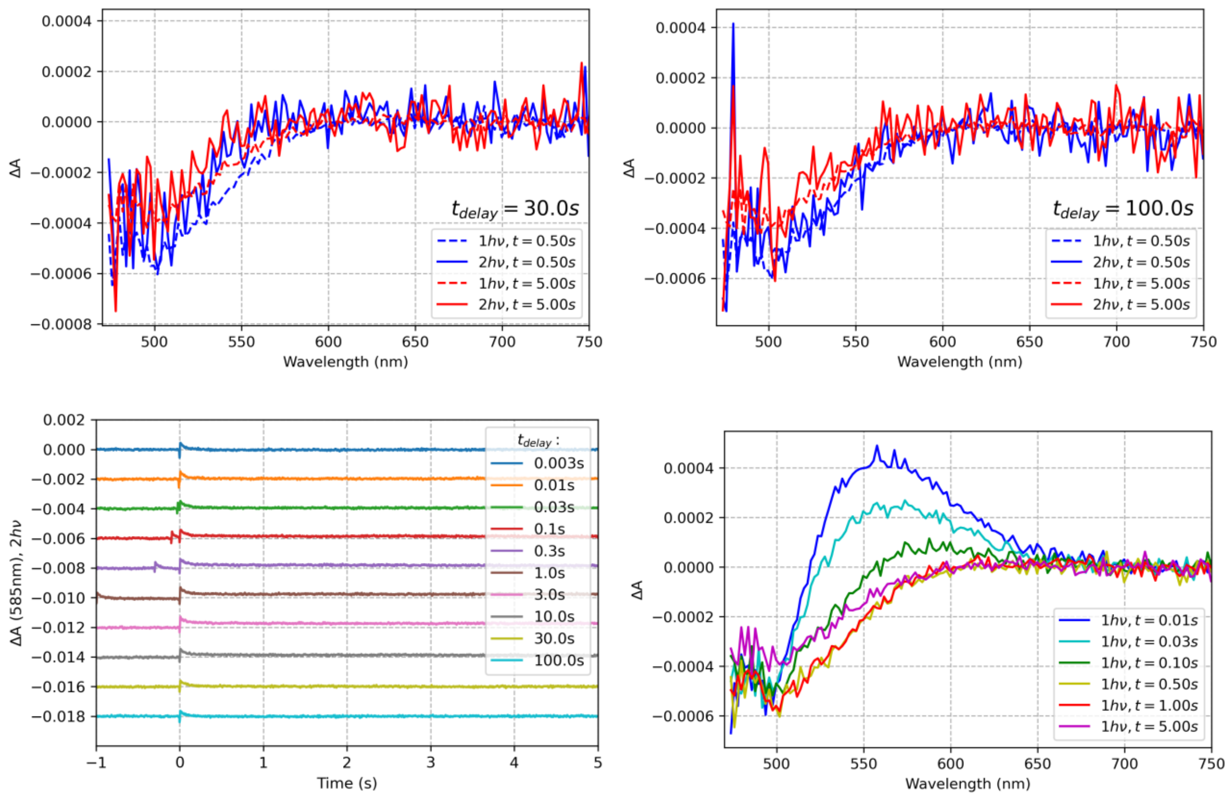


**Figure S22**. Two pulse experiment results obtained for the OCP L37V mutant from *Synechocystis* functionalized with ECN, with His-tag removed. Excitation at 512 nm, energy density about 50 mJ/cm$^2$.

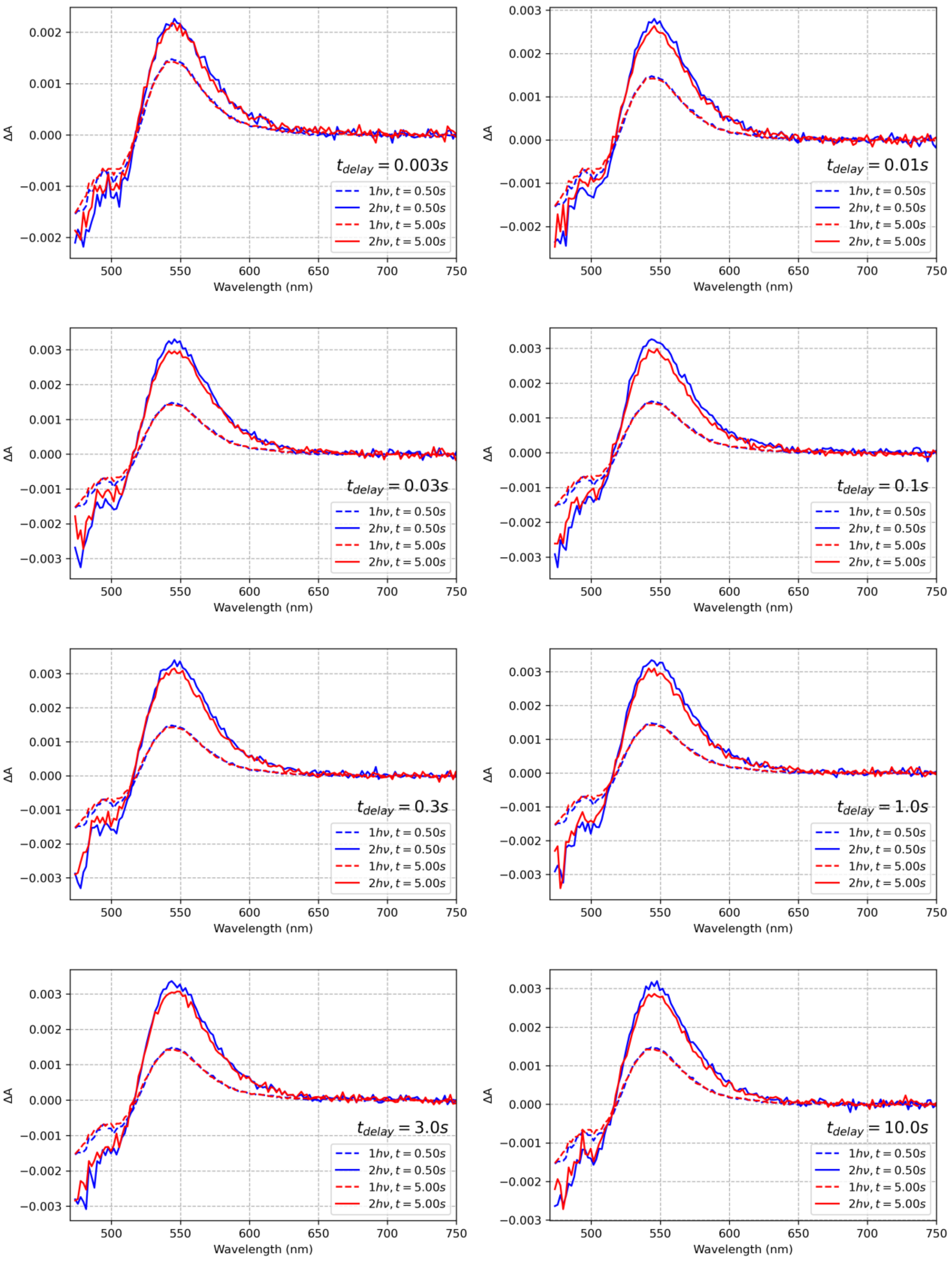
ΔA
Wavelength (nm)
t_delay = 0.003s
t_delay = 0.01s
t_delay = 0.03s
t_delay = 0.1s
t_delay = 0.3s
t_delay = 1.0s
t_delay = 3.0s
t_delay = 10.0s
1hν, t = 0.50s
2hν, t = 0.50s
1hν, t = 5.00s
2hν, t = 5.00s

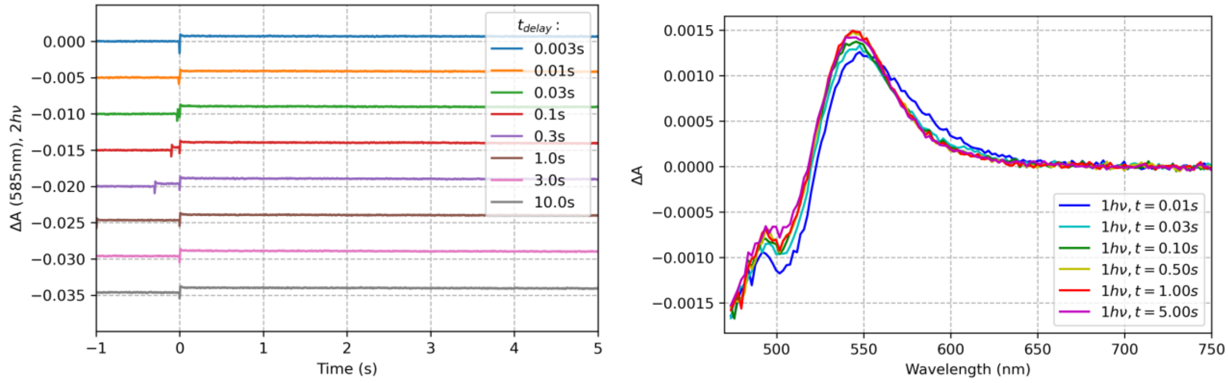


**Figure S23**. Two pulse experiment results obtained for the OCP mutant L37V from *Synechocystis* functionalized with CAN, with His-tag removed. Excitation at 512 nm, energy density about 50 mJ/cm$^2$.

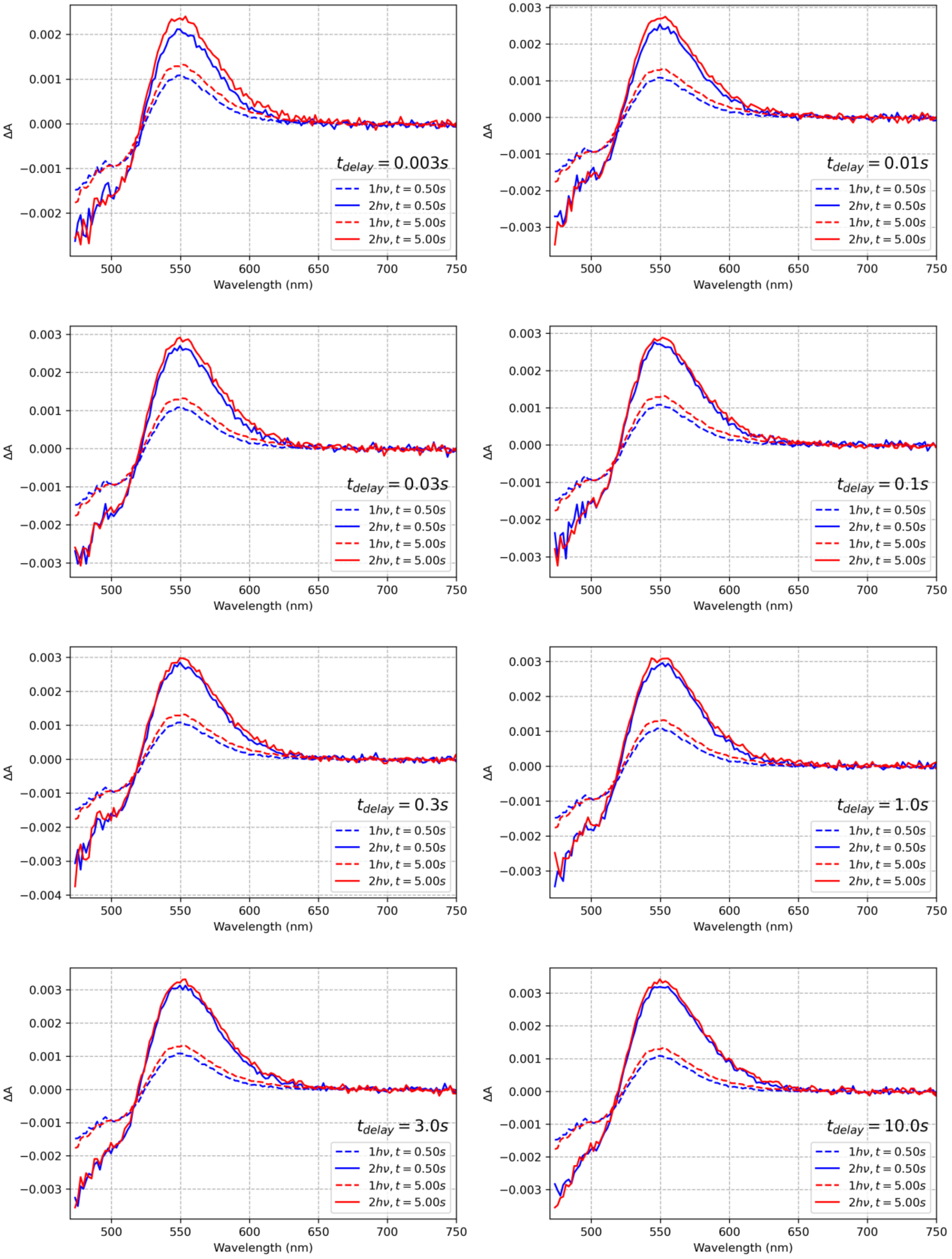

ΔA
Wavelength (nm)
500
550
600
650
700
750
0.003
0.002
0.001
0.000
−0.001
−0.002
−0.003
−0.004
$t_{delay} = 0.003s$
$t_{delay} = 0.01s$
$t_{delay} = 0.03s$
$t_{delay} = 0.1s$
$t_{delay} = 0.3s$
$t_{delay} = 1.0s$
$t_{delay} = 3.0s$
$t_{delay} = 10.0s$
$1h\nu, t = 0.50s$
$2h\nu, t = 0.50s$
$1h\nu, t = 5.00s$
$2h\nu, t = 5.00s$

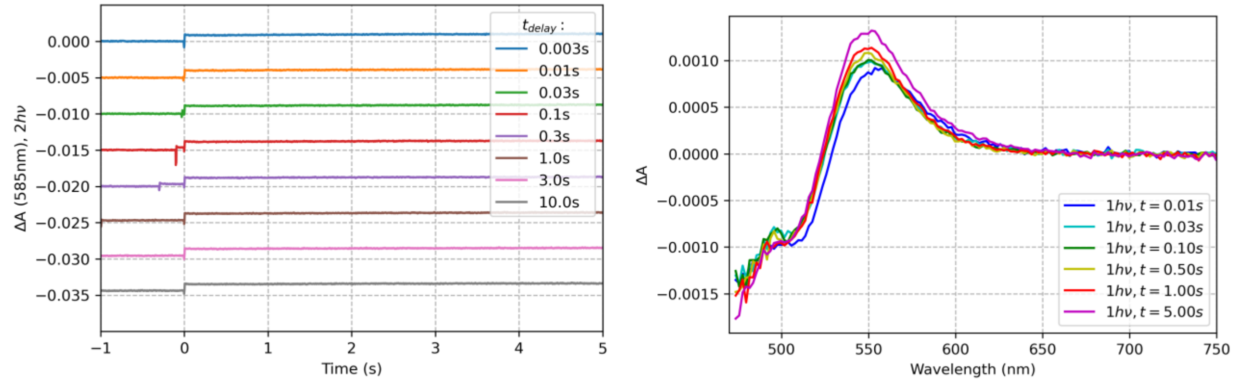


**Figure S24**. Two pulse experiment results obtained for the OCP mutant L37A from *Synechocystis* functionalized with CAN, with His-tag removed. Excitation at 512 nm, energy density about 50 mJ/cm$^2$.

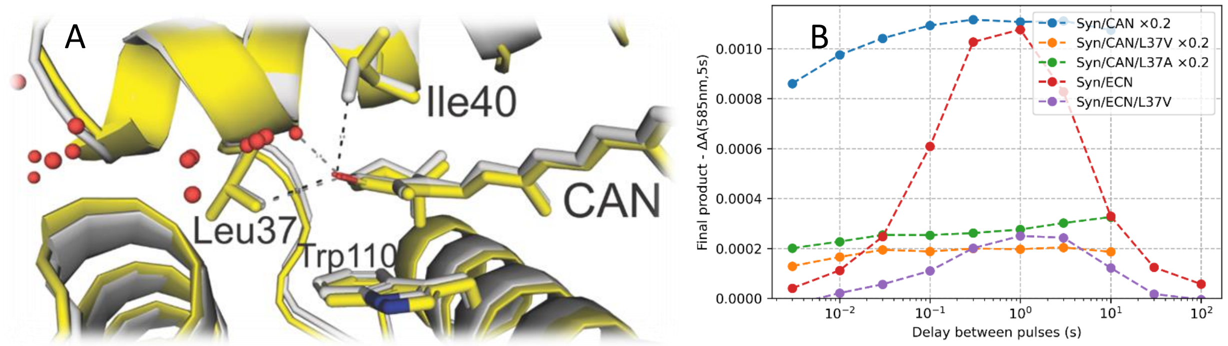


**Figure S25**. Environment of the carotenoid β2 ionone ring. A) Overlay of structures of CAN complexed OCP. In the *Synechocystis* variant (PDB 7ZSF, yellow, similar for PDB 4XB5)) the methyl group of Ile40 points away from the CAN keto-group, making room for a water molecule (which was not modeled in the structure, distance 2.6 Å). In the *Planktotrix* variant (PDB 7QD2, grey), the methyl group of Ile40 points towards the CAN keto-group (distance 3.6 Å), preventing binding of a water molecule. The distance between the keto-group and the methyl group of Leu37 is ~3.5 Å in $OCP_{Syn}$ and 3.2 Å in $OCP_{Plk}$. Water molecules are shown as red spheres. B) The change in absorption at 585 nm (5 s after the second pulse), representing the yield of $OCP^R$, plotted against $t_{delay}$. Excitation at 512 nm, energy density about 50 mJ/cm$^2$.

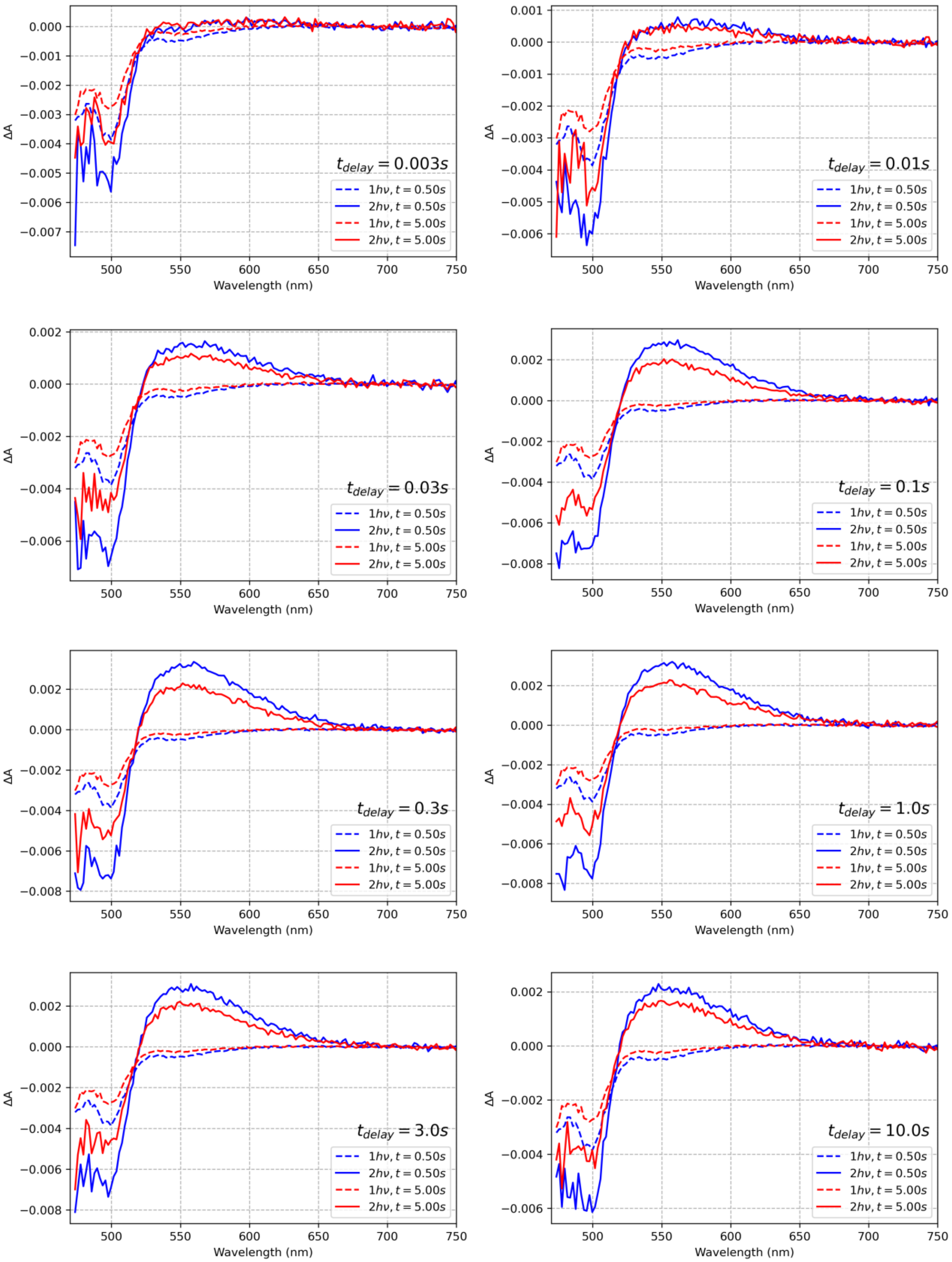
ΔA
Wavelength (nm)
$t_{delay} = 0.003s$
$t_{delay} = 0.01s$
$t_{delay} = 0.03s$
$t_{delay} = 0.1s$
$t_{delay} = 0.3s$
$t_{delay} = 1.0s$
$t_{delay} = 3.0s$
$t_{delay} = 10.0s$
$1h\nu, t = 0.50s$
$2h\nu, t = 0.50s$
$1h\nu, t = 5.00s$
$2h\nu, t = 5.00s$

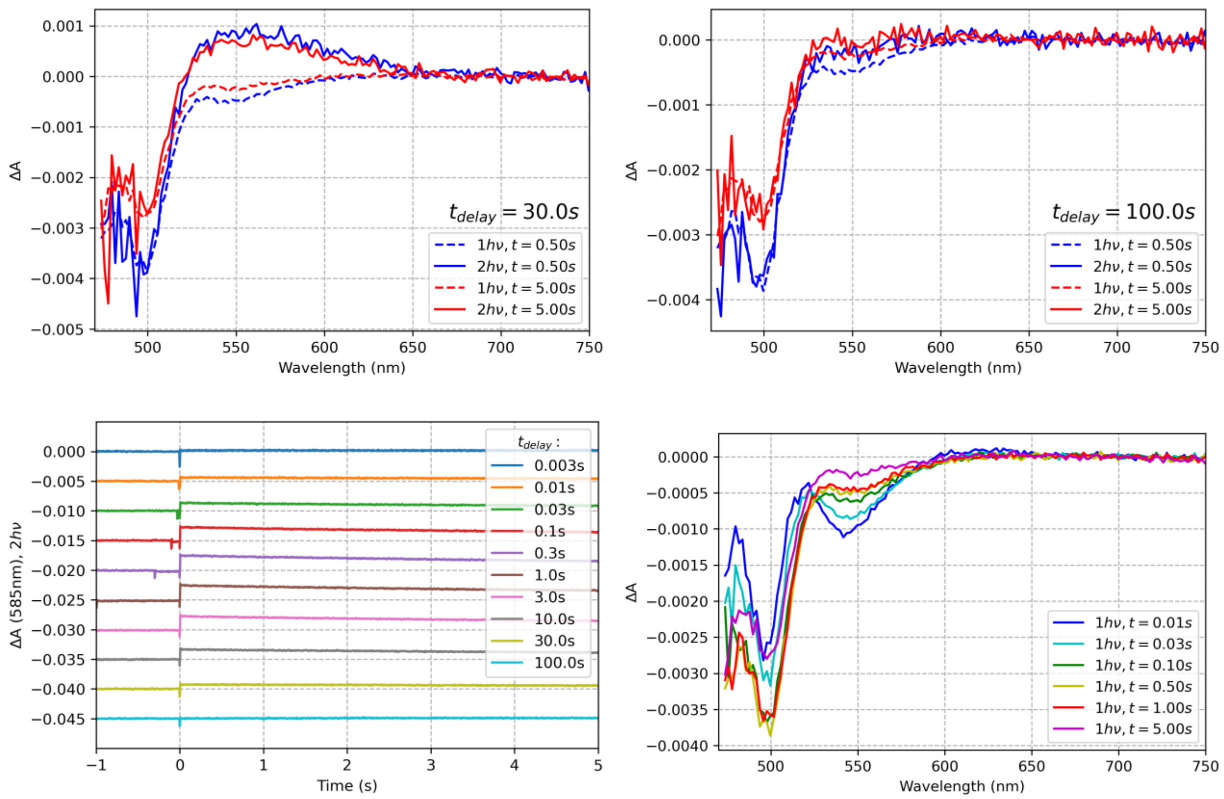


**Figure S26**. Two pulse experiment results obtained for the OCP mutant R27L from *Planktothrix* functionalized with ECN, with His-tag removed. Excitation at 512 nm, energy density about 50 mJ/cm$^2$.

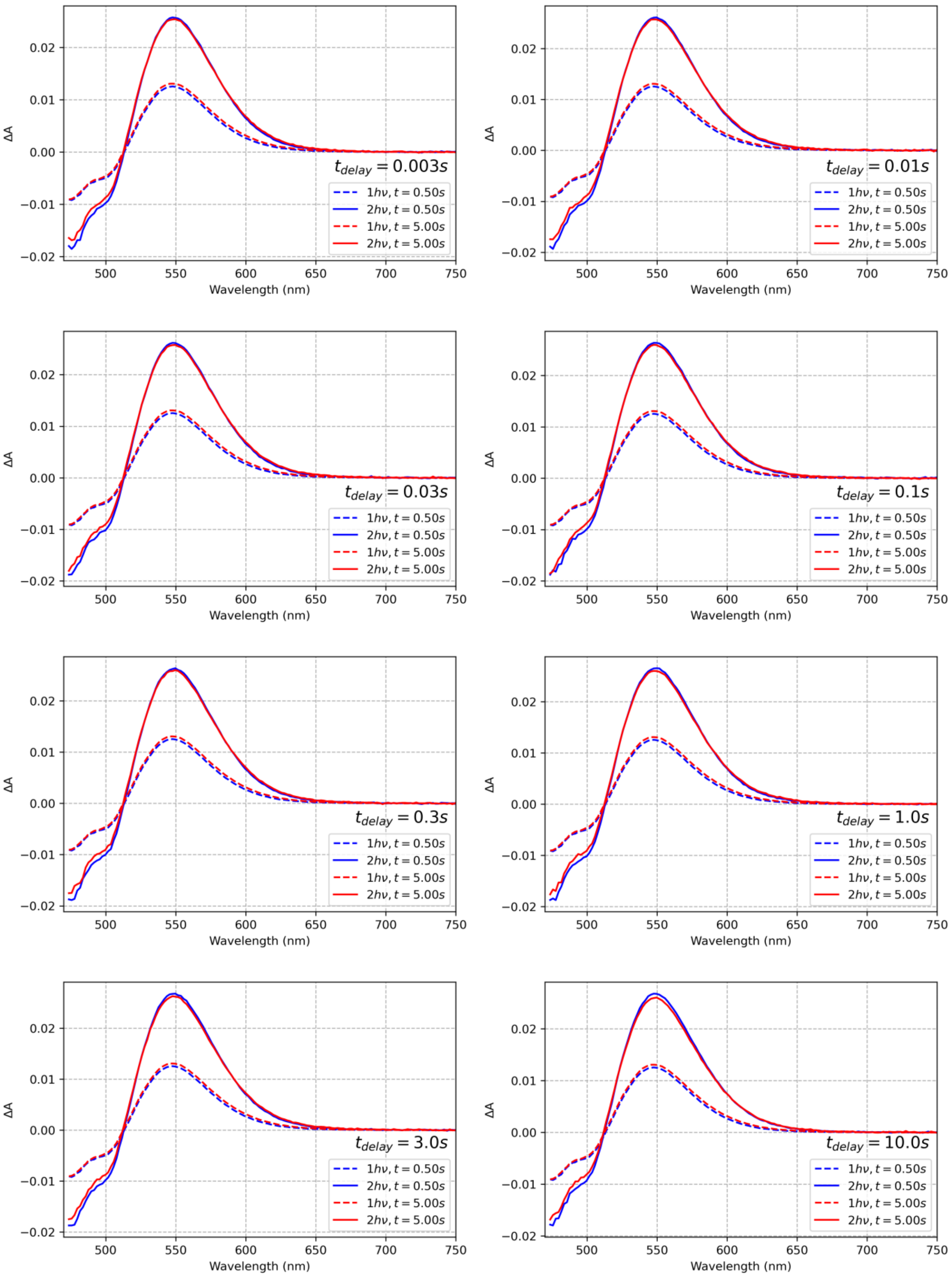

$t_{delay} = 0.003s$
$t_{delay} = 0.01s$
$t_{delay} = 0.03s$
$t_{delay} = 0.1s$
$t_{delay} = 0.3s$
$t_{delay} = 1.0s$
$t_{delay} = 3.0s$
$t_{delay} = 10.0s$
$1h\nu, t = 0.50s$
$2h\nu, t = 0.50s$
$1h\nu, t = 5.00s$
$2h\nu, t = 5.00s$
ΔA
Wavelength (nm)
0.02
0.01
0.00
−0.01
−0.02
500
550
600
650
700
750

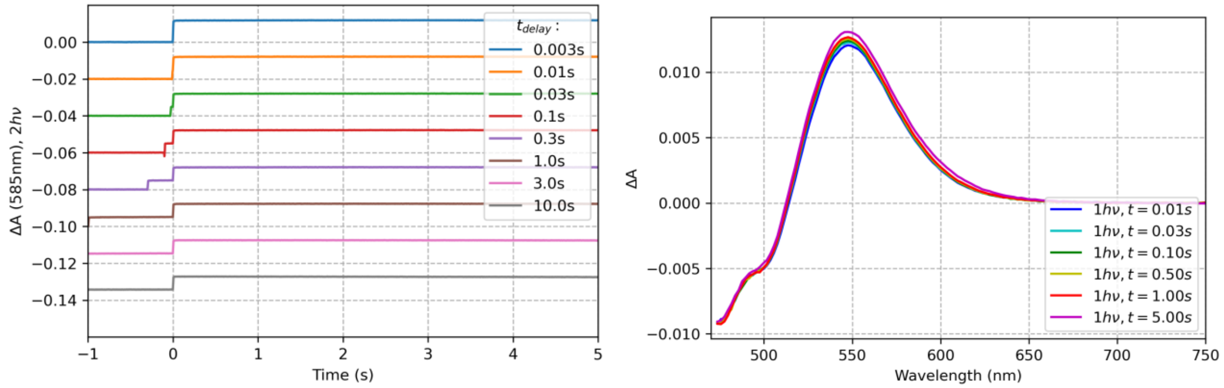


**Figure S27**. Two pulse experiment results obtained for the OCP mutant R27L from *Planktothrix* functionalized with CAN, with His-tag removed. Excitation at 512 nm, energy density about 50 mJ/cm$^2$.
